# Mutual information reveals a homonucleotide bias at 6 bp distance in promoters

**DOI:** 10.1101/786814

**Authors:** James L. Dannemiller

## Abstract

**Motivation:** Statistical dependencies between nucleotides at different positions within a DNA sequence have been used for several purposes including distinguishing coding from noncoding regions of a genome. Coding sequences show correlations within and between codon positions. This study asked whether such correlations between positions separated by short distances might also exist in noncoding DNA. To this end, positional nucleotide dependencies were examined in the promoter regions of four eukaryotic species: *Homo sapiens* (*Hs*), *Mus musculus* (*Mm*), *Drosophila melanogaster* (*Dm*), and *Saccharomyces cerevisiae* (*Sc*). The degree of dependency between pairwise positions across a set of aligned sequences was quantified by Mutual Information (MI) and visualized using a novel heatmap method.

**Results:** MI in promoter sequences aligned at their putative Transcription Start Site (TSS) generally decreased with increasing distance between two positions, but also showed a prominent increase at a distance of 6 base pairs (bp) (i.e., between nucleotides at *x* and *x* + 6 in a sequence) in the three multicellular species, but much less so in *Sc*. This dependency at a distance of 6 bp appears to reflect an *N*_1_ … *N*_7_ homonucleotide bias in promoters.

**Availability:** R code and data files available at github/dannemil/promoters.

**Supplementary information:** *Dannemiller-supplementary-documents*.*zip*

## Introduction

Statistical dependencies between bases separated by various distances have been documented in coding DNA (Datta & Asif, 2005). Coding DNA tends to show a 3–base pair (bp) periodicity because of the nonuniform use of the four nucleotides at different codon positions (Grosse, Herzel, Buldyrev, & Stanley, 2000; Yin & Yau, 2007). Indeed, this 3–bp periodicity can be used as a feature in attempting to distinguish exons from introns (Singh & Srivastava, 2020). Longer range correlations have also been reported both for prokaryotic and eukaryotic genomes (Bernaola-Galván, Carpena, Román-Roldán, & Oliver, 2002; Berryman, Allison, & Abbott, 2004; Holste, Grosse, & Herzel, 2001; W. Li & Kaneko, 1992).

There are several reasons to think that promoter sequences might show such dependencies, perhaps at different distances. Short Tandem Repeats (STRs) could contribute to correlations between positions that depend on the density, length and composition of the repeating unit. Promoter regions contain an abundance of STRs (Bolton et al., 2013; Gymrek, 2017). It has been shown that there are more pairs of identical single nucleotide polymorphisms (SNP) separated by 1, 2, 4, 6 and 8 bp than by 3, 5, 7 or 9 bp, and that this pattern is observed mainly within STRs (Madsen, Villesen, & Wiuf, 2007). There is an abundance of exceptionally long (≥ 6 repeats) STRs in the core promoter regions of human protein–coding genes, and many of these appear to be evolutionarily conserved (Ohadi, Mohammadparast, & Darvish, 2012). Microsatellites show a taxon–specific enrichment in eukaryotic genomes (Avvaru, Sharma, Verma, Mishra, & Sowpati, 2019). It is also known that transcription factor binding sites (short motifs) tend to cluster in tandem (homotypic regulatory clusters), and the repeating nature of these motifs would be expected to contribute to dependencies between positions over short ranges (Lifanov, Makeev, Nazina, & Papatsenko, 2003; Papatsenko et al., 2002).

Another reason for possible dependencies in short DNA sequences could be the 3–D helical structure of DNA. The curvature of DNA has been shown to be sequence–dependent (Fratini, Kopka, Drew, & Dickerson, 1982; Gabrielian & Pongor, 1996). Double helix structural parameters are related in very complicated ways to base pair steps geometrically through the sugar–phosphate backbone and energetically through steric base clashes (Gorin, Zhurkin, & Olson, 1995). Transcription factor binding depends both on DNA sequence as well as on structural helical features. It is significant that purine–pyrimidine steps open toward the major groove, and pyrimidine–purine steps toward the minor groove.

Molecular dynamics (MD) simulations indicate that longer range DNA allosteric characteristics can be affected by local sequence characteristics with poly d(AT) sequences leading to more bending while poly d(GC) sequences tend to lead to more twisting of the helix (Gu et al., 2015). The binding of a repressor to phage 434 depends on the recognition of a specific 14 bp sequence in six different operators, but the repressor only binds to the two 5 bp sequences on both ends of these motifs (Koudelka, Harrison, & Ptashne, 1987).

The central four base pairs are not directly contacted by the repressor, but they affect the binding affinity by sequence–dependent alterations of the bending of the helix (el Hassan & Calladine, 1996). While the five nucleotides on both ends of these binding sites are highly conserved, the four nucleotides bordering the center of symmetry (6, 7 and 8, 9) are more variable. This means that these central four nucleotides are in positions that could carry correlated variations to fine tune the binding affinity of the repressor. Thus, geometric and energetic consequences of DNA sequence might also favor certain combinations of nucleotides at specific distances within a promoter sequence.

The most straightforward way of exploring dependencies between nucleotides at different distances would be to quantify the dependency with a correlation. Computing standard correlations between positions separated by specific distances is not possible in the case of DNA sequences because the use of a Pearson’s r correlation coefficient requires that the random variables be numeric.^1^ The fact that the four DNA bases are nominal or symbolic with no intrinsic ordering or magnitude makes standard correlation metrics inapplicable (Wentian Li, 1990).

Alternatively, correlation methods that use nucleotide co–occurrence frequencies (joint nucleotide distributions) for assessing distance correlations in DNA sequences respect the categorical or symbolic nature of these sequences. An excess (or deficit) of the observed joint frequency of a particular combination (e.g. A/A) at two different positions relative to their expected frequency is a measure of the correlation of that combination at those two positions (Herzel, Weiss, & Trifonov, 1999). The same logic holds true for any of the other 15 possible joint nucleotide combinations.

Mutual Information (MI), an information–theoretic quantity related to joint entropy, uses the observed joint nucleotide frequencies between two positions across a set of aligned sequences to quantify correlations between positions. The MI statistic can be interpreted as the information carried by the nucleotides at one position about the corresponding nucleotides at another position. In this sense it is similar to a Pearson correlation that captures the degree of linear relation between two random variables. MI, however, also captures *any* form of dependency between random variables. Although, as mentioned above, MI is not strictly a correlation measure, it can be thought of as reflecting the degree to which the nucleotides at two different positions are ‘correlated’ in the intuitive sense of the word.

MI has been used previously in various ways to understand the structure of DNA and amino acid sequences (Berryman et al., 2004; Dunn, Wahl, & Gloor, 2008; Korotkov, Korotkova, & Kudryashov, 2003). It can be used to detect Short Tandem Repeats (Aktulga et al., 2007). The form of the MI vs. distance function between coding and noncoding regions is different within a species, but similar across species (Grosse et al., 2000). The MI statistic can be used as an alternative to linear regression to reveal both cis–and trans–eQTL (Huang & Cai, 2013) and to analyze gene regulatory networks (Margolin et al., 2006). MI has also been used to reveal phylogenetic relations based on mitochondrial (Kraskov, Stogbauer, Andrzejak, & Grassberger, 2005) and nuclear DNA sequences (Lichtenstein, Antoneli, & Briones, 2015). Additionally, MI has been used to reveal longer range correlations in DNA sequences (H. Herzel, Schmitt, & Ebeling, 1994). When MI is used as a measure of the correlation between positions in intronic sequences, it is found that it decays toward 0 very slowly, still being significantly above its null value in scrambled sequences as far as 800 bp away (W. Li & Kaneko, 1992).

While it has been shown that MI falls as the distance between two positions increases in non–coding DNA sequences, a close inspection of this relation in two studies showed an interesting reversal of this trend at a distance of 6 bp (Grosse et al., 2000; Holste et al., 2001). Magnification of Figures 1 in these studies shows a clear spike in the value of MI at a distance of 6 bp in human noncoding DNA and on Chromosome 22, respectively. The authors did not discuss nor explain this spike in MI at 6 bp because the focus of both studies was on longer–range correlations.

**Figure 0.1.**
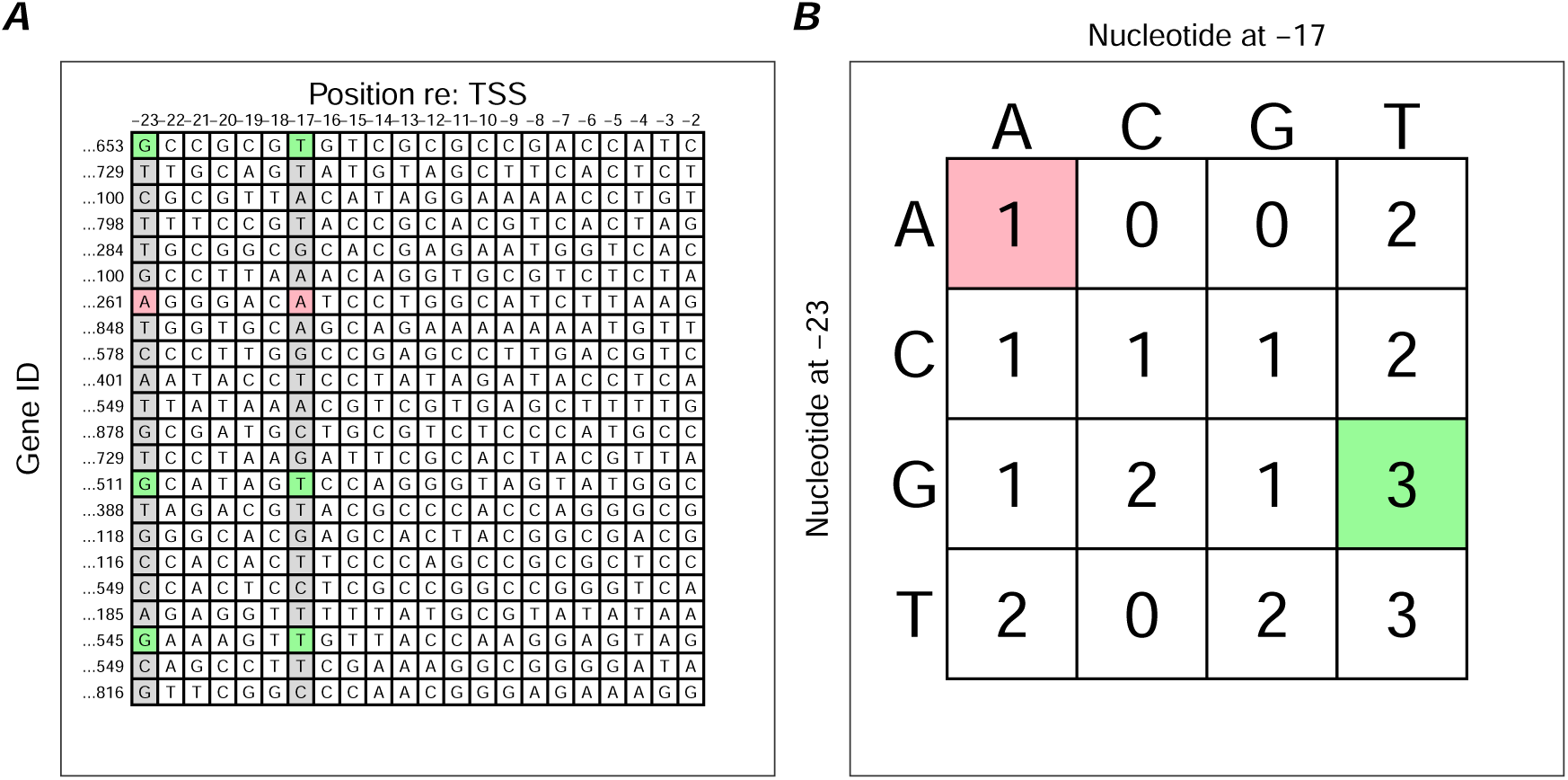
Illustration of how matrix MI is computed from aligned sequences. Panel A. Twenty-two artificial nucleotide sequences each covering the range from -23 bp to -2 bp with respect to their TSS. The number of sequences, 22, is small to make the example manageable in a figure. The pairwise nucleotide combinations from two positions (−23 and -17) are tallied and these frequencies are entered in the 4 x 4 table shown in panel B. For example, there are three instances of the combination G/T (shaded green) and one instance of the combination A/A (shaded red). Panel B. Table showing the frequencies of the 16 possible nucleotide combination across all 22 genes at the two positions, -23 and -17 bp re: TSS. The joint frequencies in the shaded cells were derived from the genes with the same shading color shown in Part A. Once this table is derived and its values are converted to proportions relative to the number of sequences, MI for the pairwise combination of these two positions is computed as shown in the Methods. The process is repeated for all pairwise combinations of the positions from -23 to -2 bp re: TSS.

The purpose of the present study was to examine more thoroughly the possibility of correlations between positions in promoter DNA sequences. Additionally, the function relating MI to distance was examined for evidence of the rise at 6 bp evident in the studies by Grosse et al. (2000) and Holste et al. (2001). Promoter sequences in four eukaryotic species, *Homo sapiens, Drosophila melanogaster, Saccharomyces cerevisiae*, and *Mus musculus*. were used to examine potential evolutionary trends in intra–promoter positional correlations. Coding Sequences (CDS) were also used as a control to ensure the ability of the MI measure to detect correlations in sequences known to possess dependencies within and between codons.

## Methods

### DNA promoter and CDS sequences

DNA promoter sequences for *Homo sapiens (Hs), Drosophila melanogaster* (Dm, dm6), *Saccharomyces cerevisiae (Sc)* and *Mus musculus (Mm)* were downloaded in FASTA format from the Eukaryotic Promoter Database (EPD) on October 20, 2019 with the download option set to *Select only the most representative promoter for a gene*. For each species promoter sequences comprising 1100 nucleotides extending from 1000 bp upstream to 99 bp downstream (TSS at + 1) were downloaded. After preprocessing to remove duplicate sequences and sequences with missing nucleotides the numbers of promoter sequences available for analysis by species were 16454, 13399, 5117 and 20203 for *Hs, Dm, Sc*, and *Mm*, respectively.

The coding sequences (CDS) for *Hs, Dm*, and *Mm* were downloaded using the UCSC Genome Browser. CDS for *Sc* were downloaded from the Saccharomyces Genome Database.

### Mutual Information

Mutual information (MI) is an information–theoretic measure of the amount of information about a random variable *Y* that is contained in the random variable *X* when *X* and *Y* are jointly distributed. Its theoretical lower bound is 0: a value that obtains only when *X* and *Y* are independent. Its upper bound depends on the number of symbols in the alphabet. In this case, with the four DNA nucleotides, the maximum MI value is 1.38.

Consider a set of aligned promoter nucleotide sequences to be a matrix with genes as rows and positions within the sequence as columns as shown in the left panel of Figure 0.1(a). To compute the pairwise MI value between positions -23 and -17 as shown in this figure, the joint nucleotide co–occurrence table on the right in Figure 0.1(a) is tallied. As shown by the pink shaded elements on the left corresponding to the joint occurrence of A/A at the two positions, there is only one sequence (ID … 261) with this combination, so a frequency of 1 is entered for the *A*_-23_*A*_-17_ combination in the table on the right. Similarly, for the combination *G*_-23_*T*_-17_ there are three sequences with that combination, so a 3 is entered in the appropriate cell. Once all of the joint occurrences for all of the sequences have been tallied, these joint frequencies are divided by N, the number of sequences, to convert them to empirical estimates of the nucleotide joint probability distribution at these two positions.

Given a 4 *×* 4 table of joint probabilities, let *p*_*i j*_ be the *joint* probability of row nucleotide *i* and column nucleotide *j*, and *p*_*i*._ and *p*_. *j*_ be the row and column marginal probabilities for that same combination of nucleotides. These marginal probabilities are just the simple, mononucleotide proportions at the two positions. MI between positions *l*_-23_ and *l*_-17_ is then defined as shown in Equation .1. MI will increase to the extent that the joint probability *p*_*i j*_ is greater than its expected value *p*_*i*._ *p*_. *j*_ derived from the assumption of independence.

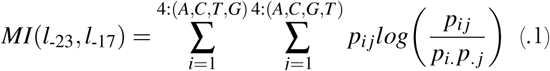

### Mutual Information vs. Distance between Positions

When MI is calculated for all possible pairs of positions in a range, an MI matrix like the one shown in Figure 0.2(b) results. The gray shaded cells show the major diagonal. All of the blue–shaded cell in this MI matrix indicate positions that are at a distance of 3 bp from each other: e.g., (−23, -20), (−22, -19), (−21, -18), etc.^2^ Similarly, all of the pink–shaded cells fall on the off–diagonal that represents a distance of 6 bp between two positions. To compute a function that reflects how MI changes with distance in this promoter region, the MI values along each of these off–diagonals were averaged along an arrow as shown in this figure. This figure is a simplified diagram of how various aspects of MI were extracted from the matrix MI matrix. It represents MI for a subsequence of 22 contiguous positions. In most of the analyses reported in this paper, the widths of these subsequences (windows) were at least 100 bp. Averaging all of the MI values along the blue line would give an estimate of MI for distance 3, and similarly averaging the MI values along the red line would give an estimate for distance 6. In the analyses reported below, mean MI was computed in this way for distances from 1 to 21 bp. All averages were based on 78 contiguous MI values along the appropriate off–diagonal.

**Figure 0.2.**
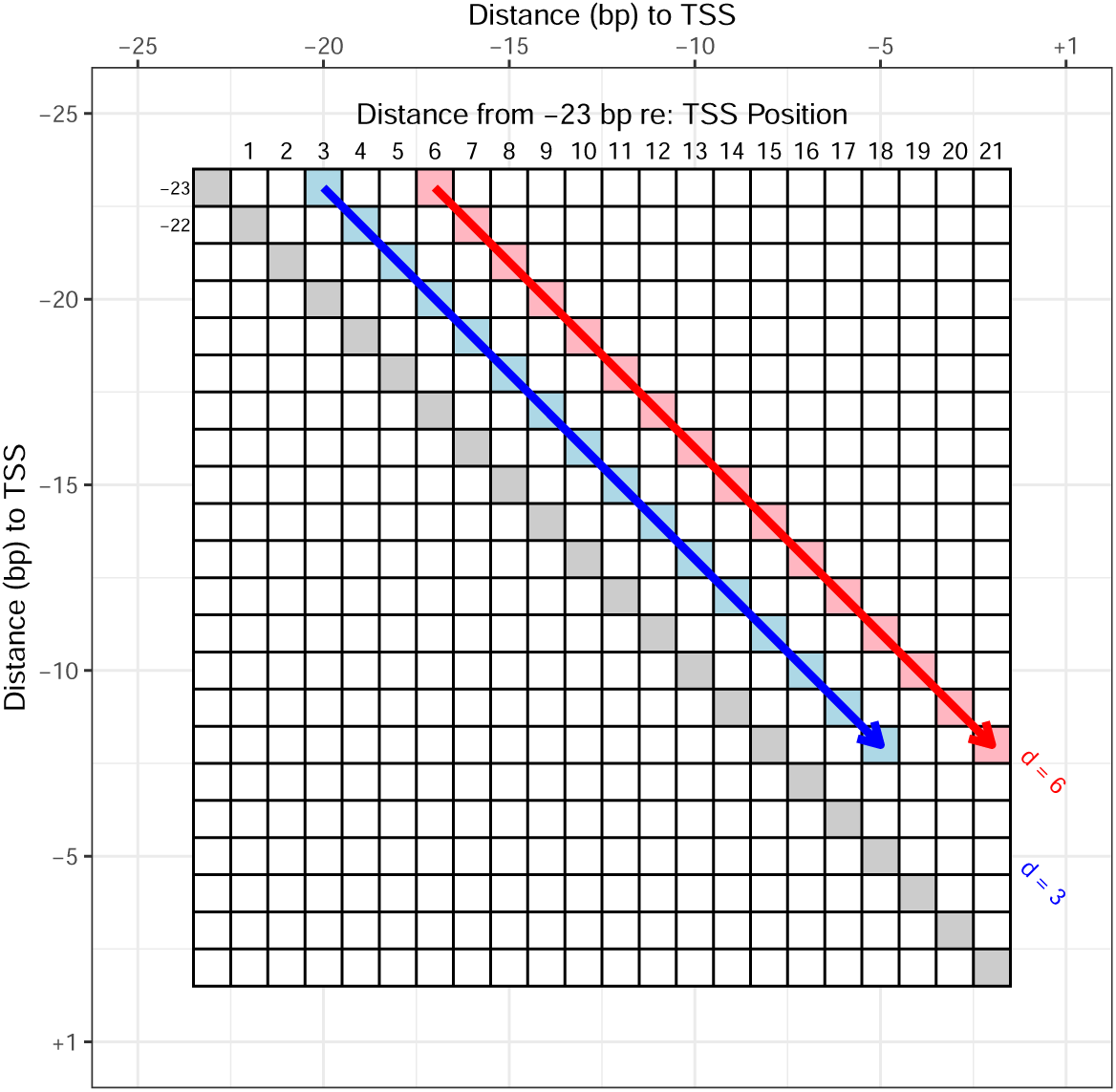
Illustration of how MI vs. Distance functions were estimated. This matrix holds 22 *×* 22 MI values (omitted for clarity); it is symmetric about the major diagonal (gray shaded cells). It covers the range from 2 bp to 23 bp upstream from the *TSS*_+1_. Each of the cells contains the pairwise MI value calculated from the two sequence positions at that cell’s coordinates. For example, the red shaded cell in the top row would hold the MI value calculated from the paired nucleotides at the sequence positions -23 and -17 bp re: TSS. The red shaded cell in the second row would hold the MI value for sequence positions -22 and -16 bp re: TSS, etc. MI vs. Distance functions were calculated by averaging the MI values along off–diagonals from upper left to lower right as shown by the red and blue arrows.

Finally, several authors have pointed out that MI values based on finite length sequences are subject both to random and to systematic error (bias). The magnitude of the sampling error scales as 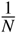 with *N* as the number of sequences from which the joint probability distribution was estimated (Holmes & Nemenman, 2019). Various methods have been proposed to correct for the bias in MI values (Holmes & Nemenman, 2019; Zhang & Zheng, 2015; Zhu, Bellanger, Shu, Yang, & Le Bouquin Jeannès, 2014). In contexts in which the absolute magnitude of MI is important, it would be necessary to correct for these finite–length effects. Because the focus of this work, however, was primarily on the relative values of MI as a function of the distance between two positions in a sequence, MI was left uncorrected. It should be noted that using the bias–corrected measure of MI in the R package *mpmi* did not materially affect the pattern of results obtained with the uncorrected MI values.

### MI covering the entire range from -1000 to +99

The promoter range from -1000 to +99 was divided for each species into 10 subranges (windows). These windows started at -950, -850, -750, etc, and were 100 bp in width.^3^ MI matrices were generated for each of these windows by calculating pairwise MI between all possible positions within the window. Each MI matrix was 100 *bp ×* 100 *bp*. These matrices were displayed as heatmaps with the color and saturation scaled to the log of the MI value. Log MI values were used as a compressive transformation because the presence of a relatively high MI value among many MI values close to the noise level resulted in visual detail being lost from the heatmap. Highly negative log MI values (i.e., MI values close to 0) appeared as white to pale yellow, and higher MI values shaded from orange through red.

### Understanding the Relations Between

#### Sequence Characteristics and Spatial Patterns in MI Heatmaps

Certain regular patterns appeared often in the heatmaps across regions and across species. To guide in the understanding of these patterns, artificial promoter sequences were produced that carried certain nonrandom relations between the nucleotides at different distance from each other. The characteristics of these artificial sequences were manipulated until a heatmap with a feature similar to the feature of interest in the observed heatmap could be produced.

Additionally, punctate (spatially delimited) features also appeared in the actual heatmaps, and these were simulated by generating artificial promoter sequences carrying short (e.g., 7 bp) motifs confined to one or to several contiguous positions in a fixed proportion of the sequences. Good examples of these features can be found in Supplementary Figures S10 and S20 showing the presence of the TATA box motif and the Initiator element in *Hs* and *Dm*, respectively.

A library of these simulated heatmaps was generated based on the observed regularities in the actual heatmaps. This library of patterns could be consulted to interpret the features present in the observed heatmaps. Relevant heatmaps from this library along with a description of how each of them was produced is shown in the Supplementary document *Understanding Mutual Information Heatmaps*.

## Results

To ensure that the method of computing MI as described in the Methods section was sensitive to the relations between nucleotides at specific distances from each other, an MI matrix was computed based on human Coding Sequences (CDS). The 47,118 transcript sequences comprised the initial 35 codons in each transcript. The start codon was eliminated from these sequences because the presence of the same trinucleotide in most of the sequences would produce a very strong punctate signal at that location in the heatmap making it more difficult to discern systematic, but somewhat weaker patterns elsewhere.

This Homo sapiens CDS heatmap is shown in Figure 0.3. The heatmap starts in the upper left corner with the first position of the second codon (position 4 with the start codon designated as +1). The positions within codons 2 to 5 are labeled in the upper left corner on the major diagonal. There is a striking regularity to this heatmap at distances that are multiples of 3 bp because of the fact that these sequences only contain strings of successive codons. Horizontally (and vertically), there is a signal (checks/pixels that are deeper shade of red) with a period of 3 bp throughout this range. Upon closer examination, this signal is present most strongly for position 3 of each codon (i.e., relative rows 3, 6, 9, etc.). In other words, nucleotides in successive codons are most strongly related in the third position. The third codon position is correlated more highly with the second position than it is with the first, and the first and second positions are themselves correlated. It is well known that there are correlations between the GC% content of the third codon position and that of the first two positions in both prokaryotic and eukaryotic species (Clay, Cacciò, Zoubak, Mouchiroud, & Bernardi, 1996; D’Onofrio & Bernardi, 1992). The method of using MI as a measure of correlation between nucleotides at different positions, and of plotting the MI matrix as a heatmap, behaved as would be predicted for sequences with a known periodicity of 3 bp.

**Figure 0.3.**
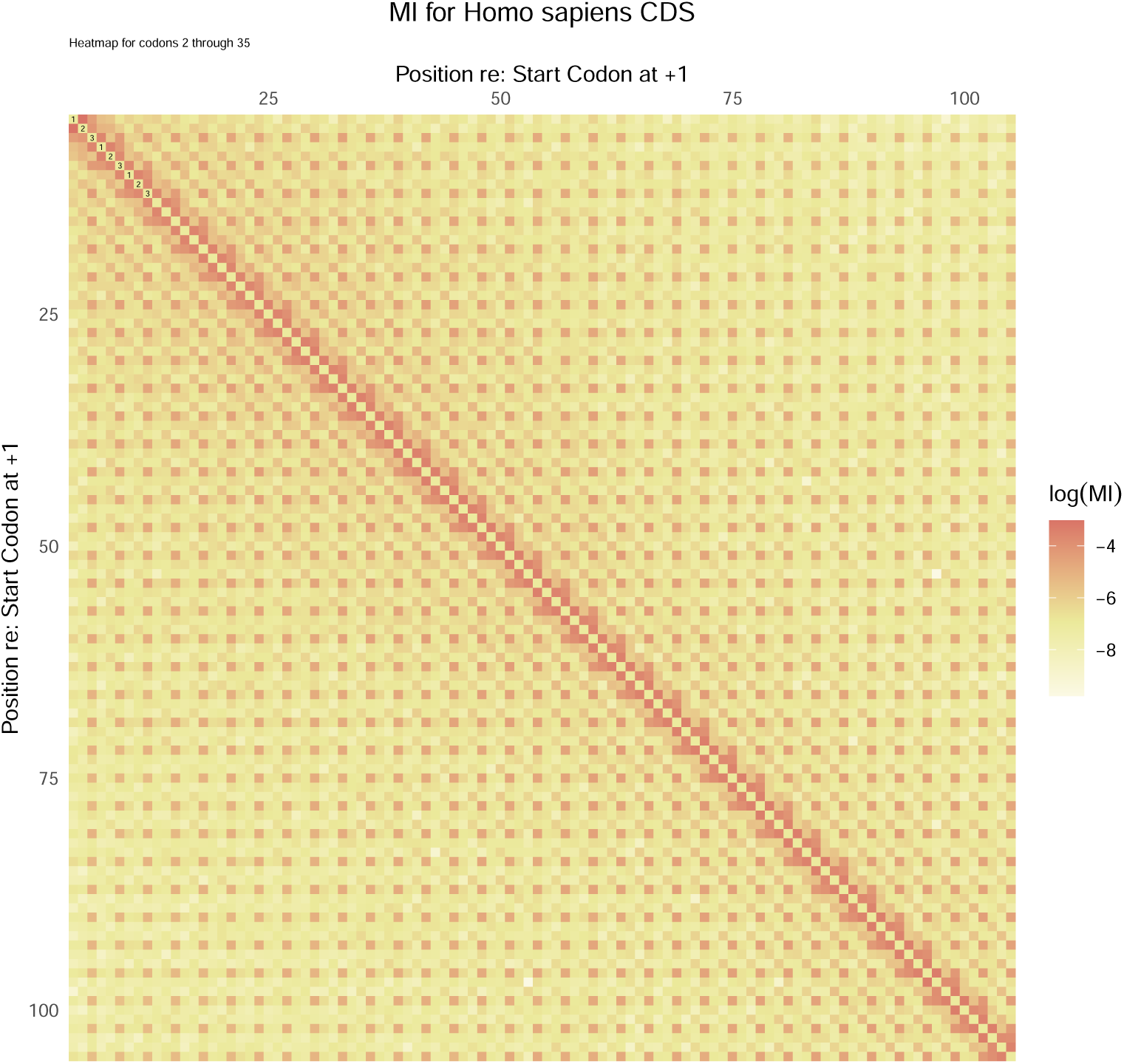
Heatmap of the 102 *×* 102 matrix of MI values generated from CDS sequences from human protein-coding genes.

### MI heatmaps from observed promoter sequences

The ten heatmaps for each species covering the range from -950 to +49 were examined for features indicative of higher than average MI values at fixed distances within the sequence. The complete set of these heatmaps is shown in *Supplementary Figures S1–S40*. MI vs. Distance curves were generated using the method shown in Figure 0.2. One feature that appeared in a large number of the heatmaps was an off–diagonal *line* at a distance of 6 bp. An example of this is shown in Figure 0.4.

**Figure 0.4.**
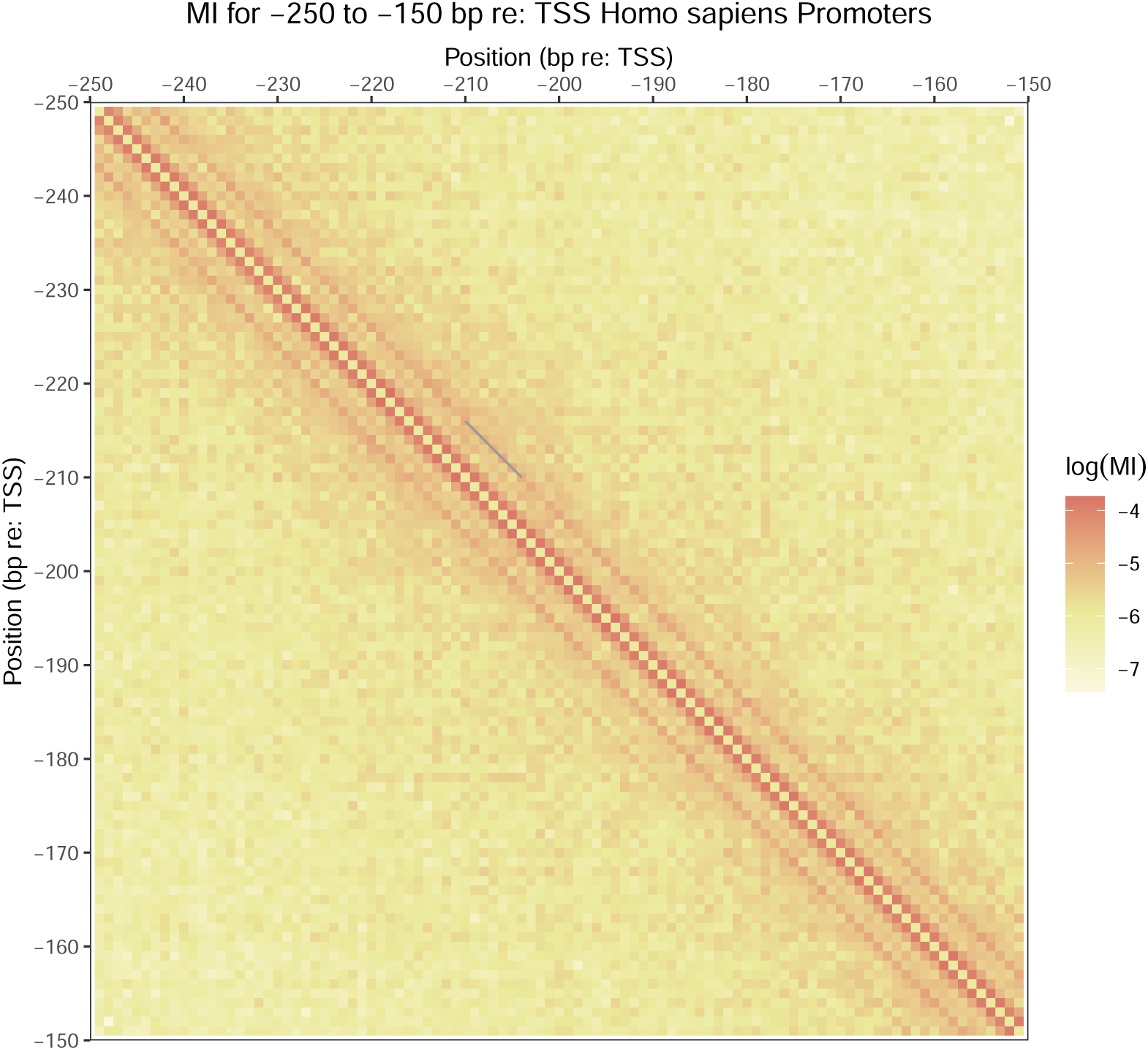
MI heatmap for Homo sapiens promoters in the range from -250 to -151 bp re: TSS. The thin gray line is positioned on the off-diagonal corresponding to a distance of 6 bp.

The thin gray line indicates positions that are related at a distance of 6 bp. By far, the strongest signal occurs along the major diagonal at a distance of 1 bp. These are neighboring positions. The signal at a distance of 6 bp implies that nucleotides at positions *l* and *l* + 6 (i.e., the first and seventh nucleotides) in a subsequence are more highly correlated than those at distances of 5 and 7 or longer distances. This Distance 6 Effect (D6E) was present most strongly in the heatmaps of the three multicellular species.

Calculations of average MI vs. Distances from 1 to 21 bp produced the plots shown in Figure 0.5 for all four species for the range from -250 to -151. Notice that as was evident in the heatmap shown in Figure 0.4 for Homo sapiens promoters, there is a signal at a distance of 6 bp that tends to be higher than at the immediately adjacent distances of 5 and 7 bp as well as at longer distances. This rise at a distance of 6 bp is less pronounced in *Sc* than in the three multicellular species. There is also some evidence for signals at distances of 3 and 12 bp for some of the species. This will be referred to as the Distance 6 Effect (D6E). This replicates the D6E that was present, but was not discussed in Grosse et al. (2000), Holste et al. (2001).

**Figure 0.5.**
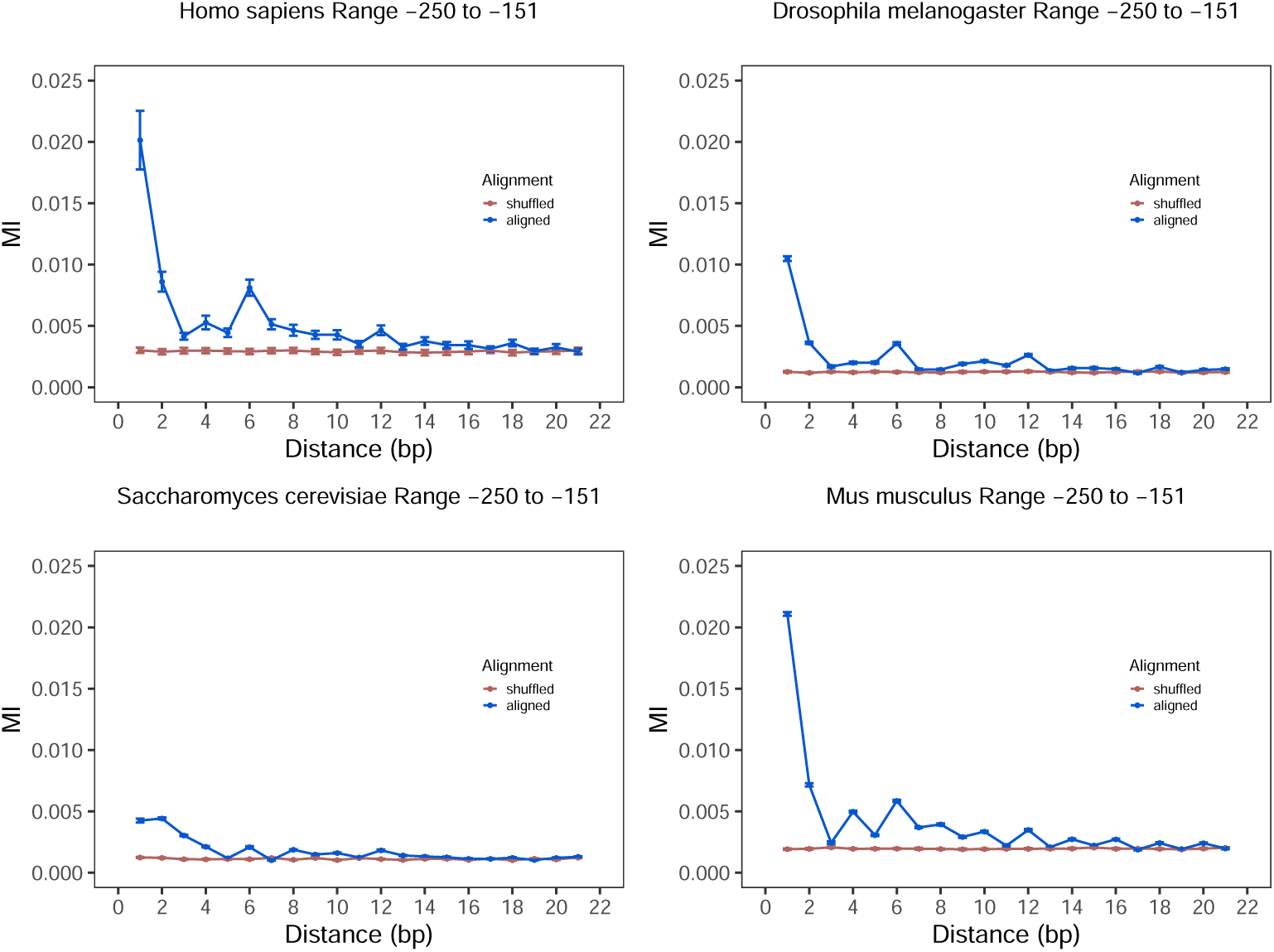
MI vs. Distance for all species over the range from -250 to -151. Errors bars where visible are *±* 1 SEM. Point means are based on averages of 78 contiguous MI values at the given distance. All plots were forced to have the same y-axis range for ease of comparison. The brown curve in each plot was generated by calculating MI on randomly shuffled sequences with the shuffling done within each original sequence.

To assess the range over which this D6E occurred, the MI values for all pairs of distance 6 positions from -950 to +49 (e.g., [-1000, -994]; [-999, -993] … [+43, +49], etc.) were compared to the MI matrix values obtained after randomly shuffling the 1100 bp sequences within genes and within species. The percentage of distance 6 MI values that exceeded the 95^th^ Percentile in each species–specific null distribution was tallied. The three multicellular species show more of these significant distance 6 MI values than are shown by *Sc* promoters: *Hs*: 71.7%, *Dm*: 90%, *Sc*: 41%, and *Mm*: 63%. The Wald-Wolfowitz Test for Randomness in run length (two–tailed) was used to determine whether these above–chance distance 6 MI values were distributed nonrandomly across the range from -950 to +49. In all species with the exception of *Sc*, the distribution of these runs was significantly different from chance meaning that runs of higher than average MI values along the distance 6 off–diagonal tended to be stronger in certain promoter regions. Parallel analyses along the distance 5 and 7 off–diagonals showed that a) the proportion of significant MI values was reduced from the distance 6 proportions for all species, and b) none of the runs tests rejected the null hypothesis of a random distribution of run lengths for distances 5 and 7 bp.

Is this D6E a local property within sequences or a more global property that depends on the precise alignment of the sequences? To discriminate between these possibilities, each sequence in the dataset for a given species was shifted randomly by an integer number of positions drawn from a uniform distribution with limits of -40 to + 40 bp (negative shifts move the sequence in the 3’ to 5’ direction). The nominal window from -250 to -151 was held fixed, and the MI matrix was computed based on the 100 bp that appeared within the window from each shifted sequence. The results for all four species are shown in *Supplementary Figures S41–S44*. In every case, the two curves are virtually indistinguishable. The function relating MI to the distance between positions does not depend on the precise alignment of the sequences. This is discussed further below.

Is there any way to determine what sequence characteristics are driving this D6E? One hypothesis is that this effect depends on a bias for the nucleotide at position *l* to repeat at position *l* + 6. Real sequences were searched for subsequences of 7 bp in which the first nucleotide repeated 6 bp downstream. This was done for all species using three pairs of positions (−250/-244, -200/-194, and -160/-154) in the range from -250 to -151. The 4 *×* 4 nucleotide co–occurrence table was generated for for each of the three pairs of positions for all four species. The standardized residuals from a chi–square test of independence on each of these tables were used to determine which of the 16 nucleotide combinations at the two positions significantly exceeded or fell below the expected frequency for that combination. For *Hs* and *Mm* 12 of 12 and for *Dm* 11 of 12 *N*_1_ … *N*_7_ homonucleotides were significantly enriched (above their expected frequencies). All of the other significant residuals represented combinations that were depleted relative to their expected frequencies. For *Sc*, in contrast, only 1 of 12 such *N*_1_ … *N*_7_ homonucleotides was significantly enriched. Thus, for the three multicellular species there is a bias in this promoter range from -250 to -151 to find the same nucleotides at positions 6 bp apart.

Finally, the generality of this *N*_1_ … *N*_7_ homonucleotide bias was tested by computing the frequencies of all of the like–nucleotide pairs at all distances from 1 to 21 bp within 100 bp windows across the promoter range from -950 to +49. The proportions of the like–nucleotides at each distance were aggregated and averaged across these 10 windows. The expected frequencies were derived by knowing the proportions of the four nucleotides within each 100 bp window and by assuming that the nucleotides across positions were independent. The observed minus expected difference was used as a measure of the homonucleotide bias.

The results are shown for all species in Figure 0.6. (please see also *Supplementary Figures S45–S48*). As shown by the red shading, all species show local maxima at distances of 6 and 12 bp. The rank order of these differences shows that the difference at a distance of 1 or 2 bp is always largest, followed by the difference at a distance of 6 bp in three of the four species. The other peaks appear to be more indiosyncratic to the species. It is interesting that the peaks in *Mm* appear at multiples of 2 bp. This pattern is similar to the results in Madsen et al. (2007) showing that the occurrence of identical SNPs is more common at distances of 1, 2, 4, 6 or 8 than at 3, 5, 7 or 9 bp in regions of the genome with periodic DNA. Indeed, the similarity between the *Mm* panel in Figure 0.6 and Figure 2B in Madsen et al. (2007) is remarkable, although the latter result held for human sequences and was not observed here.

**Figure 0.6.**
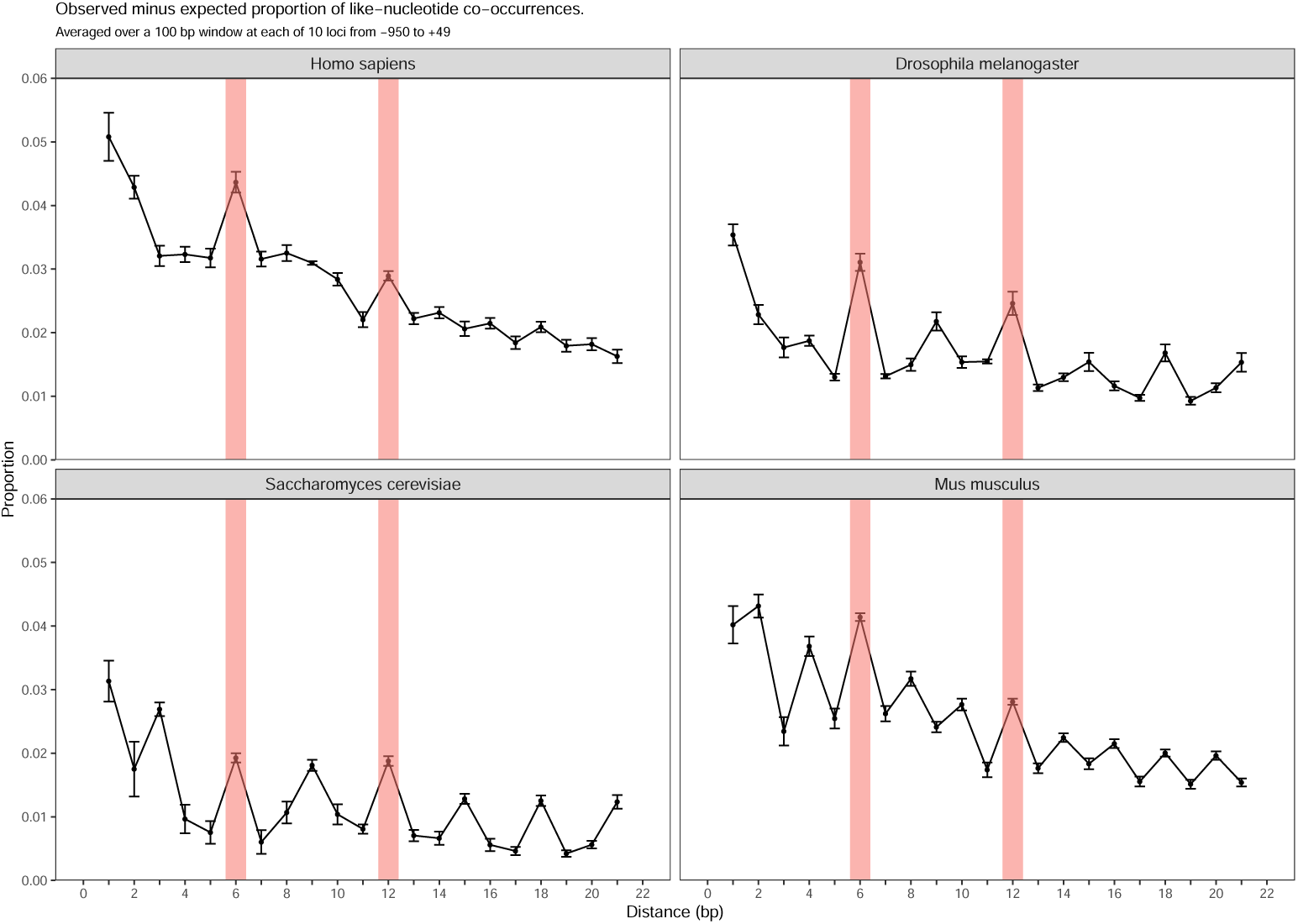
The observed minus expected proportions of like-nucleotides at distances from 1 to 21 bp averaged over the range from -950 to +49 bp re: TSS. Proportions were computed averaging over 10 different, 100 bp windows. Error bars are *±* 1 SEM. The red shaded rectangles show that all species show local maxima at distances of 6 and 12 bp, while other peaks at other distances are more idiosyncratic to the species.

## Discussion

Primarily in the multicellular species *Homo sapiens, Drosophila melanogaster* and *Mus musculus*, Mutual Information heatmaps revealed an ubiquitous regularity in promoter DNA. Across large portions of the range from -950 to +49 bp re: TSS there was a signal indicating an enhanced correlation between positions related by a distance of 6 bp: that is *l* and *l* + 6. This was not nearly as evident in *Saccharomyces cerevisiae promoters*. In several of the 100 bp subranges within this larger range, this D6E was present across the complete subrange. Runs tests showed that the distribution of runs of above chance MI values at a distance of 6 bp was not random; rather, these positions clustered. A good example of this occurred in *Dm*. The D6E was present in *Dm* promoters throughout the range from -150 to -51, but was absent in the core promoter region from -50 to +49. In contrast, the D6E was present in both of these ranges in *Hs* and *Mm* (Please *cf*. *Supplementary Figures S10 and S40 to S20*.^4^

Misaligning the sequences using random offsets with magnitudes in the range of *±*40 bp had no effect on the presence or strength of this D6E. This result, that is at first somewhat surprising, could have occurred for at least two reasons. First, it could be the case that every sequence shows the D6E across the entirety of the 100 bp that contribute to these curves. In other words, in sequence ID1, all of the nucleotides at distances of 6 bp (e.g., -250/-244, -249/-243, -248/-242, etc) as a set show this increased MI compared to other distances. The same could be true for other sequence IDs. Misaligning the sequences with respect to each other would make no difference because regardless of where a given sequence fell with respect to the start of the window at -250, all of the distance 6 pairs would still exhibit this property. A second possible explanation is that within sequences, some short segments (e.g., 14 to 28 bp) of the 100 bp window could show this D6E while other segments do not, and the phases of those segments with respect to the center of the window from -250 to -151 could be random and more or less uniform across the complete set of sequences. Once again, misaligning the sequences with respect to each other would still leave the distribution of these D6E segments within the window approximately the same.

There are several arguments against the first hypothesis above; namely, that the D6E reflects strong constraints on nucleotide choice at every pair of positions, *l* and *l* + 6, regardless of the absolute position of *l*. Even is this were true in ancestral promoters, single bp mutations, indels, and the presence of transposable elements is likely to have interrupted such a regular correlation at a distance of 6 bp. It is also not clear why such a constrain should be present in one region proximal to the TSS, but mostly absent in the neighboring core promoters region as app[ears to be the case in *Dm* (please *cf*. *S*upplementary Figures S19 and S20).

The second hypothesis is that the D6E appears in a heatmap more or less contiguously across a complete 100 bp window because a) it is a property of shorter segments of most promoter sequences, and b) the distribution of these segments across aligned sequences provides coverage across all positions within the 100 bp window. This argument hinges on how many sequences are necessary to produce a D6E. Analyses (not shown here) showed that the difference in the average MI value for distance 6 vs. the average MI at distances 5 and 7 stabilized to within 5% of its full value with as few as approximately 6400 sequences (40%) randomly chosen from the 16,454 Homo sapiens sequences. With fewer than 6400 sequences, the D6E was not consistently present. This result implies that there are short segments within sequences that have the *N*_*x*_ … *N*_*x*+6_ homonucleotide bias. As a set, the segments in these sequences cover the entire width of a given window.

These tandem segments are highly reminiscent of the homotypic clustering of short transcription factor binding sights known to exist in regulatory regions (Lifanov et al., 2003; Papatsenko et al., 2002).

It will take additional analyses beyond the scope of the current work to determine whether this D6E is tied to structural constraints imposed by the double helix. These analyses would be similar to those in Pedersen, Baldi, Chauvin, and Brunak (1998) that showed that several DNA structural parameters were tied to the density of certain trinucleotide repeats in a region proximal to the TSS. It is well known that the 3D conformation of local segments of the DNA molecule is affected by sequence characteristics. Some bp steps are preferred to others at specific positions because of the magnitudes and types changes in helical parameters necessary to accommodate those steps (Gorin et al., 1995; Gu et al., 2015).

Several important caveats are in order with respect to the inter–species comparisons in this work. First, the numbers of sequences used in the analyses differed across species. This has two potential effects. The precision with which MI values are estimated depends directly on the number of sequences used to compute pairwise MI between positions. This means that the noise in the MI measure will be greater for those species with fewer available sequences. In order of increasing precision, the species would be ranked Saccharomyces cerevisiae (5,117), Drosophila melanogaster (13,399), Homo sapiens (16,454), and Mus musculus (20,203). Second, for infinitely many sequences, the expected value of MI between two positions is 0 when the nucleotides at those positions are independent. For a finite number of sequences, even when two positions are independent, MI will be biased above 0, and the degree of bias depends inversely on the sample size (H. Herzel et al., 1994; Holmes & Nemenman, 2019; Zhang & Zheng, 2015; Zhu et al., 2014).

These effects are mitigated to some extent because the analyses are focused mostly on changes in promoter characteristics within species across different ranges of positions. Additionally, while the noise levels in the analyses will determine the extent to which certain features are visible in a heatmap, the signal–to–noise ratios for many of the features were generally well above the noise, making them easily visible. It is the case, however, that many of the indicators of the D6E were not present or present only weakly for *Sc*, and the *Sc* dataset had the fewest sequences, so conclusions regarding the *N*_*x*_ … *N*_*x*+6_ homonucleotide bias in *Sc* should be interpreted in light of this sample size difference.

Perhaps a more significant limitation arises because of differences in the specific sets of genes across species used in the analysis. This complicates inferences about species differences because the species are not being compared using a constant set of genes. To the extent that promoter architecture differs for different classes of genes (e.g., housekeeping vs. tissue–specific), this issue will complicate the interpretation of the results. It would be possible in principle to select a limited set of gene classes based on an ontology analysis, and within those classes to use only orthologous genes in the four species. Such an approach will be attempted in future analyses.

## Conclusion

Promoter sequences from -950 to +49 bp re: TSS showed an increase in Mutual Information for pairs of positions 6 bp apart that reversed the general trend for MI to fall with increasing distance. This Distance 6 Effect was most evident in *Hs, Dm*, and *Mm* promoter sequences, but not as evident in *Sc* promoters. The proximal cause of this effect appears to be an enhanced homonucleotide bias in promoter DNA for pairs of positions at a distance of 6 bp. The MI measure proved very helpful in revealing short–range correlations among positions in promoter sequences.

## Footnotes

1 It is possible to use a standard correlation coefficient by coding the DNA sequences into binary sequences using only pyrimidines and purines rather than all four nucleotides (Peng et al., 1993). This requires a trade–off against position resolution.

2 The first position is the row, and the second is the column (row, column).

3 Unless otherwise noted, distances are always with respect to the TSS, and the designation of ‘bp re: TSS’ is implied.

4 Supplementary Figure S20 shows a striking example in *Dm* of how MI can be used to reveal correlations between core promoter elements, in this case Inr and DPE.

## Notes

https://github.com/dannemil/promoters

### Competing Interest Statement

The authors have declared no competing interest.

### Summary of Updates

Additional evidence that nucleotides in many promoter sequences at arbitrary positions N and N + 6 (5’ to 3’) are more likely to be the same than would be expected by chance.

