## Supplementary material for "Mutual information reveals a homonucleotide bias at 6 bp distance in promoters": Supplementary-Document-Understanding-MI-Heatmaps.pdf

### 1 Supplementary Document: Understanding Mutual Information Heatmaps

When Mutual Information matrices are displayed as heatmaps, there is a monotonic mapping of the MI values in the matrix (here typically $100 \times 100$) to the color shades on the heatmap. The calculation of MI for a pair of positions is described in the Methods section. In the heatmaps used here, white represented MI = 0, pale yellow to yellow shades represented low MI values, and orange to red represented higher MI values. An MI matrix shows the mutual information between every position in a range and all of the other positions. The diagonal represents the mutual information between a position and itself, which is uninformative for the current purposes. Additionally, these self – MI values were typically an order of magnitude larger than the highest off – diagonal MI value. For this reason, the MI values on the diagonal were set to the mode of all of the non – diagonal MI values in the matrix.

Because the mapping from sequence characteristics to MI heatmap is not always intuitive, several of observed heatmap features presented in this work are discussed below. These features are:

1. *The signature of a motif:* Punctate (delimited) hotspots (deeper shades of red) located on the diagonal
2. *The signature of repetitive elements:* Diagonally – running 'lines' at various distances off the major diagonal

To understand how to interpret the conspicuous features in these heatmaps, a synthetic process was used. When an MI matrix is displayed as a heatmap, the process looks like this:

$$sequences_{real}\ (ATGCGC\ldots) \rightarrow \left|MI\ matrix_{real}\right| \Mapsto heatmap_{real}$$ [1]

Working backward from the spatial characteristics of the heatmap to sequence characteristics is complicated by the nature of the mapping described above. One way to approach this problem is to synthesize artificial sequences designed to contain a target feature or features observed in a heatmap. The process is *synthetic* in the sense that artificial sequences are constructed *de novo*. An MI matrix is computed based on these artificial sequences, and the corresponding heatmap is displayed.

The process in pseudocode was as follows:

$$1)\ simulate\ sequences_{artificial}\ (ATGCGC\ldots) \rightarrow \left|MI\ matrix_{artificial}\right| \Mapsto heatmap_{artificial}$$
$$2)\ heatmap_{artificial}\ \ cf.\ \ heatmap_{real}\ \ (compare\ the\ heatmaps)$$
$$3)\ if\ heatmap_{artificial}\ \ \neg\sim\ \ heatmap_{real}$$
$$go\ to\ 1$$
$$4)\ else\ if\ heatmap_{artificial}\ \ \sim\ \ heatmap_{real}$$
$$stop$$

To accomplish this, artificial sequences of width 200 bp were constructed.[2] Each attempt to produce a given target feature started with the same set (usually n = 16452) of these artificially – constructed, completely random sequences. Equal proportions were assumed for the four nucleotides so as not to bias the

[1] The $\Mapsto$ symbol indicates *maps to*

[2] The artificial heatmaps shown below are 100 bp wide. To avoid potential edge effects, sequences that were 200 bp wide were synthesized and trimmed to 100 bp after the various modifications had been applied.

results. This basic set of random sequences was then modified in various ways in an attempt to produce heatmaps containing features similar to those in the list above. This often required an iterative procedure in which some modification was introduced to the artificial sequences with the goal of reproducing a target feature in one of the real heatmaps. This was followed by rounds of adjusting the modification until the MI heatmap of the artificially – constructed sequences contained a feature that was judged to be the same as or similar to the target feature in the real heatmap. Often, it was necessary to try another approach to producing a given feature because the initial approach produced the feature, but it also produced correlated features not seen in the target heatmap. It is also the case that more than one way was occasionally found to reproduce a particular feature.

An artificial MI heatmap based on 16,454 random sequences each 100 bp wide is shown in Figure 1. The designations of position are completely arbitrary. It might seem odd to present a heatmap based on pairwise MI values calculated from random nucleotide sequences. This heatmap is presented so as to have a baseline with which to compare other artificial heatmaps intended to have targeted features. The nonuniformity in MI values is the result of sampling error and the finiteness of the sample size. With an infinite number of sequences, all of the MI values would be 0, and the heatmap would be entirely white. The nonuniformity also serves the purpose of giving a visual sense of the level of noise in the artificial heatmaps.

#### 1.1 Motifs: Punctate (delimited) hotspots (deeper shades of red) located on the diagonal

An example of this feature can be seen in the upper left quadrant of Figure 8. The 'smeared' square area indicates a hotspot — a punctate area with higher MI values. The fact that this feature is contained in a small area indicates that a small group of neighboring positions are all related to each other. This occurs when a short subsequence (e.g., a TATA box) is present at approximately the same place in a significant proportion of the sequences.

Figure 3 reproduces a similar feature. To produce this feature, the arbitrary subsequence TCAGACT was embedded within half of the completely random sequences with the initial T in the position -825 bp re: TSS. The red hotspot is seven bp wide as is the subsequence. The MI value is high because in half of all of the sequences a T at position -825 is followed by a C at position -824, and so forth for the other positions in the sequence. The point in the upper right corner of the hotspot indicates that the initial T at position -825 is followed at higher than chance frequencies 6 bp later by a T at position -819 . The feature is solid because the subsequence is embedded as an invariant whole in half of the sequences. Thus, punctate areas tend to indicate a short sequence occurring at the same position with a frequency that distinguishes it from chance levels. One important point to take away from this artificial heatmap is that it is not necessary for *all* of the sequences to contain a certain characteristic to see that characteristic appear as a distinguishable feature on the heatmap. To produce the heatmap shown in Figure 3 only half of the random sequences were modified. The proportion of sequences that carry a given characteristic will determine the saturation of the reddish color on the heatmap with smaller proportions leading to less saturated red features.

These punctate, square – like features on the diagonal are the signature of a motif in this kind of heatmap. For example, if a transcription factor binding site tends to occur at the same positions across a large set of sequences, then that motif will be evident in the heatmap as a reddish more or less square area sitting on the major diagonal. There are three criteria that must be met for such motifs to show up in a heatmap. First, the motif must be of limited length. By definition, this is true of binding site motifs. Second, a minimum proportion of the aligned sequences must carry the motif with limited variants. Third, the motif must appear at the same position in a set of aligned sequences with some minor tolerance for the exact position. These different ways of observing a punctate, on – diagonal feature will be addressed next.

Another example of such an artificial punctate feature is shown in Figure 4. The short 6 bp sequence TATAAA was embedded in 3 mutually exclusive sets of the random sequences (n = 1371 each: 8.33%) with the initial T appearing at positions -30, -29, and -28 in the three sets, respectively. The remaining sequences were completely random. The square area is 8 bp wide because across the set of sequences as a whole, the initial T appears at position -30 (in approximately one – twelfth of the sequences), and the final A appears at position -23 (in approximately one – twelfth of the sequences). Across the complete set of sequences, the TATA subsequence occurs in 25% of all of the sequences.

One important feature of this last heatmap is that if the same subsequence appears in different promoters at slightly offset positions, this produces a 'smeared' feature in the heatmap relative to the case shown in Figure 3. Had the subsequence TATAAA occurred in a large portion of the artificial random sequences in exactly the same six contiguous positions, the punctate feature would be exactly 6 bp square. Instead when it appears at positions offset by 1 or two bp, the aggregate 'square' region results from the superimposition of smaller square features. The length of a side in the aggregate square is determined by the union of the positions comprising the superimposed squares. That is, the subsequence appears in mutually exclusive sets of artificial promoters in three sets of positions: $\langle -30 \;\; to \;\; -25 \rangle$, $\langle -29 \;\; to \;\; -24 \rangle$ and $\langle -28 \;\; to \;\; -23 \rangle$. The union of these three sets is the set $\langle -30 \;\; to \;\; -23 \rangle$, so the 'square' feature in the heatmap will have a side that is $-23-(-30)+1=8$ bp. Note that the 'square' feature corresponding to the TATAAA box in the real heatmap in Figure 8, has the appearance of being a superimposition of several smaller squares offset from each other.

There are at least two reasons why essentially the same subsequence might appear in slightly shifted positions in different promoter sequences. First, the TSS used in the multiple sequence alignment might not be precisely correct for some of those sequences. If the TSS were off by one bp either way in a large fraction of the sequences, but the TATA – box subsequence always appeared at the same position relative to the TSS in all sequences, then this would lead to slight offsets in the punctate squares that appear in the heatmaps. Second, the TSS loci could be correct, but the TATA – box subsequence could start at slightly different positions across sequences. Either of these would result in a feature that appears as superimposed, shifted smaller punctate features.

Another way that a similar square – looking punctate feature can be produced is by embedding three short TATA-related sequences (TATAAA, TATATAA, TATAAAAG) of different lengths in mutually exclusive sets of sequences (n = 1371 each: 8.33%) with the initial T appearing at position -30 in all three sets. This is shown in Figure 5. The square area is 8 bp wide because all three subsequences start at the same position, and the longest of the subsequences is 8 bp. There are subtle differences between these 'square' punctate features shown in Figures 4 and 5. The square feature in the actual Homo sapiens heatmap in Figure 8 appears to be more similar to the feature in the artificial heatmap with the superimposed squares, Figure 4, than to the one with superimposed squares of different sizes, Figure 5. This can be seen in Figure 6 that presents an expanded (zoomed) version of the region near the TATA box in Figure 8. The constituent squares that are superimposed in this expanded heatmap are each 7 bp in length with their union resulting in a square region that is 8 bp in length. This could imply that the typical TATA – box in real human promoters is 7 bp length, but it starts primarily at two different positions offset from each other by 1 bp relative to the TSS across the sequences in which it occurs.

Finally, in all of the cases above, particular motifs with no variation in the nucleotides at the different positions within the motif were embedded. It is also possible to create this feature by embedding variant motifs in which the nucleotides at the positions within the motif are not fixed, but are allowed to vary in a constrained way. Figure 7 shows such a heatmap. A fixed – length motif was embedded in 80% of aligned sequences at exactly the same position. The embedded motif was actually a collection of similar motifs with nucleotide variants at some of the positions. In this case, the consensus sequence for the Inr motif,

YYANWYY, was embedded starting at position -2. IUPAC symbols for variants (e.g., Y = T/C) were used to create each motif variant with the permissible nucleotides at any position being equiprobable, and the nucleotides in all of the positions (with the exception of the third position, A) being chosen independently.

It might at first appear puzzling that the nucleotides at 6 of the seven positions within the Inr motif could be chosen independently, yet the reddish square feature indicates that positions within the motif have elevated MI values. The process of producing these variants is not completely random, but is instead constrained by the variability implied by the consensus sequence at each position. This essentially leaves positions still biased to contain certain nucleotides relative to completely random sequences. The low MI cross through the middle of the feature occurs because that position contained only A in all sequences. Because there is no variation at that position, it ca carry no information about any other position. The same result occurs when one of the ’variables’ in a Pearson r computation is in fact a constant; in that case, r = 0.

#### 1.2 Repetitive elements: Diagonally – running ’lines’ at various distances off the major diagonal

One of the most striking and ubiquitous features of these promoter MI heatmaps is the presence of extended diagonally – running ’lines.’ Several examples from observed heatmaps are shown in Figures 8 and 9. In the first of these figures, there is a prominent diagonally – running line at a distance of 6 bp in the upper left quadrant. In the lower right quadrant there are similar lines at distances of 6, 9, and 12 bp as well as fainter lines beyond a distance of 12 bp. In the second of these figures, there is a line at a distance of 6 bp that runs the entire length of this range from -250 to -150. The strength of the signal varies across this range, but it is evident nonetheless.

The small gray line segment in Figure 9 is meant to indicate the minor diagonal that holds all pairs of positions in this range at a distance of 6 bp from each other (e.g., -207 and -201). The fact that these off – diagonal lines extend for substantial distances, in some cases across the whole extent of the heatmap (100 bp), means that the nucleotides at any position, $x$, and those at the position 6 bp downstream, $x+6$, are related to each other in some nonrandom way in a nontrivial proportion of the full set of sequences. Notice that it is not necessary that every sequence exhibits this relation between the nucleotides at $x$ and $x+6$ for *every* position, $x$, within some range. Rather, as long as across the set of sequences in the aggregate covers the range, then the MI at all pairs of positions 6 bp apart will be increased above the background noise for the complete set of sequences.

To produce this ’line’ feature, each random sequence was modified by embedding 16 short, 7 bp subsequences in which the initial nucleotide, N, repeats in the $7^{th}$ position, and the intervening nucleotides (2, 3, 4, 5, 6) are free to vary. These subsequences can be represented as N.....N with the initial and final nucleotides being the same, and the intervening nucleotides free to vary. Within each random sequence the position of the initial N was chosen from the ranges 1 to 6, 7 to 12, 13 to 18, etc. until all 16 subsequences had been embedded[3]. Additionally, the identity of the repeated nucleotide, N, varied randomly across the 16 instances that were embedded within a given sequence. So, while positions 2 and 8 might contain a repeated A, positions 9 and 15 could contain a repeated G.

The result of this is shown in Figure 10. There is a clear signal at a distance of 6 bp across the entire range. Notice that the procedure described above produces sequences with the following characteristics. Starting at any position from 1 to 100, for example at position 11, approximately $1/6$ of the sequences, or 2742 sequences, would have the same nucleotides at positions 11 and 17, with one – quarter of those

[3]Embedding 16 of these short sequences will allow the nucleotide in the $7^{th}$ position to extend beyond the 100 bp width of the sequence. This was allowed to ensure that the signal at a distance of 6 bp that appeared in the heatmap extended across the full range. The MI matrix was trimmed to be $100 \times 100$ before plotting the heatmap.

2742 having A/A, one – quarter having C/C,etc. Another $^{1}/_{16}$ of the remaining 13,712 sequences, or, on average 857 sequences, would also, for example, have C/C nucleotides at positions 11 and 17 by chance. Thus, in approximately 3,599, or 21.9% of these sequences, the nucleotide in position 11 will repeat in position 17 (A/A, C/C, G/G or T/T). By induction, any pair of positions at a distance of 6 bp will show approximately 3,599 of the sequences with the nucleotide at position x repeated at position x + 6. Compare this to the number that would be expected to repeat a specific nucleotide by chance alone. This would be $^{4}/_{16}$, or 2,056 sequences, with this characteristic. The signal at 6 bp in Figure 10 results from the excess $(3,599 - 2,056 = 1,543)$ of like – nucleotide pairs in all of the positions related by a distance of 6 bp.

It is tempting to think that the signal at 6 bp in Figure 10 means that *within each sequence* the nucleotide at every position x carries significant information about the nucleotide 6 bp downstream. But this is not the case. In fact, in the artificial sequences described above, there would be on average $16 + {}^{(100-32-6)}/_{4} = 31.5$ pairs of 100 possible pairs of positions in which the nucleotides at a distance of 6 bp are the same. The reason that the 6 bp signal extends the complete length of the 100 bp range in the artificial heatmap is that *in the aggregate*, all pairs of positions from 1 to 100 are covered by a sufficient number of sequences with this characteristic.

An alternative way of producing the 6 bp signal is as follows. Making the nucleotide at position $x$ within a sequence dependent on the identity of the nucleotide in that sequence at position $x - 6$ can be accomplished by using a First Order Markov Process. The Markov process has 4 states at a given position within a sequence: A, C, G and T. Typically, a Markov processes is used to model time – varying systems. In this case, however, it can also be used to model the series of nucleotides along a promoter sequence with the distance step playing the role of the time step in a time – varying Markov process. It is only necessary to specify the initial state of the system (the nucleotide at the first position in the sequence) and the transition probability matrix to produce a sequence of nucleotides that exhibits a dependency between positions at any distance. In this case, the initial state of each sequence was taken to be the first nucleotide in a random sequence of nucleotides that was 100 bp wide (represented as the positions from -250 to -151).

For example, Table 1 shows the transition matrix that results (bottom matrix) from the weighted combination of two different transition matrices. The matrix on the top left indicates that when the nucleotides A, G or T appear at position $x$, the nucleotide that appears 6 bp downstream is random (25% probability each, A, C, G and T). However, when the nucleotide C appears at position $x$, it is always followed 6 bp downstream by the nucleotide G. The matrix on the top right is a completely random transition matrix. When these two matrices are weighted at 0.5 and 0.5, respectively, and summed, the resulting transition matrix on the bottom results.

This matrix was used to generate 16,454 artificial sequences, each of width 100 bp. The $100 \times 100$ pairwise MI matrix was calculated between all of the positions, and the resulting heatmap is shown in Figure 11. Without showing the results here, the same procedure can be used to produce a signal at any distance (e.g., 1 or 2 bp) that runs the length of the heatmap. There is clearly a signal at a distance of 6 bp. Notice, however, that there is also a weak signal at a distance of 12 bp. In a first order Markov process, the future state of the system depends only on the current state and the transition matrix. Translating this to the case of sequences, this would imply that the nucleotide at position x should depend only on the nucleotide at position x - 6 and the transition probability matrix. Why then does this heatmap show a dependency between a nucleotide at position x and the nucleotide 12 bp upstream at position x - 12?

First order Markov processes with certain characteristics (those characteristics were deliberately chosen to apply to the process used here) possess a unique stationary distribution. The stationary distribution is the probability distribution over the possible states of the system that the system exhibits if it is run long enough. The phrase “run long enough” in the current context refers to positions that are far enough downstream from the current position. Once the system reaches its stationary distribution, all future probability distributions over the possible states will remain unchanged, and the future states of the system will be completely independent of the initial state. This does not mean, however, that a first order Markov process will reach this stationary distribution after one step. In the current context, the first 'step' from position 1 in each sequence will determine the nucleotide at position 7. The second step in the Markov process (second transition) will determine the nucleotide at the position 12 bp downstream from position 1 in each sequence (i.e., position 13). The third, fourth and fifth steps would then determine the nucleotides at positions 19, 25, and 31. The question is: Why are the MI values at a distance of 12 bp throughout the length of the heatmap slightly above the noise?

To answer this question, the Markov process described by the transition matrix in Table 1 was run for 7 steps assuming an equiprobable distribution of nucleotides across sequences in position 1. If a Markov process is run for n steps, the state of the system (distribution of the four nucleotide classes) will be given as $s_1P^n$ where $s_1$ is the initial state, and P is the transition matrix. In this case $s_1$ is the nucleotide in position 1 of each sequence, and the process was used to produce the nucleotides at positions 7, 13, 19, 25, 31, 37 and 43. After each 6 bp step, the pairwise MI was calculated between position 1 and position n ($n = \langle 7, 13, 19, 25, 31, 37, 43 \rangle$) and recorded. The value of MI vs. distance (6, 12, 18 ...) is shown in Figure 12. Log MI is plotted on the y – axis vs. distance on the x – axis. The red line is the mean of all of the off – diagonal log MI values and serves as the noise level in this plot. It is clear that signals at distances of 6 and 12 bp should be detectable in a heatmap like the one shown in Figure 11. Although the MI value is still not 0 at a distance of 18 bp (3 transitions steps), it is low enough as to fall below the background noise.

The number of such off – diagonal lines at multiples of the initial distance that appear above the noise depends on the percentage of such sequences that contain nucleotides generated by this First Order Markov process as well as on how strongly the transition probabilities deviate from equality (25%). The effect of varying the proportion of sequences generated using a First Order Markov process can be seen in Figure 13. In the heatmap on the left, 10% of the signals were generated using a First Order Markov process in which the nucleotide at position x depended on the nucleotide at position x - 6. 90% of the sequences were completely random. In the heatmap on the right 35% of the signals were generated using a First Order Markov process, and 65% of the sequences were completely random. The presence of off – diagonal lines at multiples of 6 bp was explained in the preceding paragraph.

It is notable that even with only 10% of the sequences following a First Order Markov process, the diagonally – running line is clearly visible above the noise, and the line at a distance of 12 bp is just discernible above the noise. The transition probability matrices used to generate the sequences following a First Order Markov process are shown in Table 2. The matrix on the left was applied to 10% of the sequences, and the matrix on the right was applied to 90% of the sequences. Alternatively, the matrix on the bottom in this table that is the weighted (10%/90%) sum of these two matrices was applied to 100% of the sequences. It is remarkable that in the matrix on the bottom, 12 of the 16 transition probabilities deviate by only 1% from random (equality: 25%), while 4 of the 16 deviate in the opposite direction by only 3%, yet a signal at a distance of 6 bp is still clearly visible in the heatmap. It is not the case that for 10% of the sequences (n = 1645 sequences), there was an invariant mapping from the nucleotide at position x to the nucleotide 6 bp downstream at position x + 6. This, perhaps, would make sense as an explanation for the visibility of the signal in the heatmap on the left in Figure 13. That is if an A at position x were always followed by a G at position x + 6, a C followed by a T, a G followed by an A, and a T followed by a C, this would mean that in 1645 of the 16454 sequences, regardless of where position x fell in the 100 bp range, this mapping would always apply. But this is not the case. Instead, the mapping is simply slightly biased probabilistically toward one or another of the four nucleotides at position x + 6 by the identity of the nucleotide at position x.

As an approximate way of determining the parameters of these sequences generated using a First Order Markov process that lead to heatmaps with signals that are just above the noise visually, the percentage of signals following the First Order Markov process was held constant at 10% and the nonrandom transition matrix shown on the upper left in Table 2 was varied. After trial – and – error, it was determined that setting the four most deviant cells in this matrix to 0.35 (where 0.25 is equal probability) with the other 12 cells set to $^{(1-0.35)}/_{3} = 0.2167$ produced the heatmap shown in Figure 14. The signal at a distance of 6 bp is just visible.

Finally, statistically, with no bias present at all in the First Order Markov process, the variance in the MI values would reflect multinomial sampling in a $4 \times 4$ contingency table with n = 16454 and all of the cells having equal probability: $p = {}^{1}/_{16}$. The standard deviation of this null MI matrix was approximately 0.00014 with a mean MI of 0.00028. Resampling was used to compose a random $100 \times 100$ symmetric MI matrix, but the MI values corresponding to a distance of 6 bp were sampled from a normal distribution with the same standard deviation and a mean that was 1.5SD above the mean of the null MI distribution. The result is shown in Figure 15. The signal at a distance of 6 bp is weakly visible in this heatmap.

#### 1.3 Summary

The synthetic method proposed here to understand the relation between spatial features of the MI matrix heatmap and corresponding sequence characteristics proved useful. As noted above, this method does not definitively determine the cause of particular heatmap features in heatmaps produced from real promoter sequences. In some case, it is fairly clear how a particular feature is related to sequence characteristics. Small square – like regions reflect short subsequences occurring at a limited set of positions. The presence of diagonally running lines in a heatmap, however, does not lead back unambiguously to the sequence characteristics that produced those lines. This is primarily because that feature can be created using mutually exclusive subsets of the sequences with slightly different sequence characteristics *between* the subsets, or by allowing structure *within* sequences at multiple scales. The synthetic method has the advantage of also revealing unintended heatmap features that are correlated with an intended feature. The process is not unlike trying to understand the mapping between the frequency domain and the time domain when a Fourier transform is used to represent a time – varying signal.

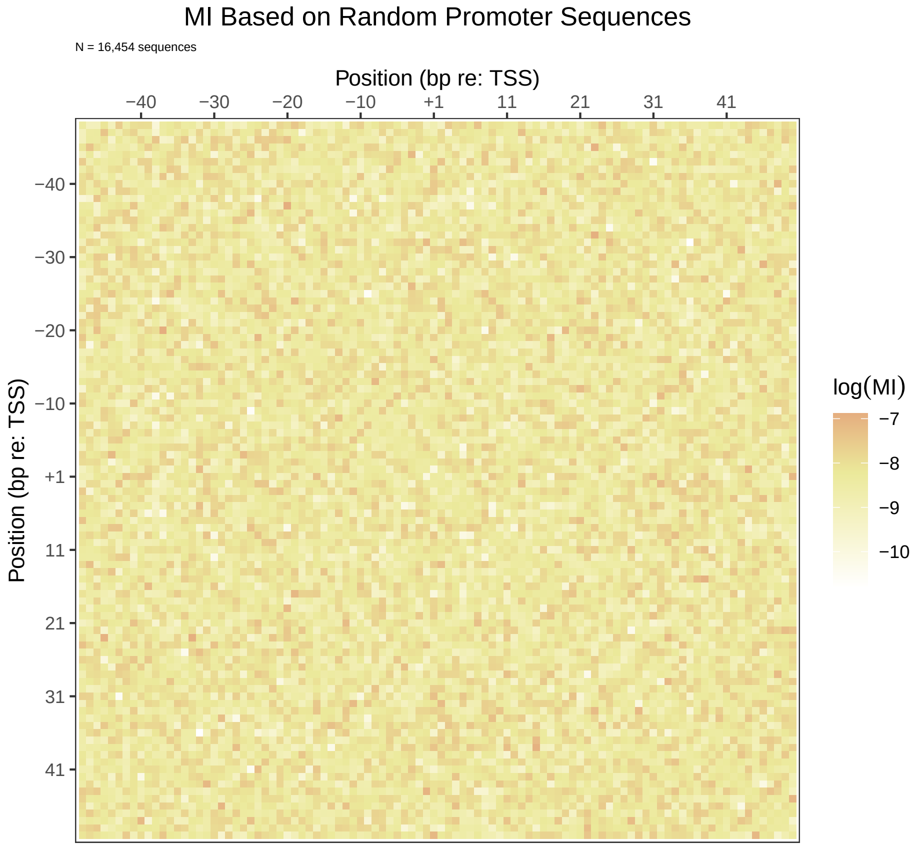


Figure 1: Artificial MI heatmap based on 16,454 random sequences each 100 bp wide. The designations of position are completely arbitrary. This heatmap is presented so as to have a baseline with which to compare other artificial heatmaps with target features. The nonuniformity in MI values is the result of sampling error and the finiteness of the sample size. With an infinite number of sequences, all of the MI values would be 0, and the heatmap would be entirely white.

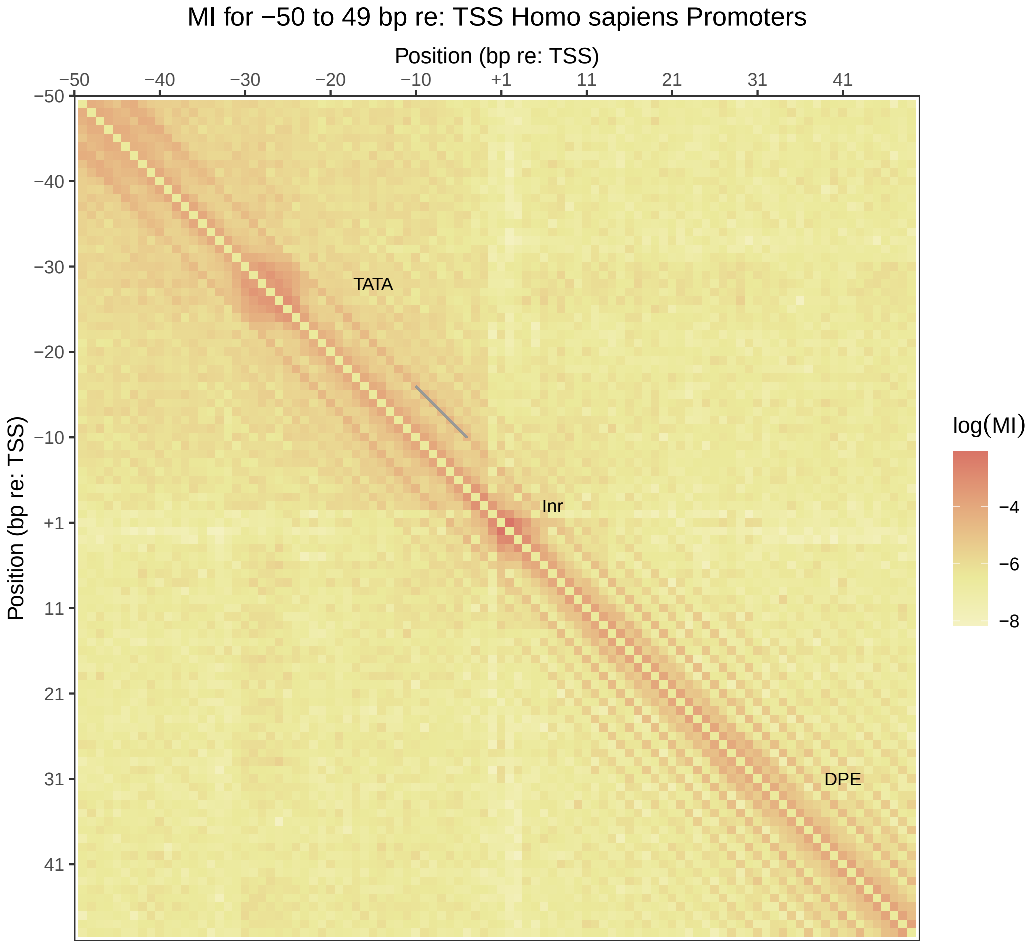


Figure 2: An example of an observed heatmap with a punctate feature in the upper left quadrant and a smaller one near the center of the heatmap. The square region contained within the blue dotted lines is expanded and presented in Figure 6. This outlined region is larger than the reddish punctate feature produced by presumptive TATA-box subsequences.

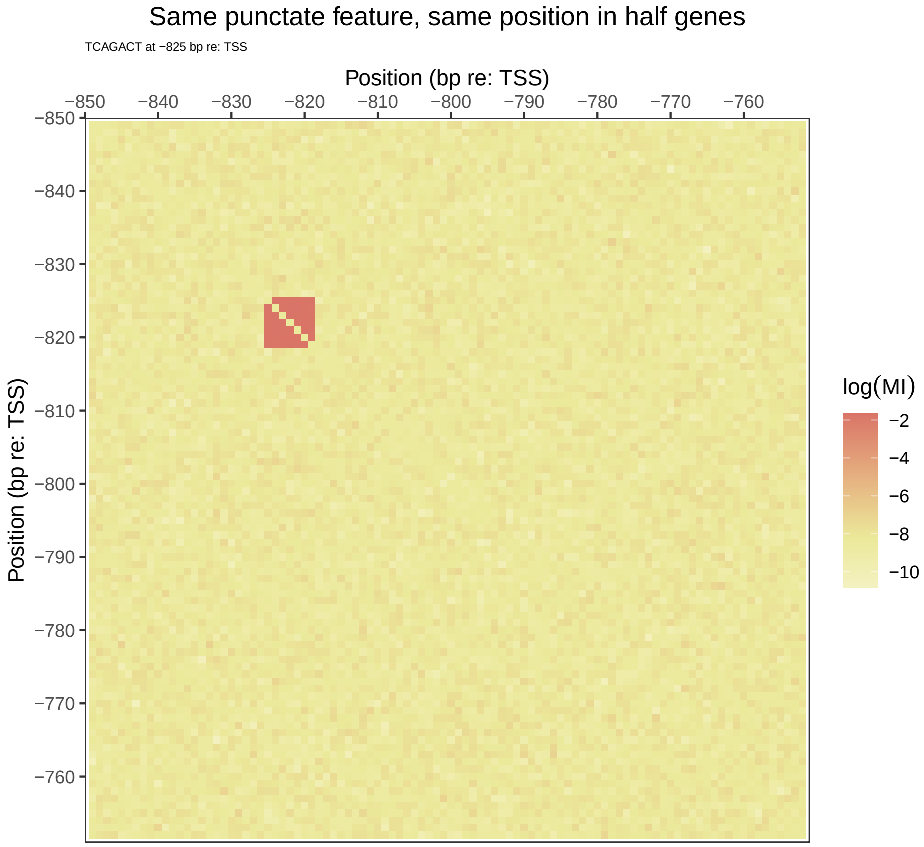


Figure 3: An artificial heatmap with a punctate feature. In this case the subsequence TCAGACT appears starting at position -825 bp re: TSS.

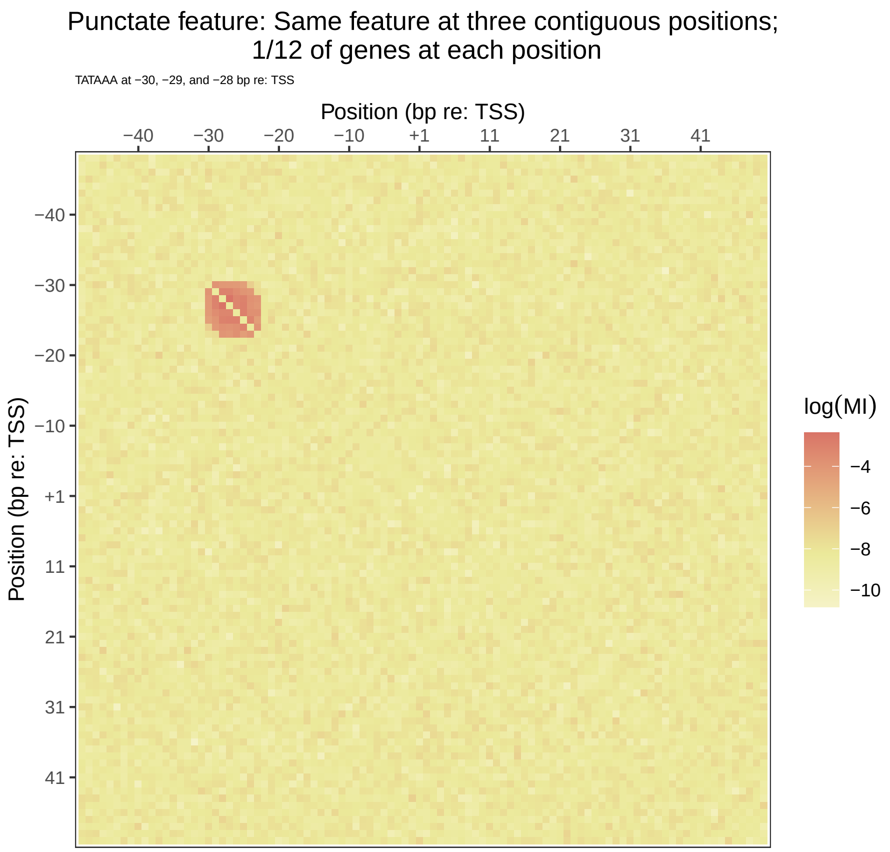


Figure 4: An artificial heatmap with a punctate feature. The short 6 bp sequence TATAAA was embedded in mutually exclusive sets of sequences (n = 1371 each: 8.33%) with the initial T appearing at positions -30, -29, and -28 in the three sets, respectively. The 'square' area is created by the superimposition of the TATAAA sequence shifted by 1 bp in each set. The full extent of the feature is 8 bp because across the set of sequences as a whole, the initial T appears at position -30 (in approximately one-twelfth of the sequences), and the final A appears at position -23 (in approximately one-twelfth of the sequences)
.

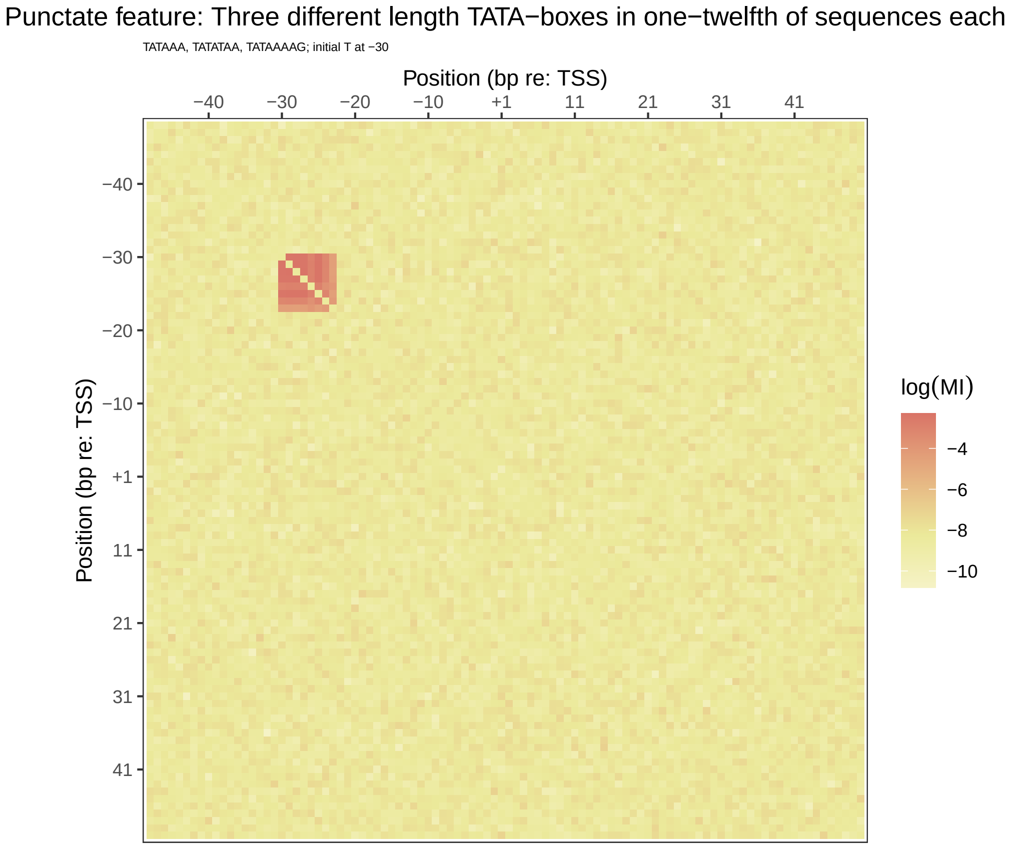


Figure 5: An artificial heatmap with a punctate feature. Three short TATA-related sequences (TATAAA, TATATAA, TATAAAAG) were embedded in mutually exclusive sets of sequences (n = 1371 each: 8.33%) with the initial T appearing at position -30 in al three sets. The square area is 8 bp wide because all three subsequences start at the same position, and the longest of the subsequences is 8 bp.

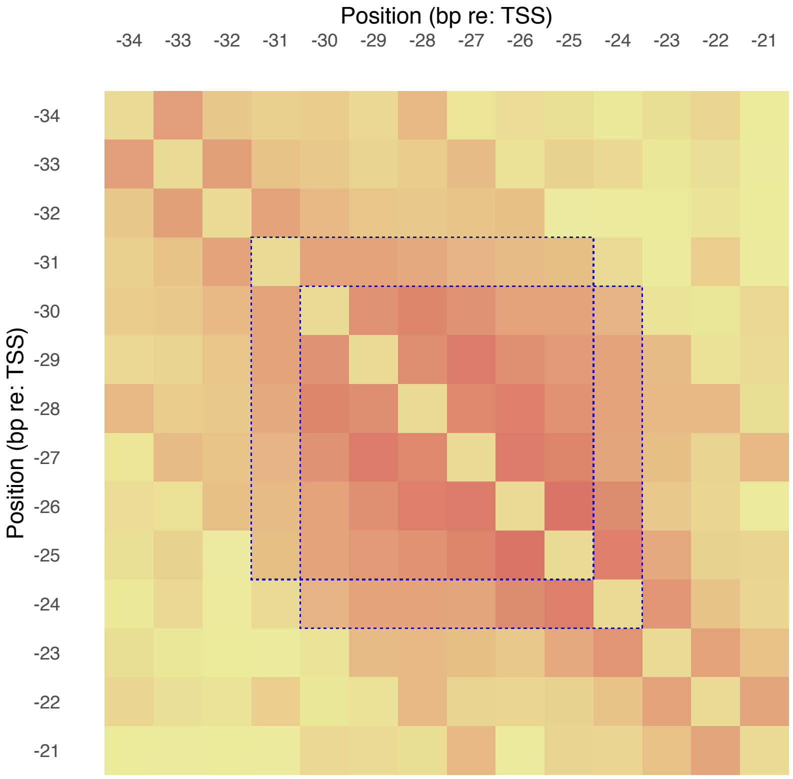


Figure 6: The region in Figure 8 that is outlined with blue dotted lines is expanded in this heatmap. Two superimposed square regions shifted by 1 bp are outlined here by way of comparison with the artificial heatmap shown in Figure 4.

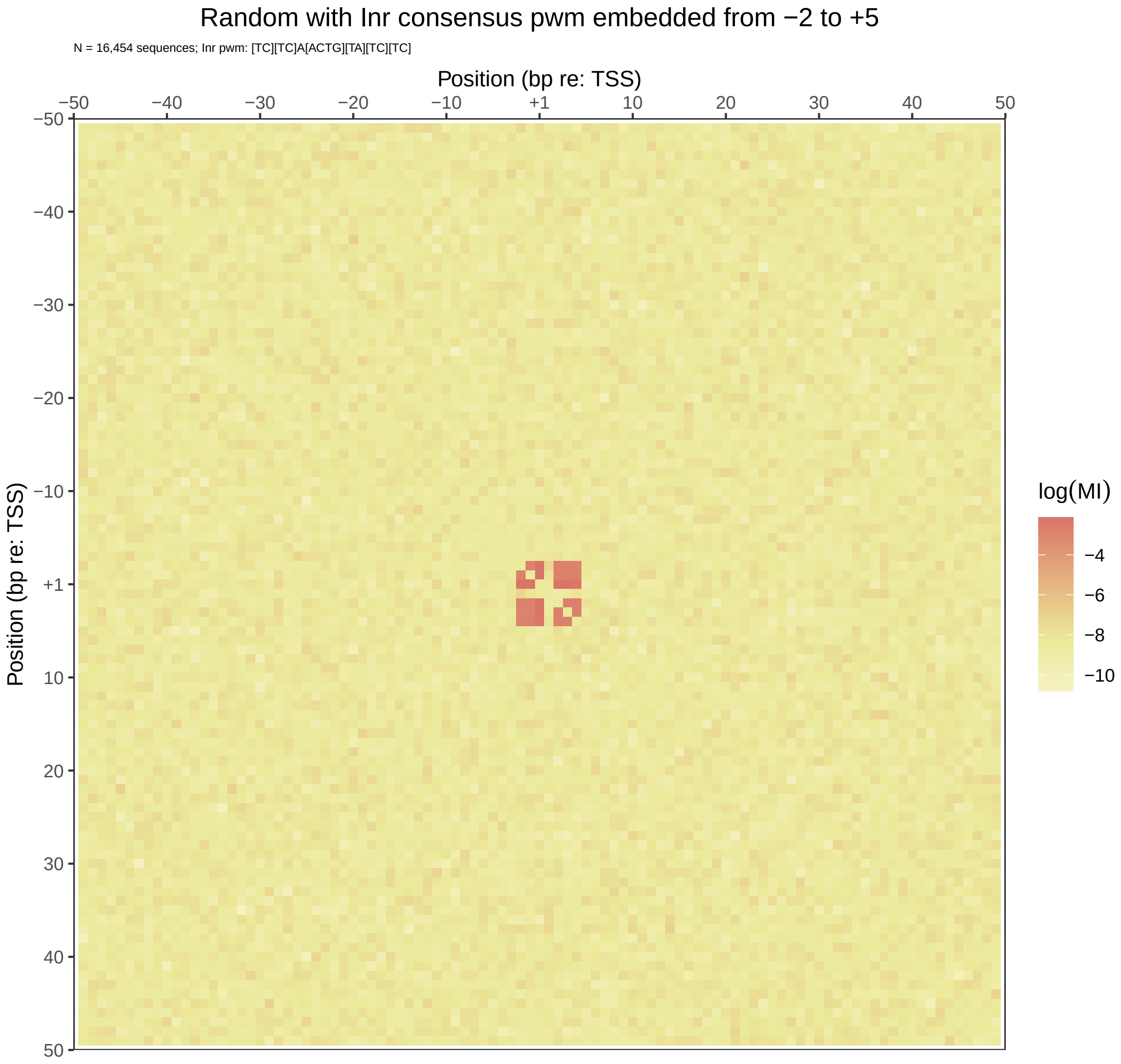


Figure 7: Heatmap produced from artificial sequences with a fixed – length motif embedded in 80% of aligned sequences at exactly the same position. The embedded motif is actually a collection of similar motifs with variants at some of the positions. In this case, the consensus sequence for the Inr motif, YYANWYY, was embedded starting at position -2. IUPAC symbols for variants (e.g., Y = T/C) were used to create each motif variant with the permissible nucleotides at any position being equiprobable, and the nucleotides in all of the positions (with the exception of the third position, A) being chosen independently.

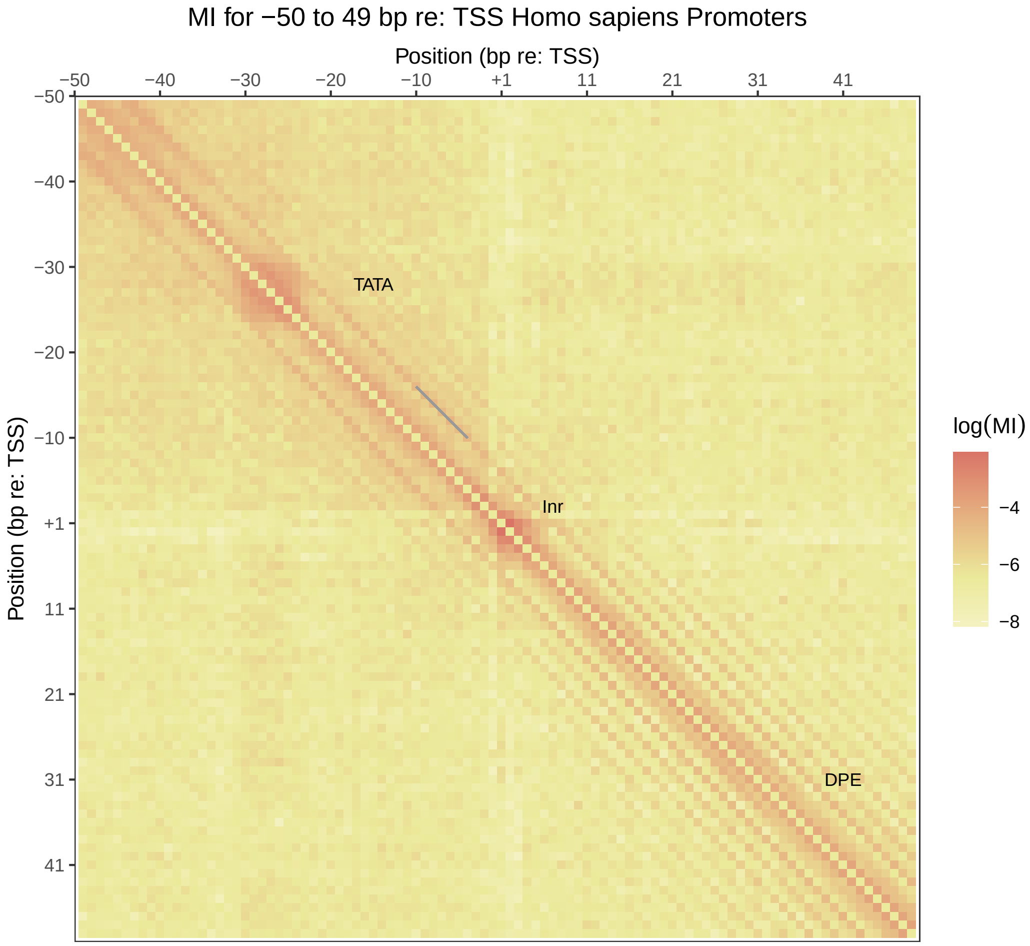


Figure 8: An example of an observed heatmap with a diagonally-running feature at a distance of 6 bp.

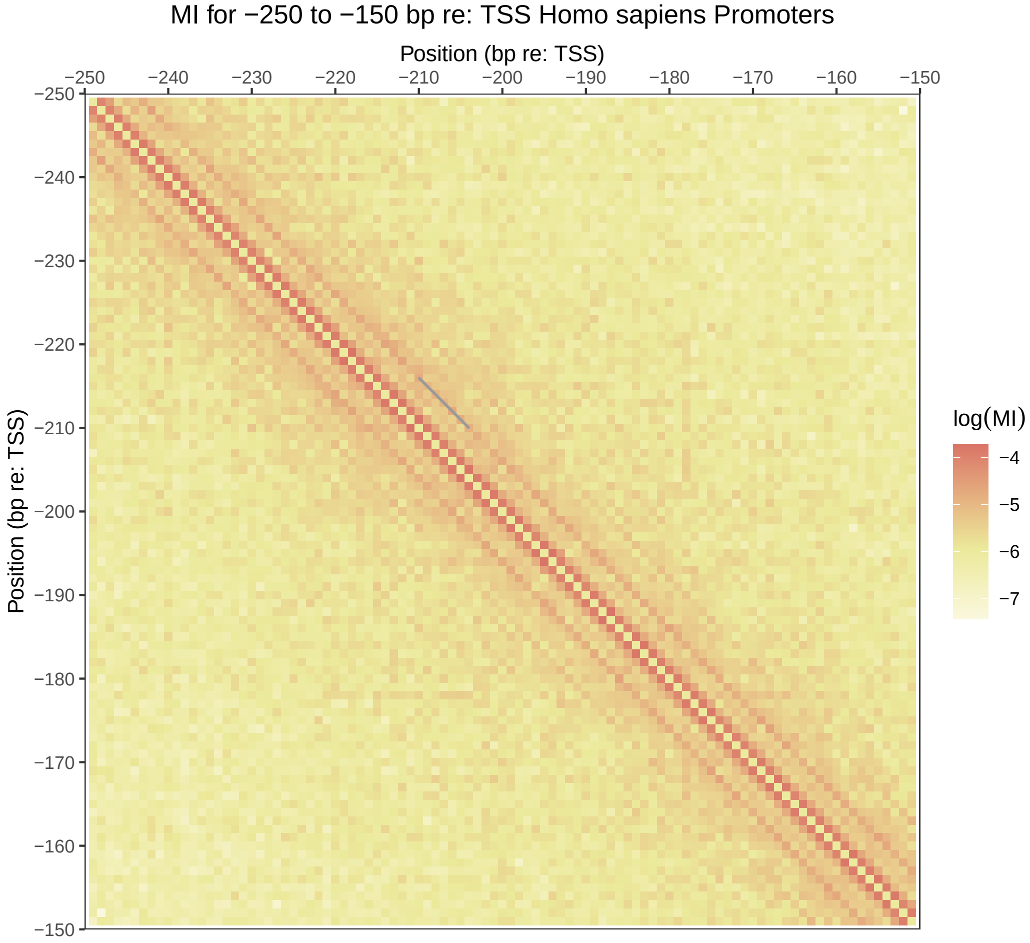


Figure 9: An example of an observed heatmap with a diagonally-running feature at a distance of 6 bp.

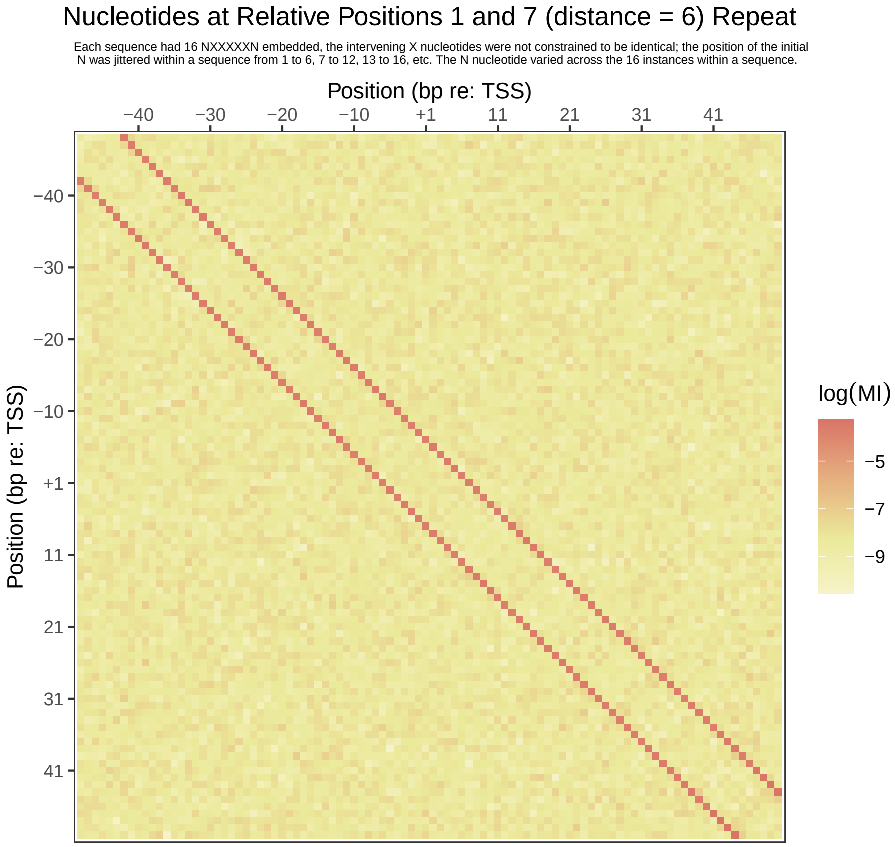


Figure 10: Artificial heatmap produced by embedding short sequences in which the first and the seventh nucleotides were always the same, but the intervening nucleotides were free to vary: e.g., AGTCAGA or more generally NVWXYZN. Sixteen of these short sequences were embedded in each otherwise random sequence 100 bp in width. The nucleotide that repeated, N, was chosen randomly for each of the 16 instances that were embedded in a given sequence. The starting position of the first embedded subsequence was chosen randomly from positions 1 to 6, the second from positions 7 to 12, etc. within each 100 bp sequence until all 16 subsequences had been embedded. Note: the position of the initial N in the last of the 16 embedded subsequence would allow the second N in the subsequence to occupy a position outside the 100 bp range. This was done, and the MI matrix was then trimmed to $100 \times 100$.

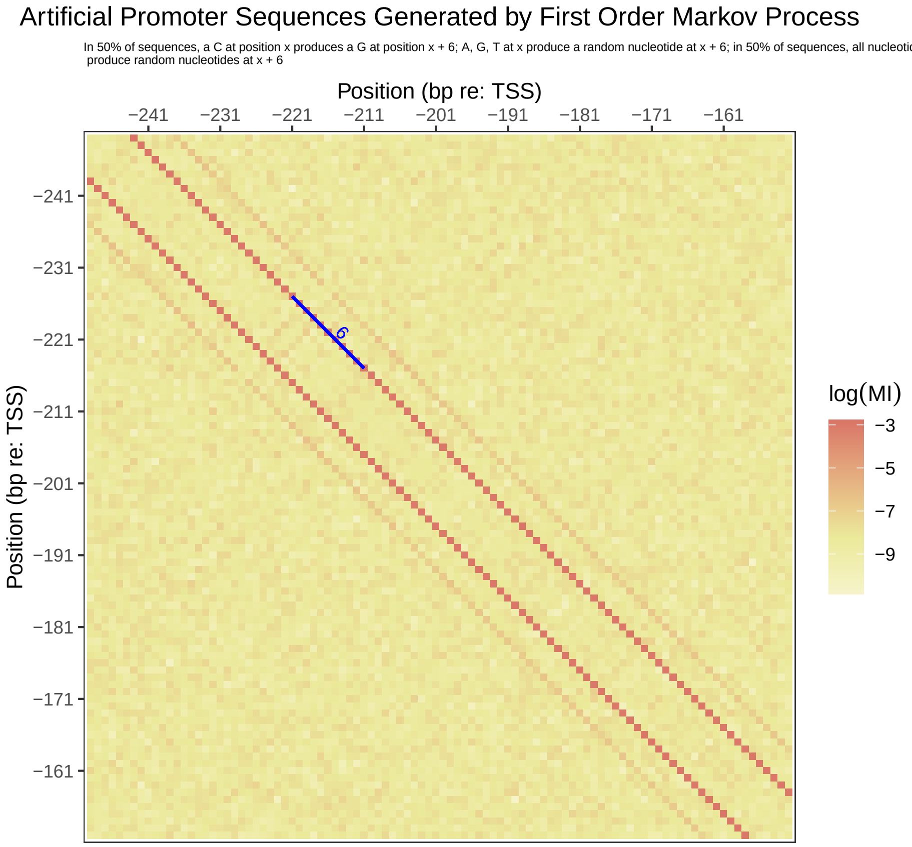


Figure 11: Artificial sequences generated using a first order Markov process. See Table 1 (bottom matrix) for the state transition matrix. When there was a dependency between nucleotides at different positions introduced by the Markov process, it was always at a distance of 6 bp. That is, the nucleotide at a given position depended on the nucleotide at the position 6 bp upstream.

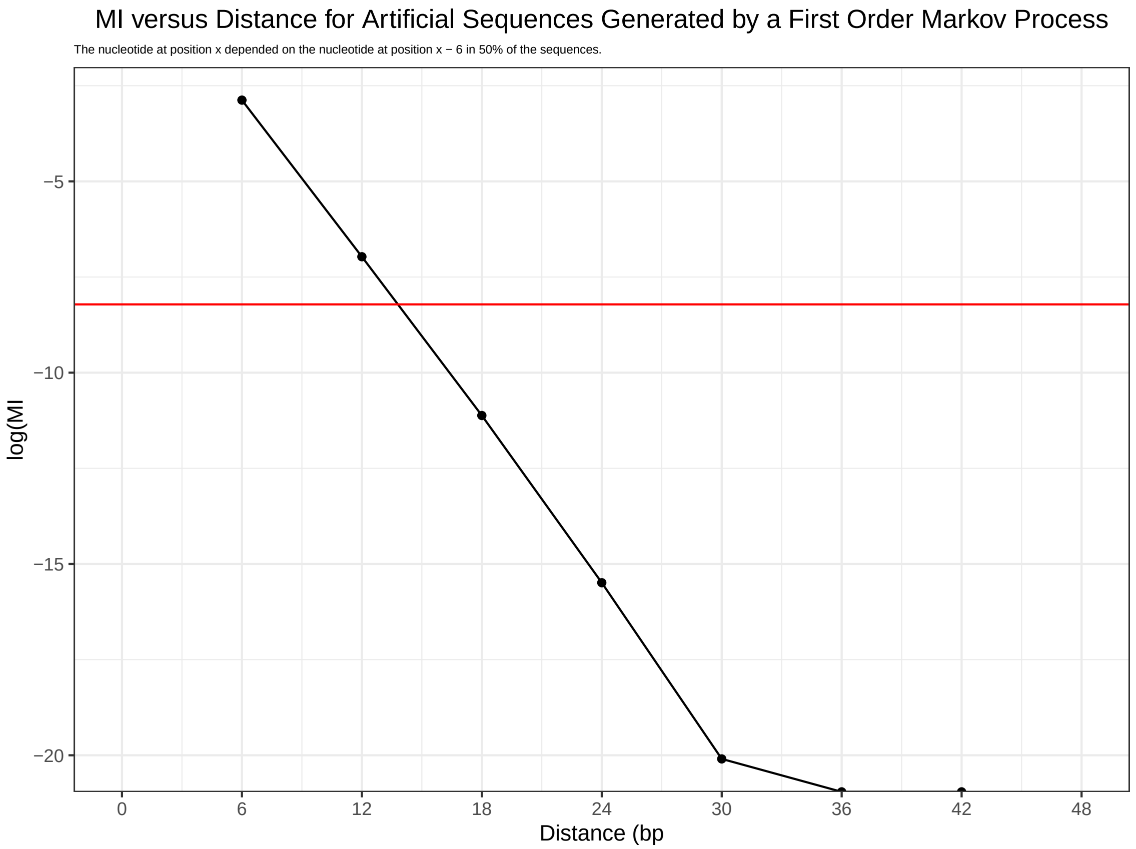


Figure 12: Log MI vs. distance for artificial sequences generated using a first order Markov process. See Table 1 (bottom matrix) for the state transition matrix. The red horizontal line is the mean of all of the off-diagonal log MI values. Any log MI value below the red line should not be detectable above the background noise in a heatmap like the one shown in Figure 11. In that figure, only the signals as distances of 6 and 12 bp are detectable as predicted from this curve.

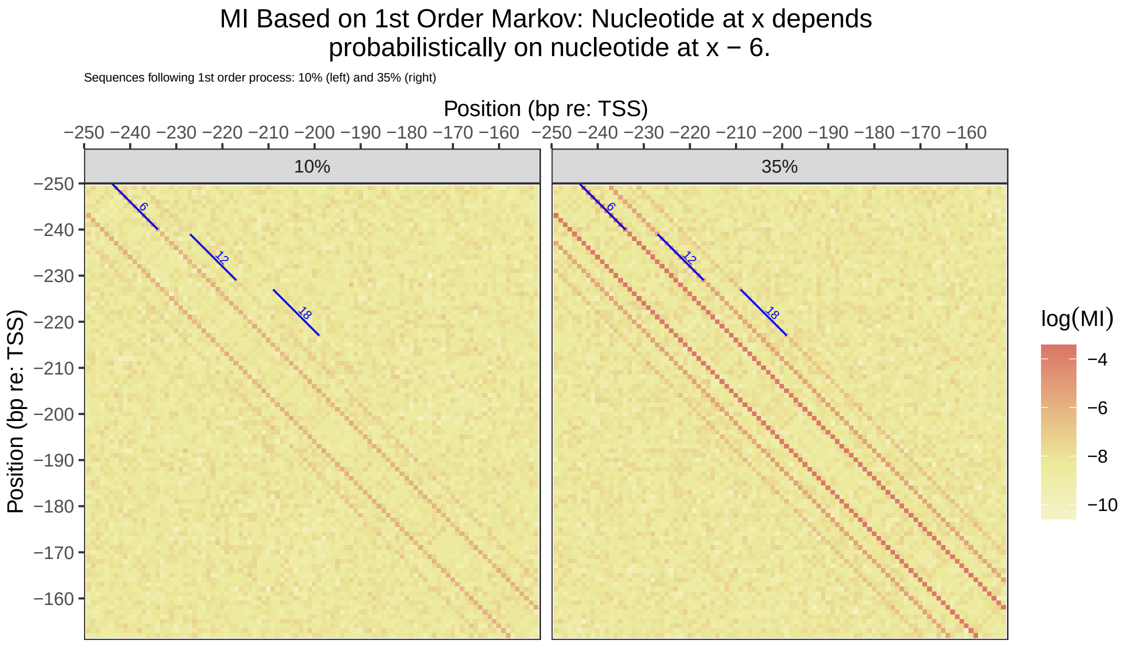


Figure 13: MI matrix heatmaps based on First Order Markov process in which the nucleotide at position x depends probabilistically on the nucleotide at position x - 6. (Left) 10% of the sequences generated using First Order Markov process and 90% random. (Right) 35% of the sequences generated using First Order Markov process and 65% random. Please see text for an explanation of why lines at multiples of a distance of 6 bp are present and visible.

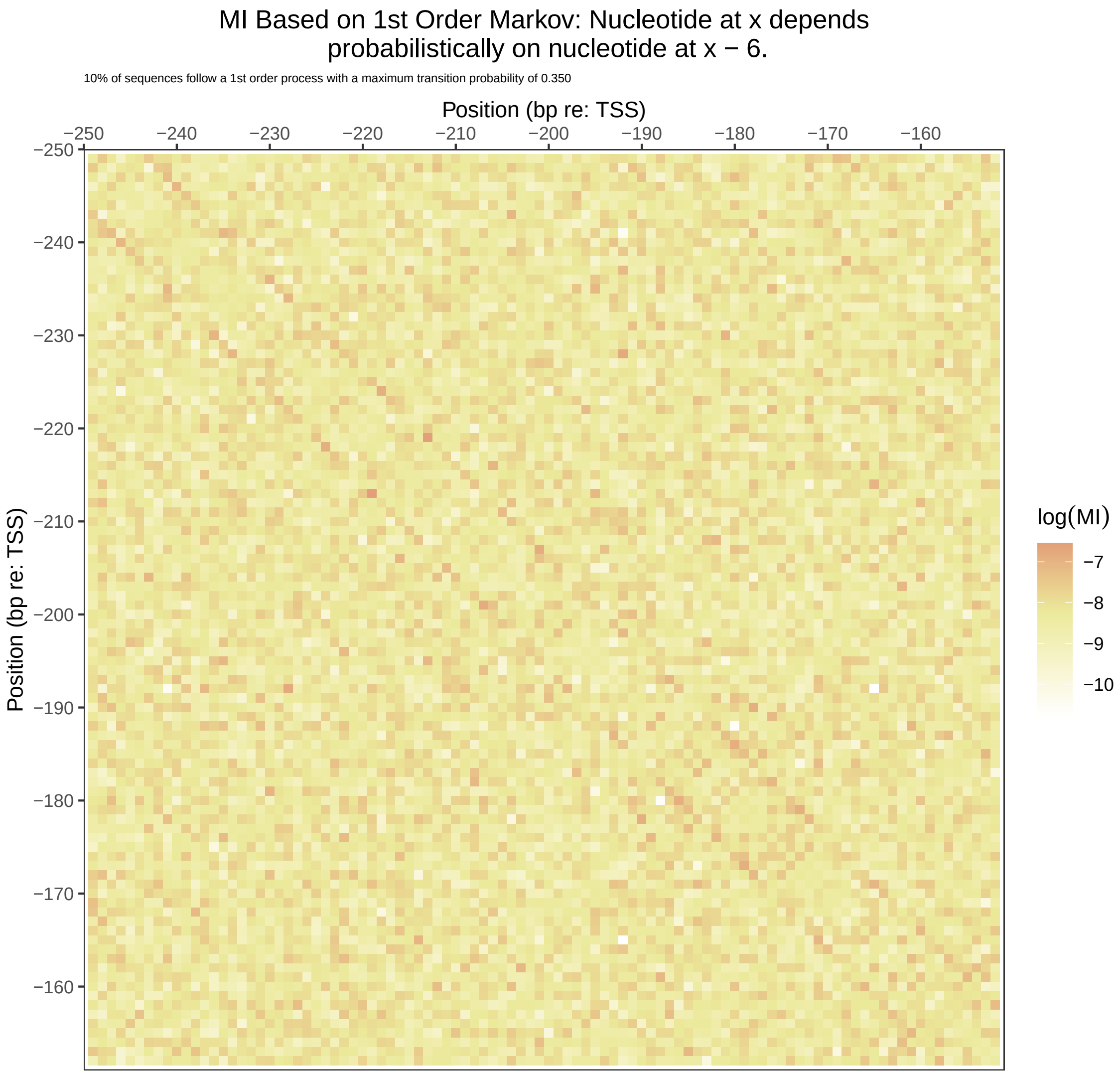


Figure 14: MI matrix heatmaps based on First Order Markov process in which the nucleotide at position x depends probabilistically on the nucleotide at position x - 6. All of the sequences were generated using a transition probability matrix like the one shown at the bottom of Table 2. The maximum probability in each row was 26% with the remaining probability equally divided between the other three nucleotides.

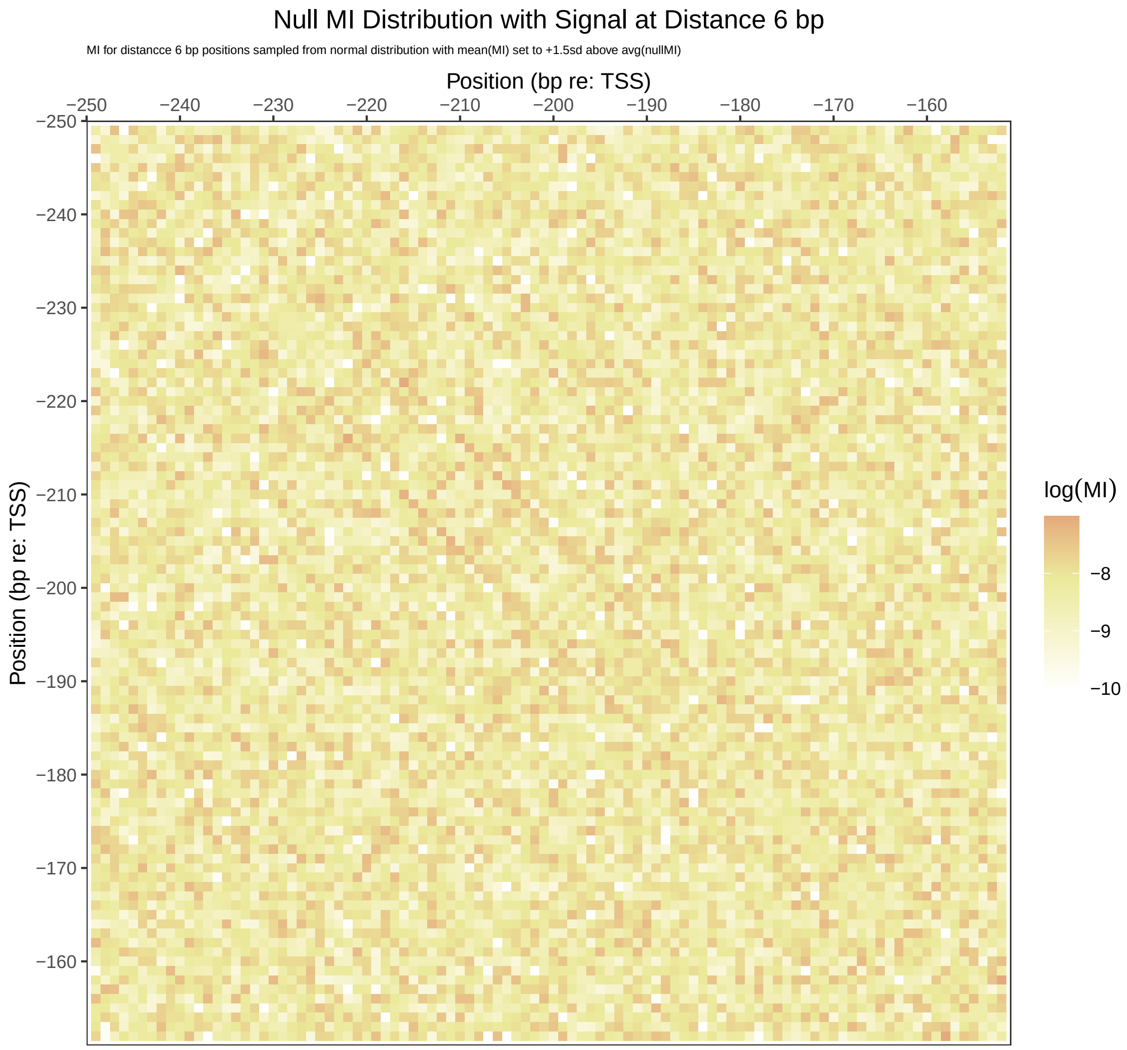


Figure 15: Heatmap based on a null MI matrix with the positions at a distance of 6 bp sampled from a normal distribution with a mean that was 1.5SD above the mean of the null MI values and a standard deviation equal to the SD of the null MI matrix.

Table 1: Transition matrix for first-order Markov process in which the nucleotide at position x depends in 25% of the sequences on the nucleotide at position x - 6 (left matrix). In the other 75% of the sequences the relation between the nucleotides at these two positions is random.

| | | Left: 50% | | | | Right: 50% | | | |
|---|---|---|---|---|---|---|---|---|---|
| | | Position: x + 6 | | | | Position: x + 6 | | | |
| Position | | A | C | G | T | A | C | G | T |
| | A | 0.25 | 0.25 | 0.25 | 0.25 | 0.25 | 0.25 | 0.25 | 0.25 |
| x | C | 0 | 0 | 1.0 | 0 | 0.25 | 0.25 | 0.25 | 0.25 |
| | G | 0.25 | 0.25 | 0.25 | 0.25 | 0.25 | 0.25 | 0.25 | 0.25 |
| | T | 0.25 | 0.25 | 0.25 | 0.25 | 0.25 | 0.25 | 0.25 | 0.25 |

| | | Resulting Transition Matrix | | | |
|---|---|---|---|---|---|
| | | Position: x + 6 | | | |
| Position | | A | C | G | T |
| | A | 0.25 | 0.25 | 0.25 | 0.25 |
| x | C | 0.125 | 0.125 | 0.625 | 0.125 |
| | G | 0.25 | 0.25 | 0.25 | 0.25 |
| | T | 0.25 | 0.25 | 0.25 | 0.25 |

Table 2: Transition matrix for first-order Markov process in which the nucleotide at position x depends in 10% of the sequences on the nucleotide at position x - 6 (left matrix). In the other 90% of the sequences the relation between the nucleotides at these two positions is random.

| | | Left: 50% Position: x + 6 | | | | Right: 50% Position: x + 6 | | | |
|---|---|---|---|---|---|---|---|---|---|
| Position | | A | C | G | T | A | C | G | T |
| | A | 0.55 | 0.15 | 0.15 | 0.15 | 0.25 | 0.25 | 0.25 | 0.25 |
| x | C | 0.15 | 0.55 | 0.15 | 0.15 | 0.25 | 0.25 | 0.25 | 0.25 |
| | G | 0.15 | 0.15 | 0.55 | 0.15 | 0.25 | 0.25 | 0.25 | 0.25 |
| | T | 0.15 | 0.15 | 0.15 | 0.55 | 0.25 | 0.25 | 0.25 | 0.25 |

| | | Resulting Transition Matrix Position: x + 6 | | | |
|---|---|---|---|---|---|
| Position | | A | C | G | T |
| | A | 0.28 | 0.24 | 0.24 | 0.24 |
| x | C | 0.24 | 0.28 | 0.24 | 0.24 |
| | G | 0.24 | 0.24 | 0.28 | 0.24 |
| | T | 0.24 | 0.24 | 0.24 | 0.28 |
