## Supplementary material for "Mutual information reveals a homonucleotide bias at 6 bp distance in promoters": Supplementary-Figures.pdf

### 1 Supplementary Figures

All Mutual Information Heatmaps

Mi vs. Distance for aligned and misaligned promoter sequences

Observed and expected proportions of like – nucleotides

#### 1.1 All Mutual Information Heatmaps

For each of the species Homo sapiens, Drosophila melanogaster, Saccharomyces cerevisiae, and Mus musculus, ten heatmaps covering the range from -950 to + 49 bp re: TSS (+1) are presented. The subrange covered by each heatmap is included in the title of each plot. Features of these heatmaps are discussed in the text. They are presented here for completeness.

**Homo sapiens**

**-950:-851** Figure S1

**-850:-751** Figure S2

**-750:-651** Figure S3

**-650:-551** Figure S4

**-550:-451** Figure S5

**-450:-351** Figure S6

**-350:-251** Figure S7

**-250:-151** Figure S8

**-150: -51** Figure S9

**-50: +49** Figure S10

**Drosophila melanogaster**

**-950:-851** Figure S11

**-850:-751** Figure S12

**-750:-651** Figure S13

**-650:-551** Figure S14

**-550:-451** Figure S15

**-450:-351** Figure S16

**-350:-251** Figure S17

**-250:-151** Figure S18

**-150: -51** Figure S19

**-50: +49** Figure S20

**Saccharomyces cerevisiae**

**-950:-851** Figure S21

**-850:-751** Figure S22

**-750:-651** Figure S23

**-650:-551** Figure S24

**-550:-451** Figure S25

**-450:-351** Figure S26

**-350:-251** Figure S27

**-250:-151** Figure S28

**-150: -51** Figure S29

**-50: +49** Figure S30

**Mus musculus**

**-950:-851** Figure S31

**-850:-751** Figure S32

**-750:-651** Figure S33

**-650:-551** Figure S34

**-550:-451** Figure S35

**-450:-351** Figure S36

**-350:-251** Figure S37

**-250:-151** Figure S38

**-150: -51** Figure S39

**-50: +49** Figure S40

#### 1.2 MI vs. Distance for aligned and misaligned sequences.

| | |
|---|---|
| *Homo sapiens* | *Figure* S41 |
| *Drosophila melanogaster* | *Figure* S42 |
| *Saccharomyces cerevisiae* | *Figure* S43 |
| *Mus musculus* | *Figure* S44 |

##### 1.3 Observed and expected proportions of like – nucleotides at distances from 1 to 21 bp.

| | |
|---|---|
| *Homo sapiens* | *Figure* S45 |
| *Drosophila melanogaster* | *Figure* S46 |
| *Saccharomyces cerevisiae* | *Figure* S47 |
| *Mus musculus* | *Figure* S48 |

#### 1.4 Opportunity (length) corrected MI vs. Distance

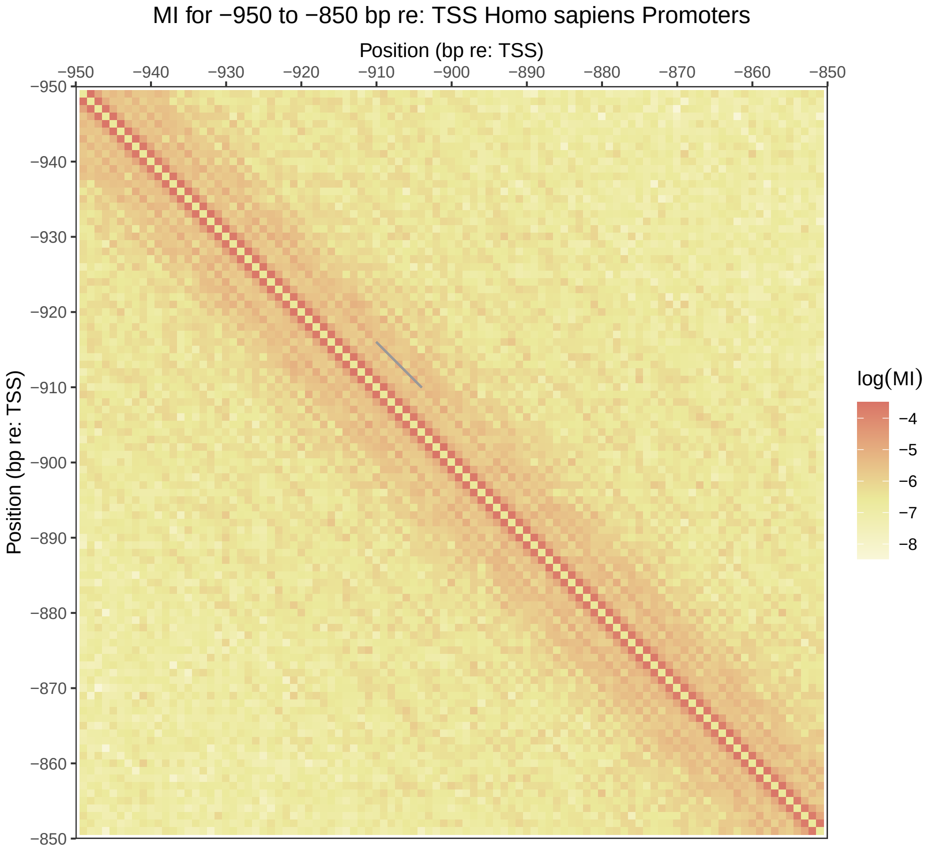
MI for −950 to −850 bp re: TSS Homo sapiens Promoters
Position (bp re: TSS)
−950
−940
−930
−920
−910
−900
−890
−880
−870
−860
−850
Position (bp re: TSS)
log(MI)
−4
−5
−6
−7
−8

Figure S1

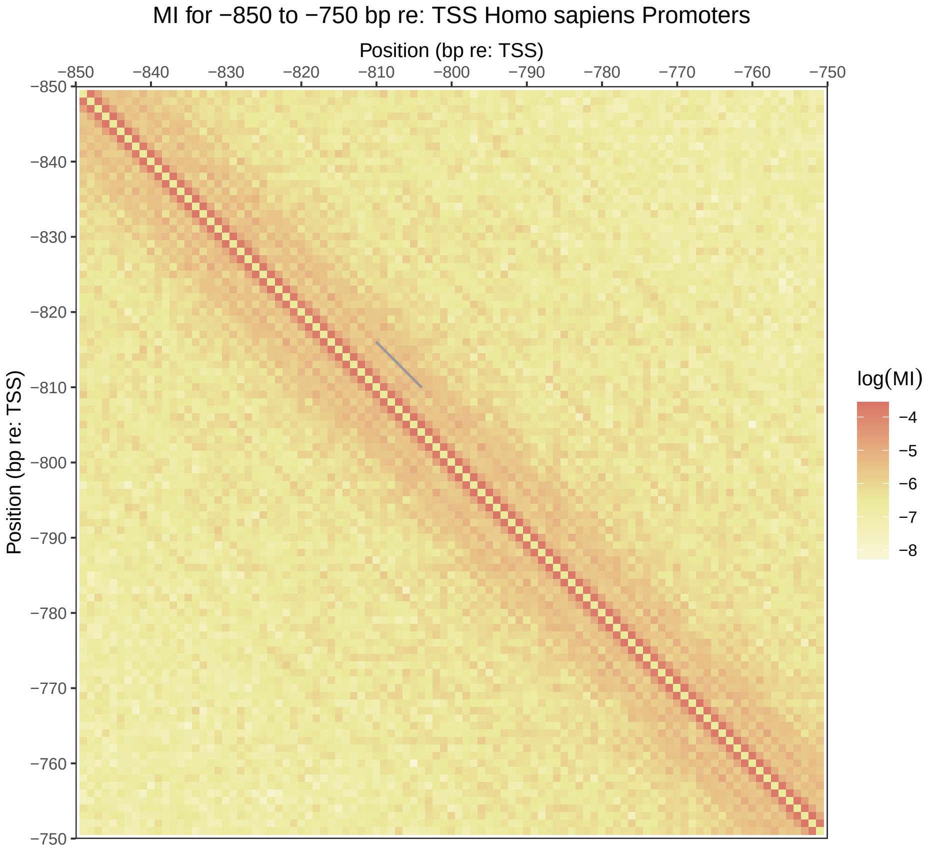
MI for −850 to −750 bp re: TSS Homo sapiens Promoters
Position (bp re: TSS)
−850
−840
−830
−820
−810
−800
−790
−780
−770
−760
−750
Position (bp re: TSS)
log(MI)
−4
−5
−6
−7
−8

Figure S2

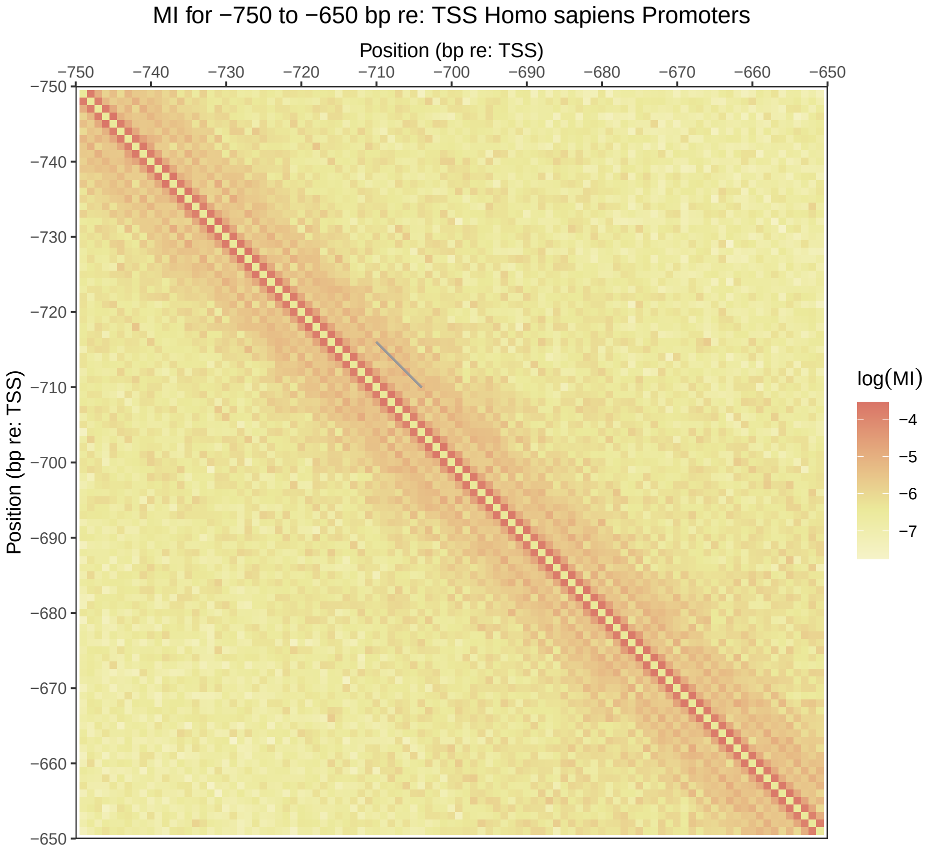
MI for −750 to −650 bp re: TSS Homo sapiens Promoters
Position (bp re: TSS)
−750
−740
−730
−720
−710
−700
−690
−680
−670
−660
−650
Position (bp re: TSS)
log(MI)
−4
−5
−6
−7

Figure S3

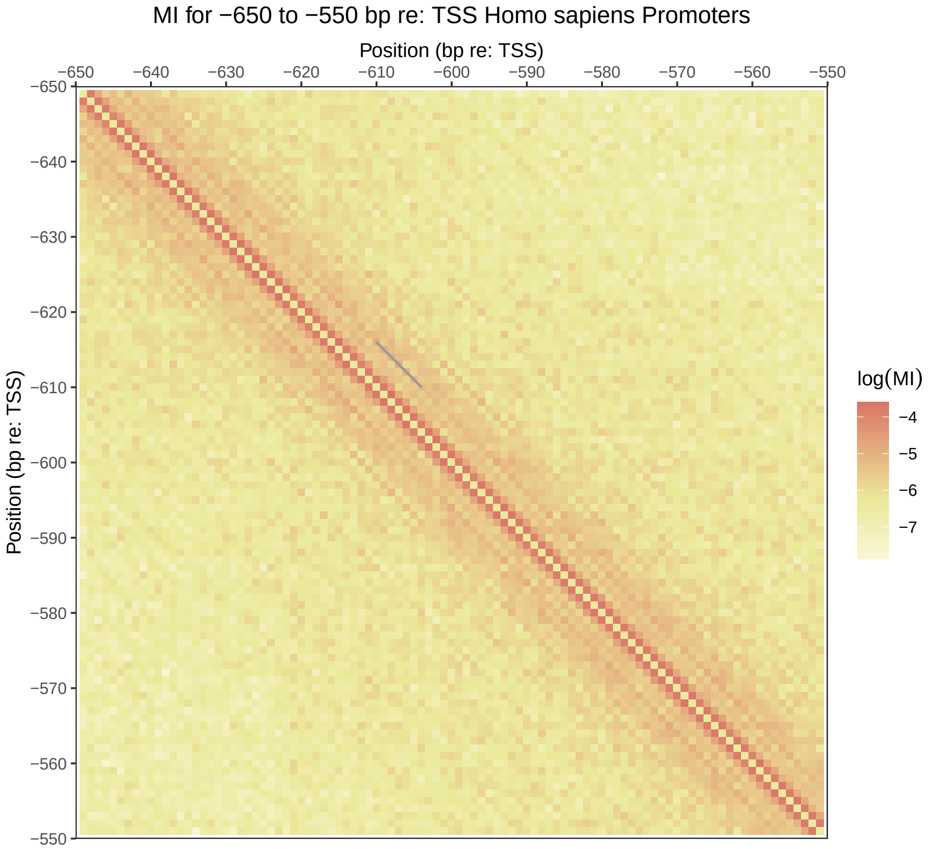
MI for −650 to −550 bp re: TSS Homo sapiens Promoters
Position (bp re: TSS)
−650
−640
−630
−620
−610
−600
−590
−580
−570
−560
−550
Position (bp re: TSS)
log(MI)
−4
−5
−6
−7

Figure S4

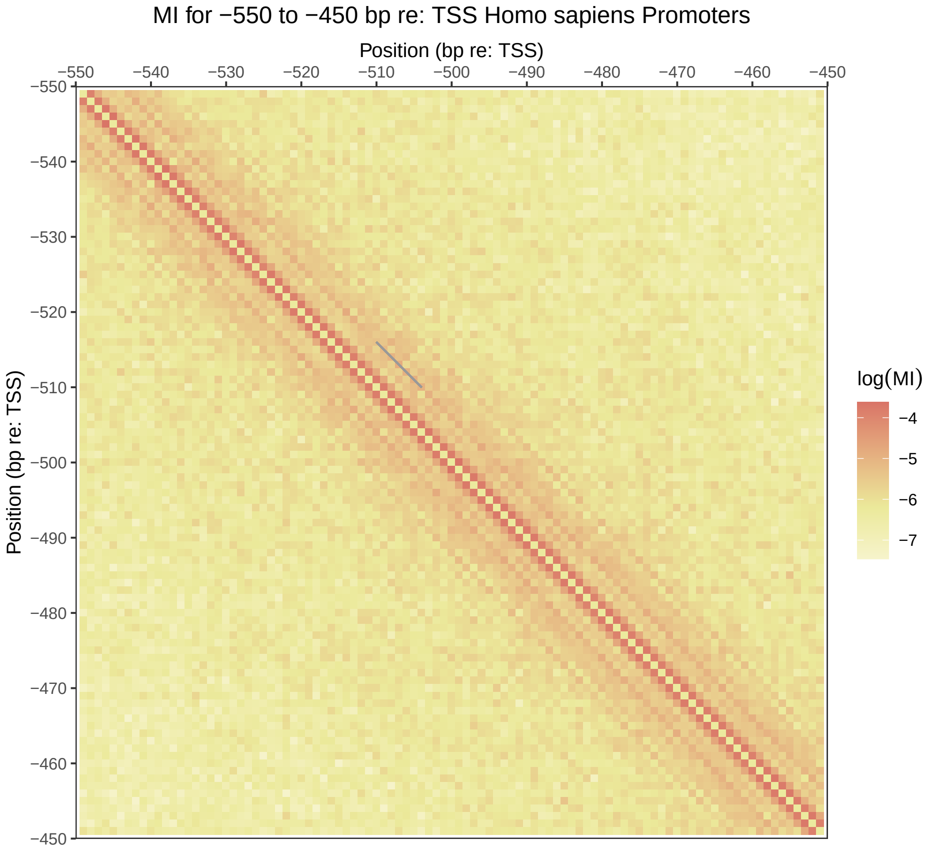
MI for −550 to −450 bp re: TSS Homo sapiens Promoters
Position (bp re: TSS)
−550
−540
−530
−520
−510
−500
−490
−480
−470
−460
−450
Position (bp re: TSS)
log(MI)
−4
−5
−6
−7

Figure S5

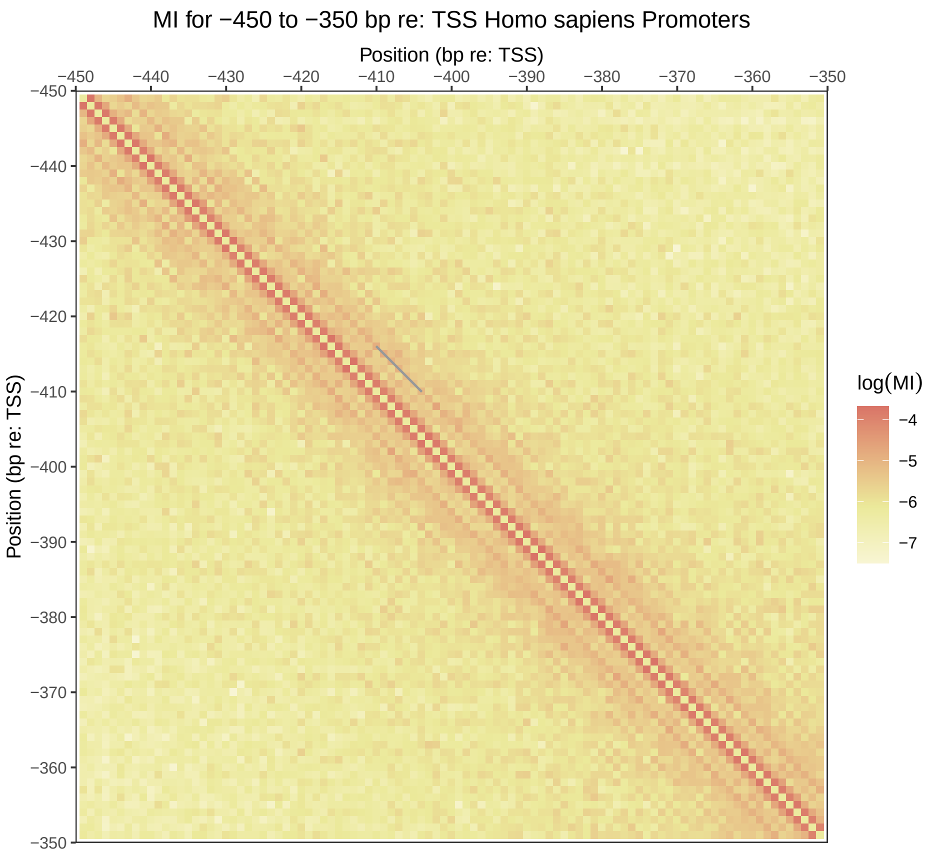
MI for −450 to −350 bp re: TSS Homo sapiens Promoters
Position (bp re: TSS)
−450
−440
−430
−420
−410
−400
−390
−380
−370
−360
−350
Position (bp re: TSS)
log(MI)
−4
−5
−6
−7

Figure S6

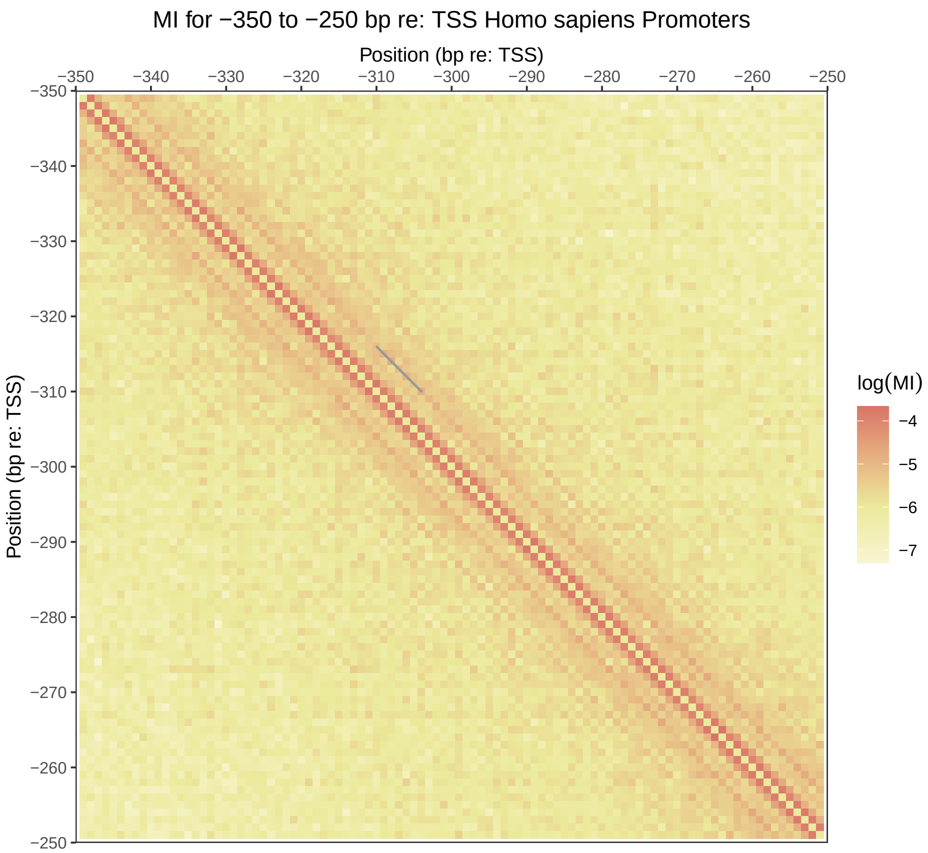
MI for −350 to −250 bp re: TSS Homo sapiens Promoters
Position (bp re: TSS)
−350
−340
−330
−320
−310
−300
−290
−280
−270
−260
−250
Position (bp re: TSS)
log(MI)
−4
−5
−6
−7

Figure S7

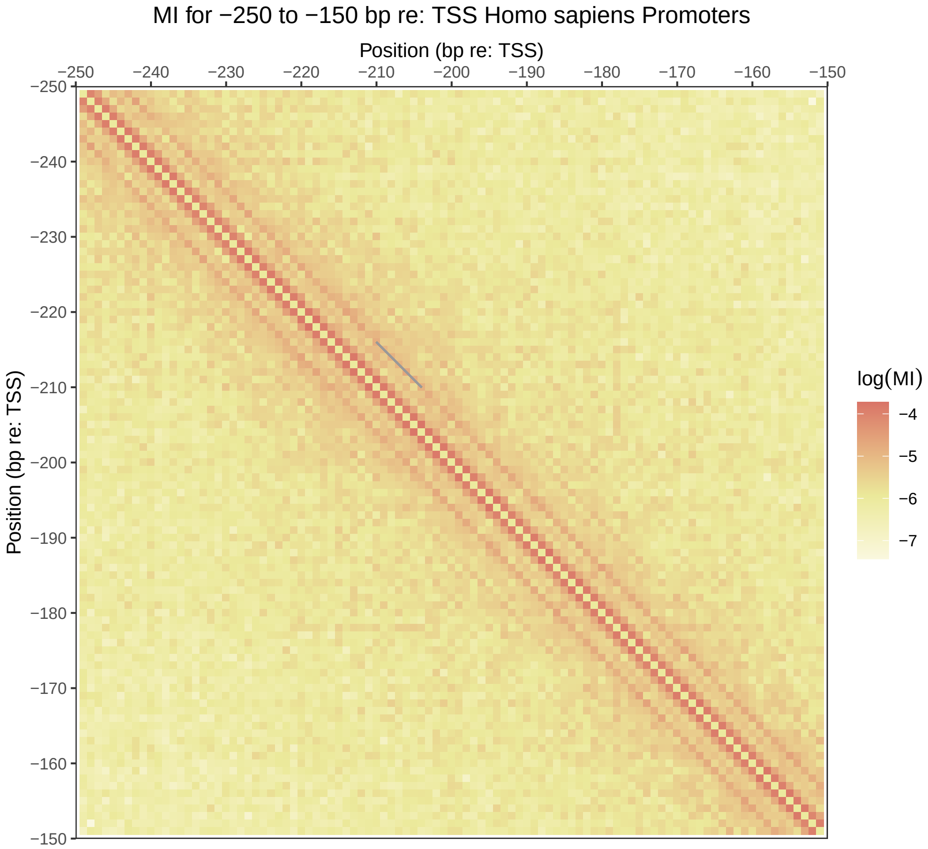
MI for −250 to −150 bp re: TSS Homo sapiens Promoters
Position (bp re: TSS)
−250
−240
−230
−220
−210
−200
−190
−180
−170
−160
−150
Position (bp re: TSS)
log(MI)
−4
−5
−6
−7

Figure S8

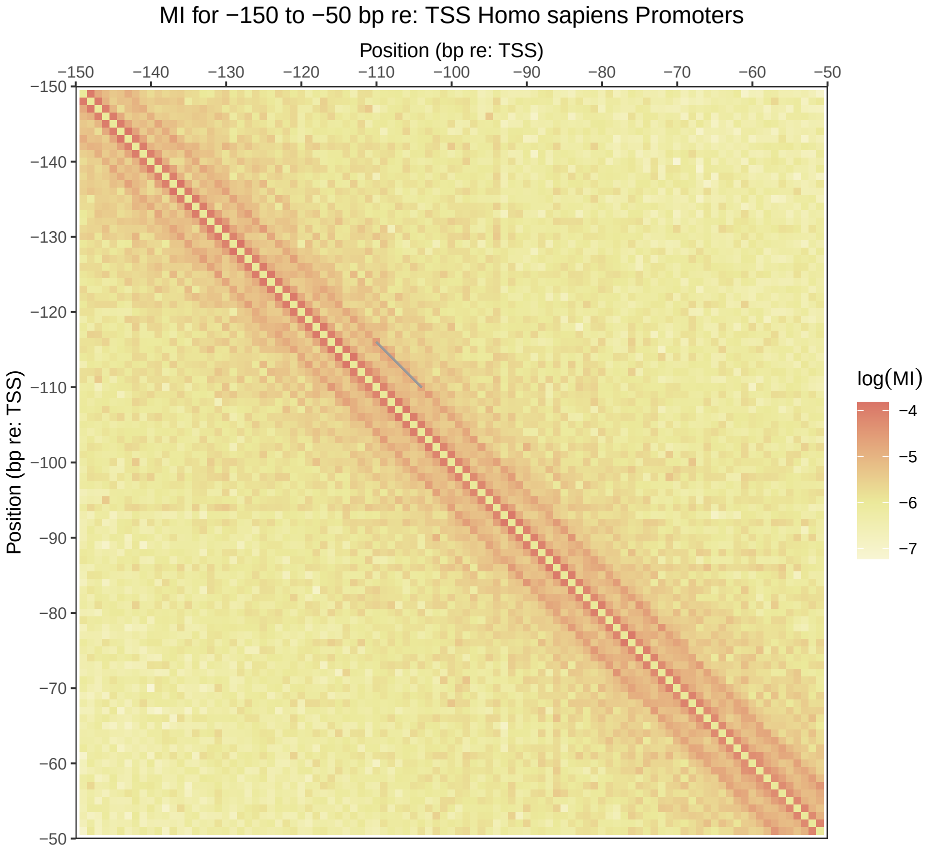

MI for −150 to −50 bp re: TSS Homo sapiens Promoters
Position (bp re: TSS)
−150
−140
−130
−120
−110
−100
−90
−80
−70
−60
−50
Position (bp re: TSS)
log(MI)
−4
−5
−6
−7


Figure S9

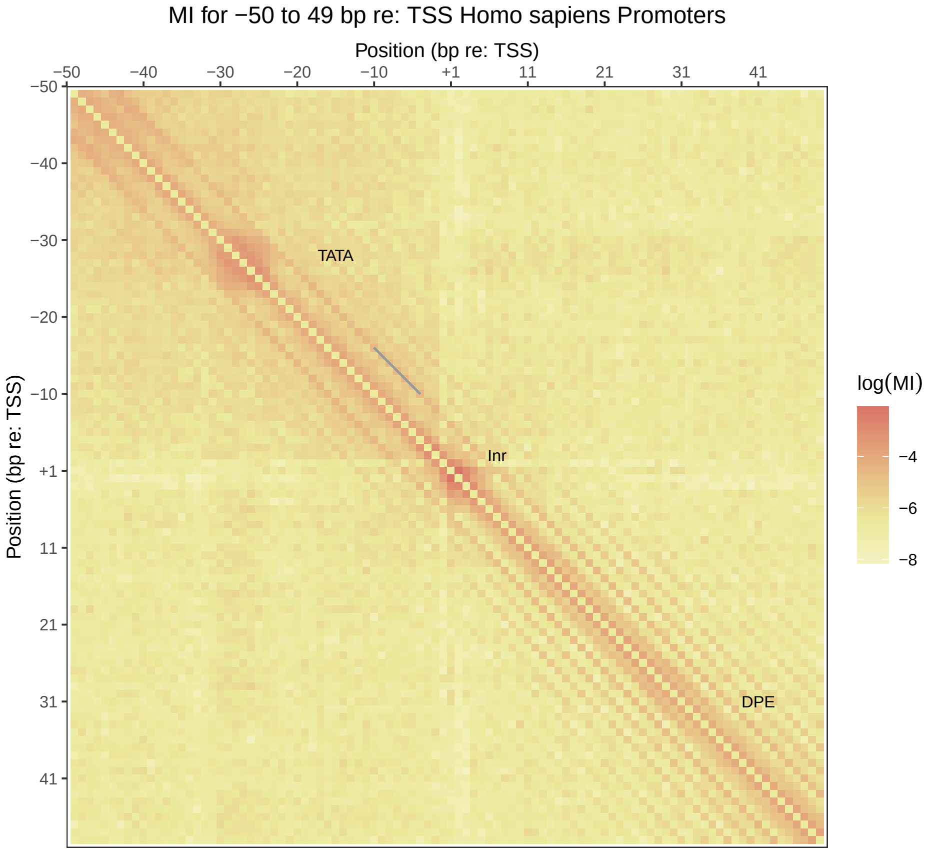
MI for −50 to 49 bp re: TSS Homo sapiens Promoters
Position (bp re: TSS)
−50
−40
−30
−20
−10
+1
11
21
31
41
Position (bp re: TSS)
TATA
Inr
DPE
log(MI)
−4
−6
−8

Figure S10

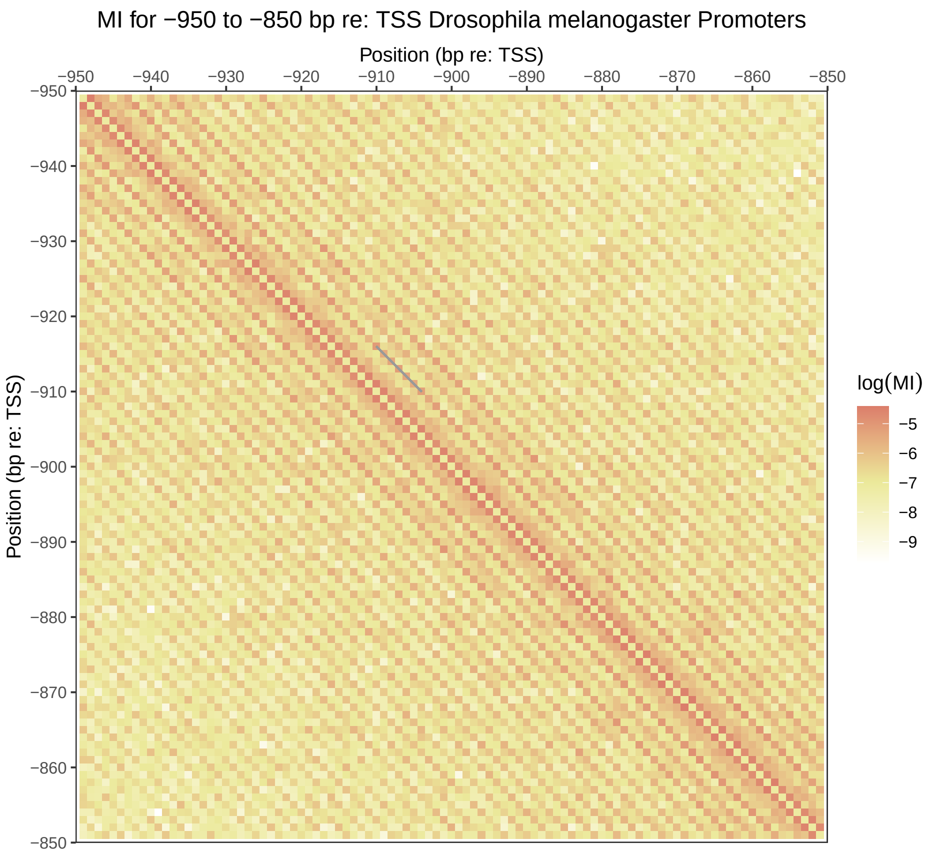
MI for −950 to −850 bp re: TSS Drosophila melanogaster Promoters
Position (bp re: TSS)
−950
−940
−930
−920
−910
−900
−890
−880
−870
−860
−850
Position (bp re: TSS)
log(MI)
−5
−6
−7
−8
−9

Figure S11

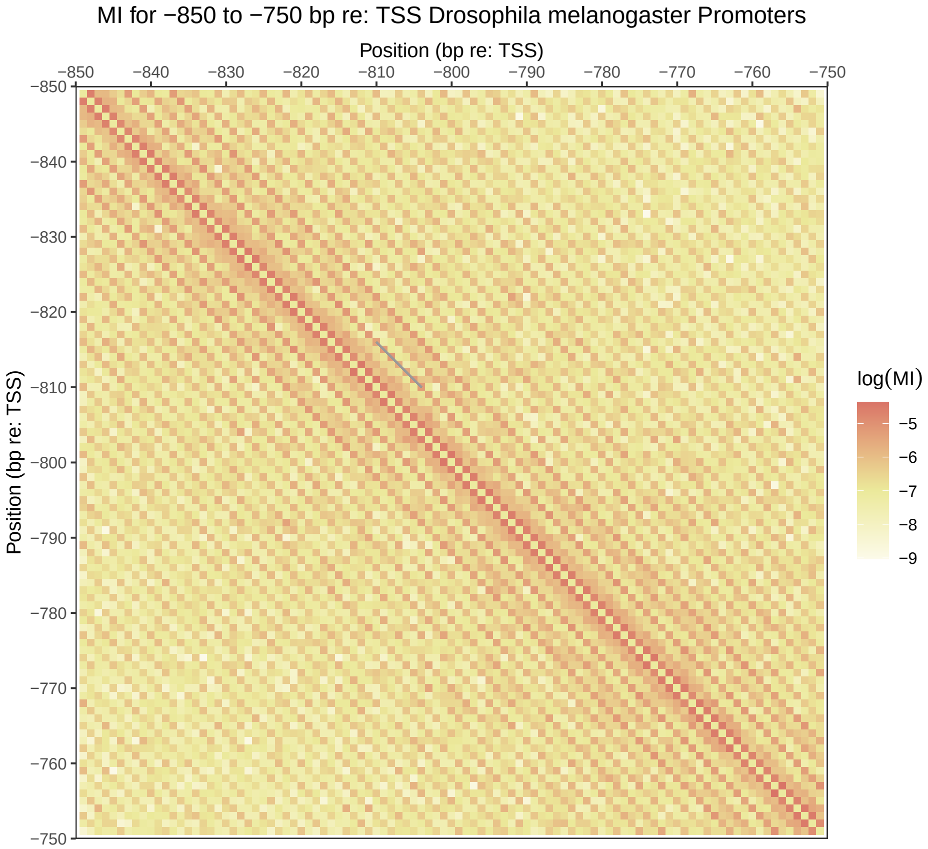
MI for −850 to −750 bp re: TSS Drosophila melanogaster Promoters
Position (bp re: TSS)
−850
−840
−830
−820
−810
−800
−790
−780
−770
−760
−750
Position (bp re: TSS)
log(MI)
−5
−6
−7
−8
−9

Figure S12

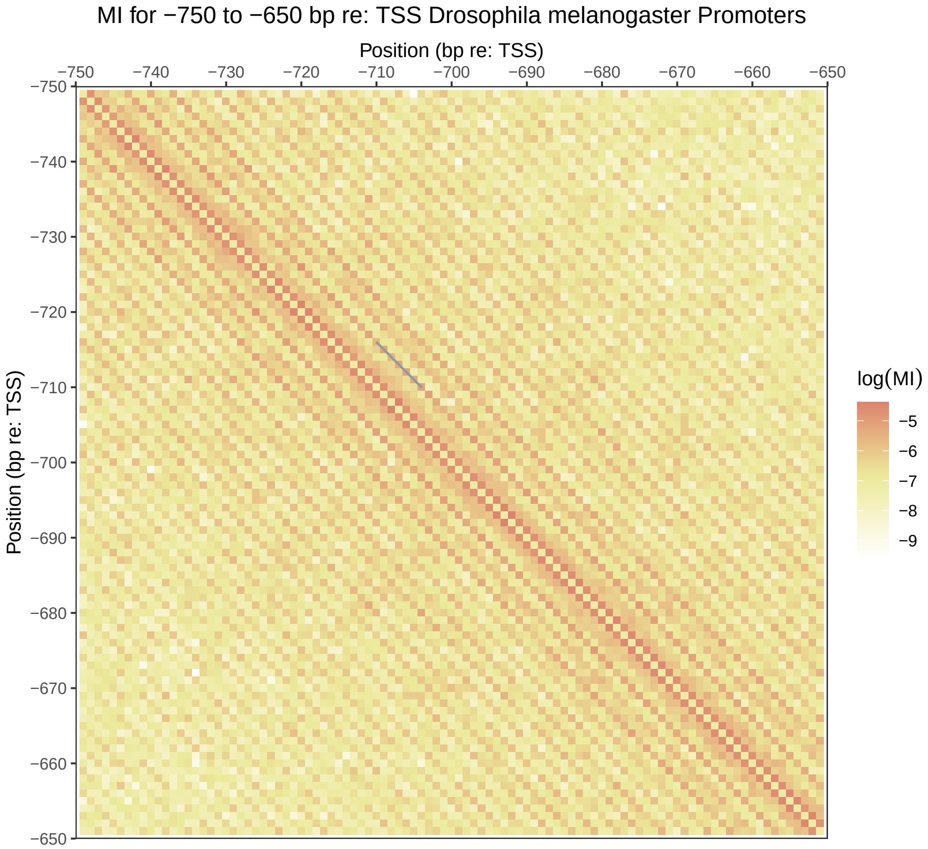
MI for −750 to −650 bp re: TSS Drosophila melanogaster Promoters
Position (bp re: TSS)
−750
−740
−730
−720
−710
−700
−690
−680
−670
−660
−650
Position (bp re: TSS)
log(MI)
−5
−6
−7
−8
−9

Figure S13

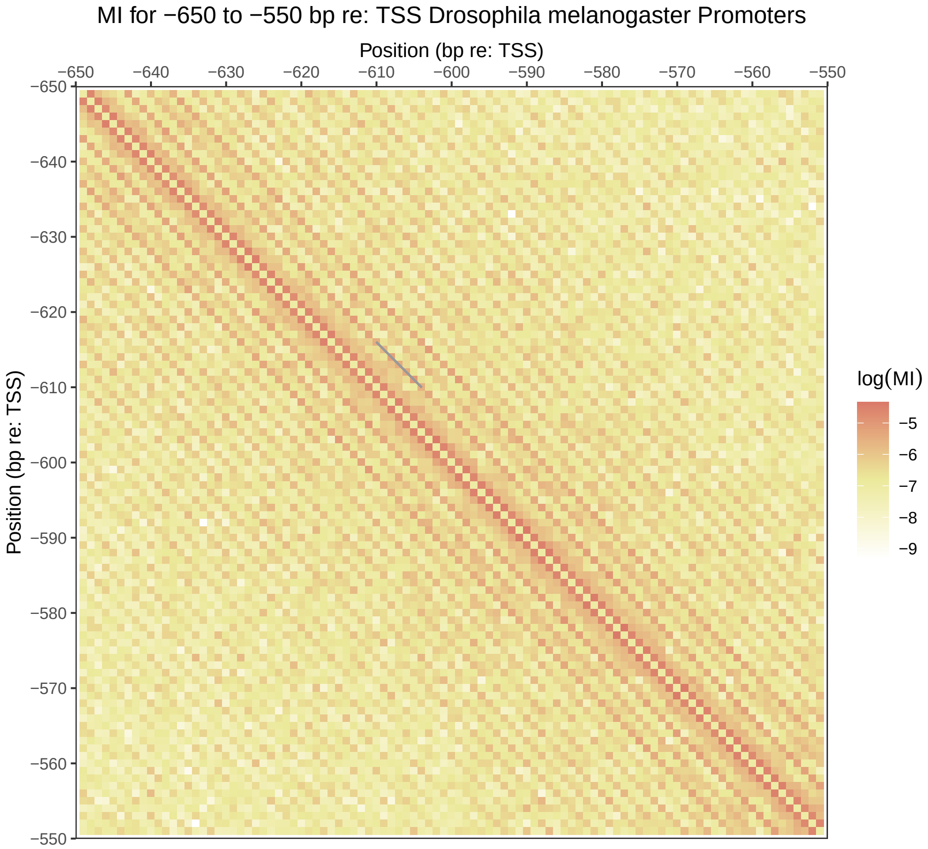
MI for −650 to −550 bp re: TSS Drosophila melanogaster Promoters
Position (bp re: TSS)
−650
−640
−630
−620
−610
−600
−590
−580
−570
−560
−550
Position (bp re: TSS)
log(MI)
−5
−6
−7
−8
−9

Figure S14

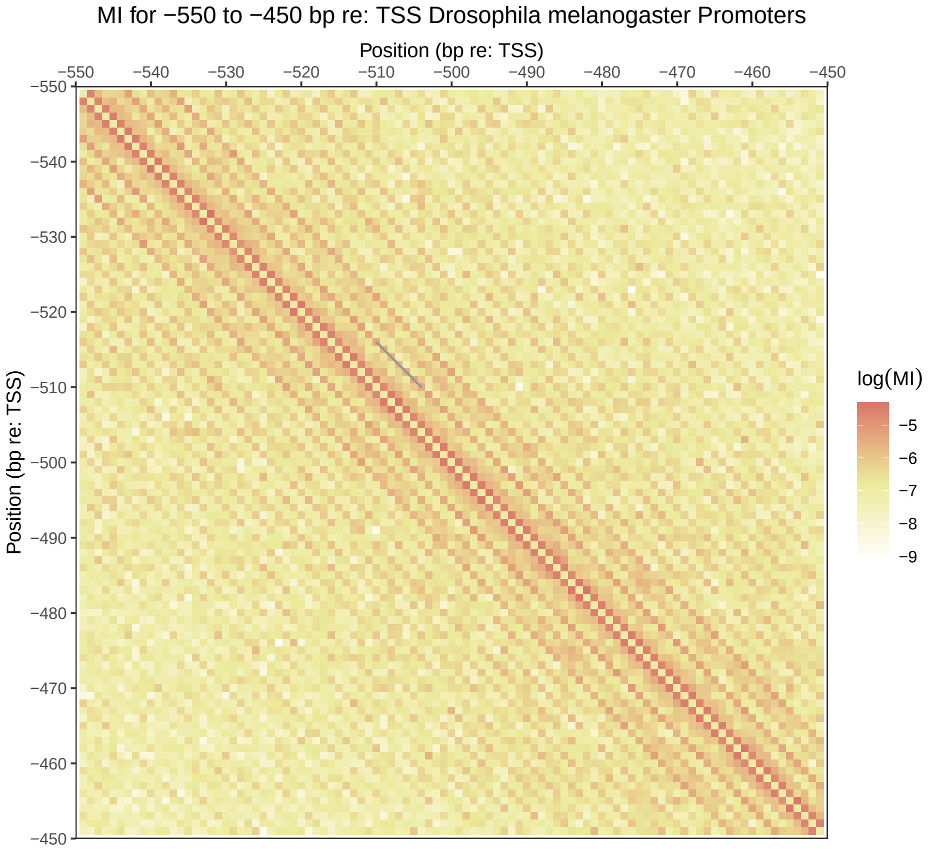
MI for −550 to −450 bp re: TSS Drosophila melanogaster Promoters
Position (bp re: TSS)
−550
−540
−530
−520
−510
−500
−490
−480
−470
−460
−450
Position (bp re: TSS)
log(MI)
−5
−6
−7
−8
−9

Figure S15

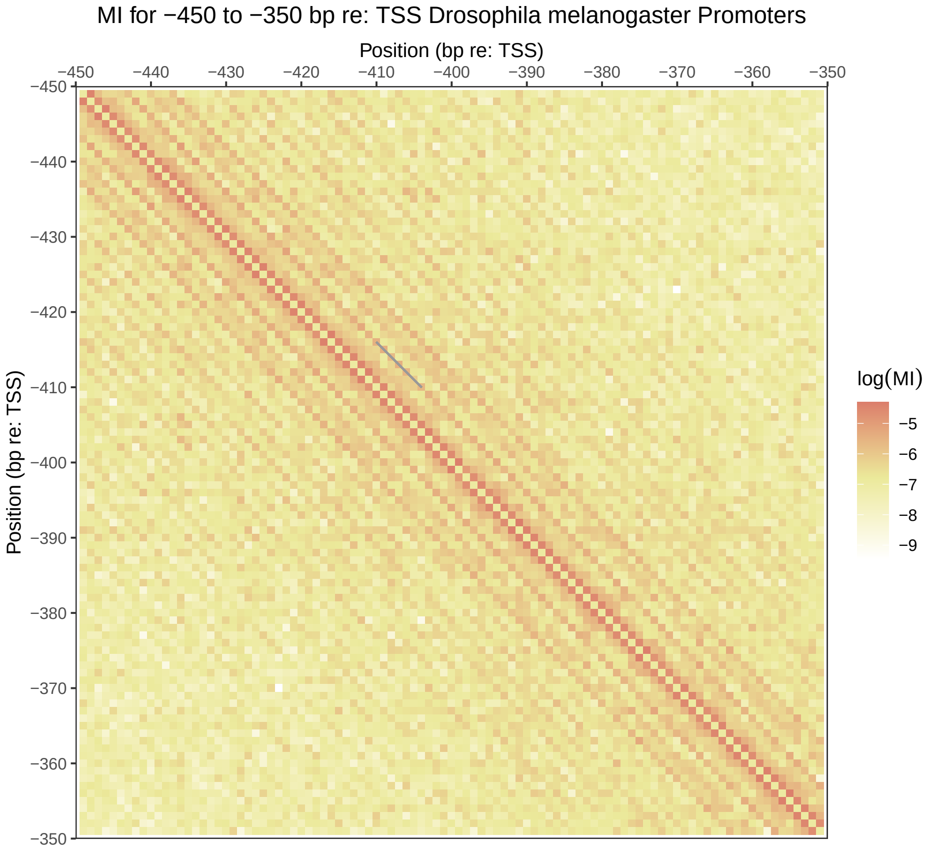
MI for −450 to −350 bp re: TSS Drosophila melanogaster Promoters
Position (bp re: TSS)
−450
−440
−430
−420
−410
−400
−390
−380
−370
−360
−350
Position (bp re: TSS)
log(MI)
−5
−6
−7
−8
−9

Figure S16

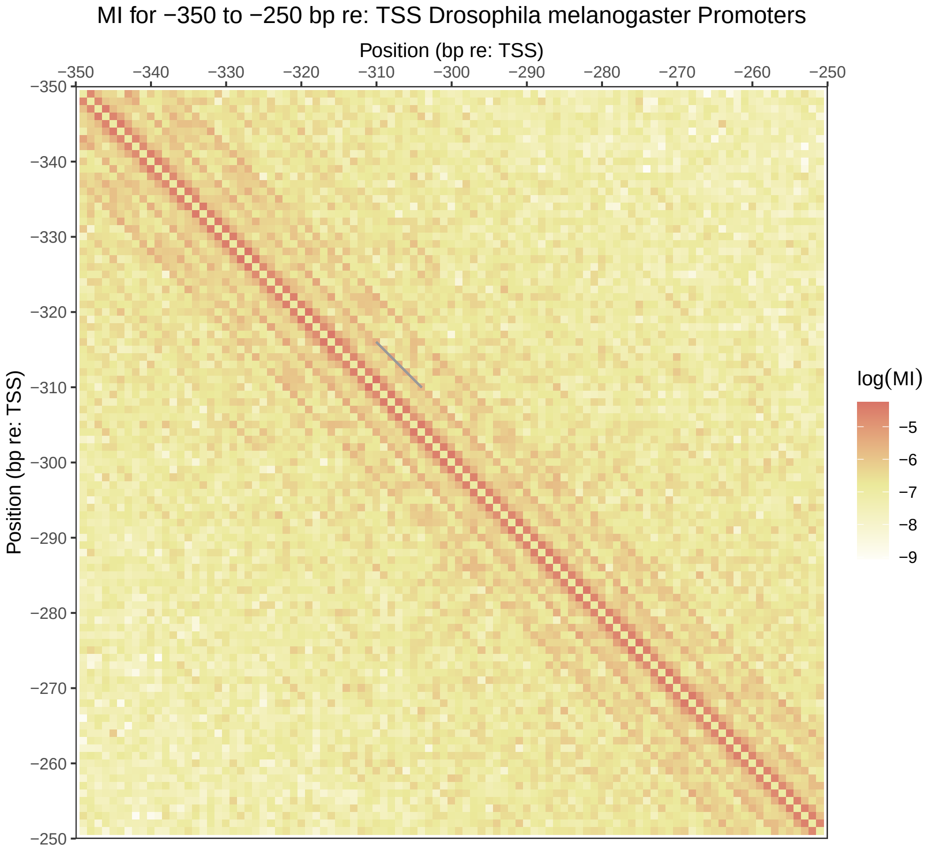
MI for −350 to −250 bp re: TSS Drosophila melanogaster Promoters
Position (bp re: TSS)
−350
−340
−330
−320
−310
−300
−290
−280
−270
−260
−250
Position (bp re: TSS)
log(MI)
−5
−6
−7
−8
−9

Figure S17

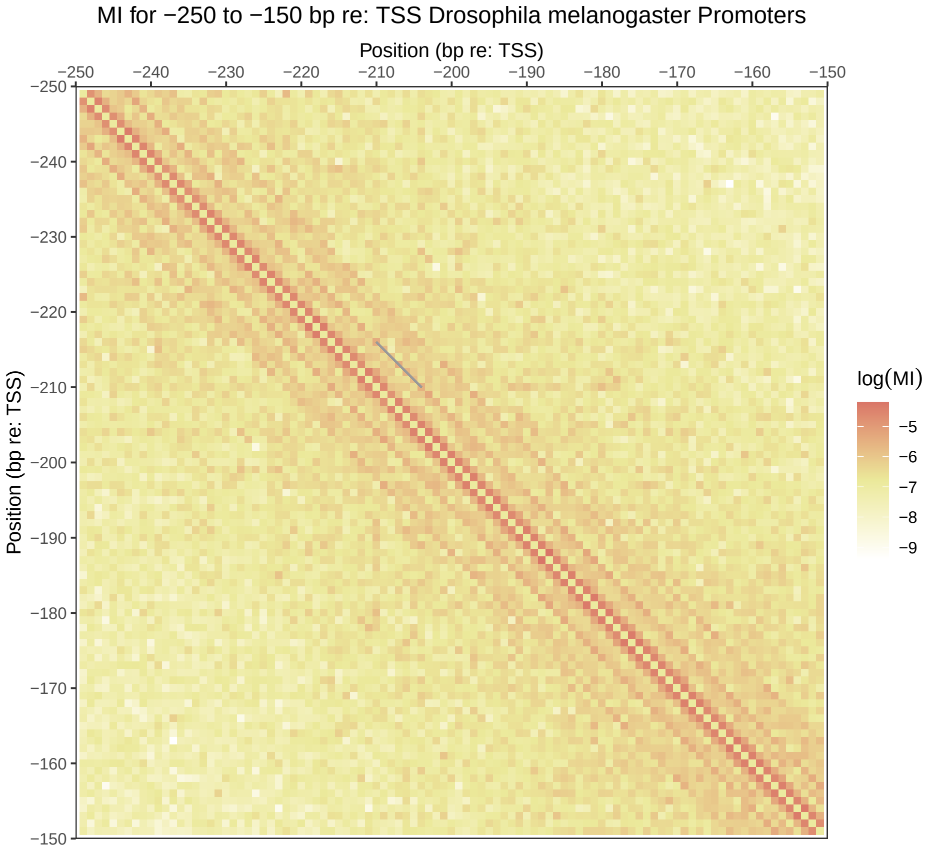
MI for −250 to −150 bp re: TSS Drosophila melanogaster Promoters
Position (bp re: TSS)
−250
−240
−230
−220
−210
−200
−190
−180
−170
−160
−150
Position (bp re: TSS)
log(MI)
−5
−6
−7
−8
−9

Figure S18

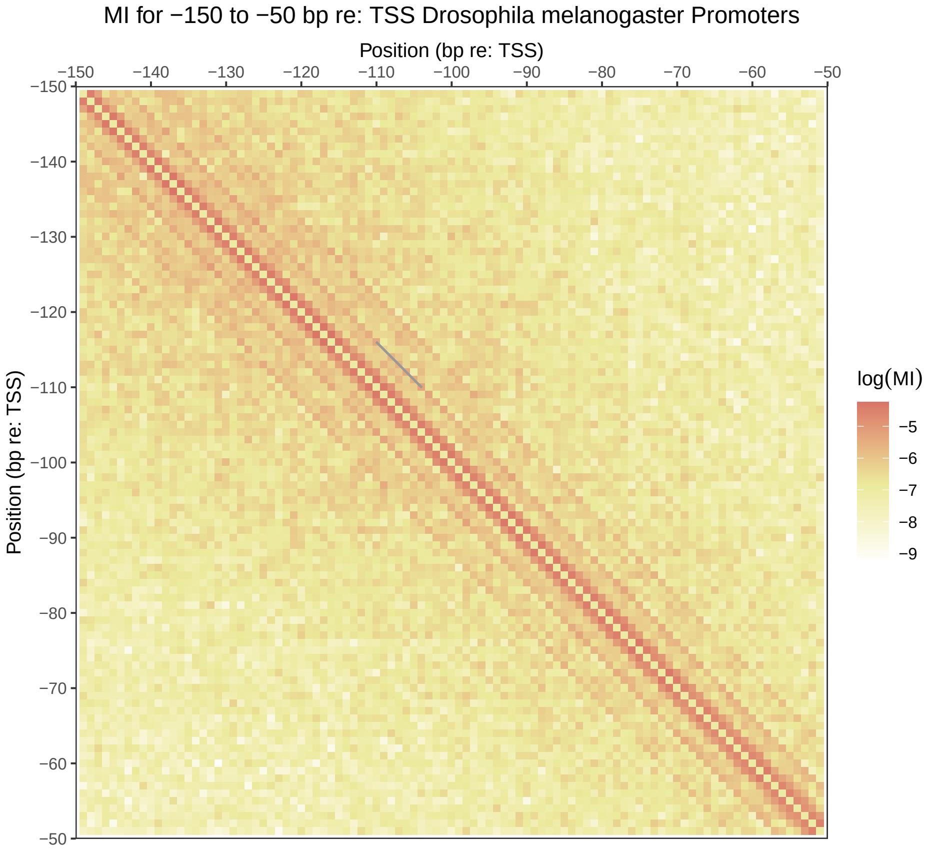
MI for −150 to −50 bp re: TSS Drosophila melanogaster Promoters
Position (bp re: TSS)
−150
−140
−130
−120
−110
−100
−90
−80
−70
−60
−50
Position (bp re: TSS)
log(MI)
−5
−6
−7
−8
−9

Figure S19

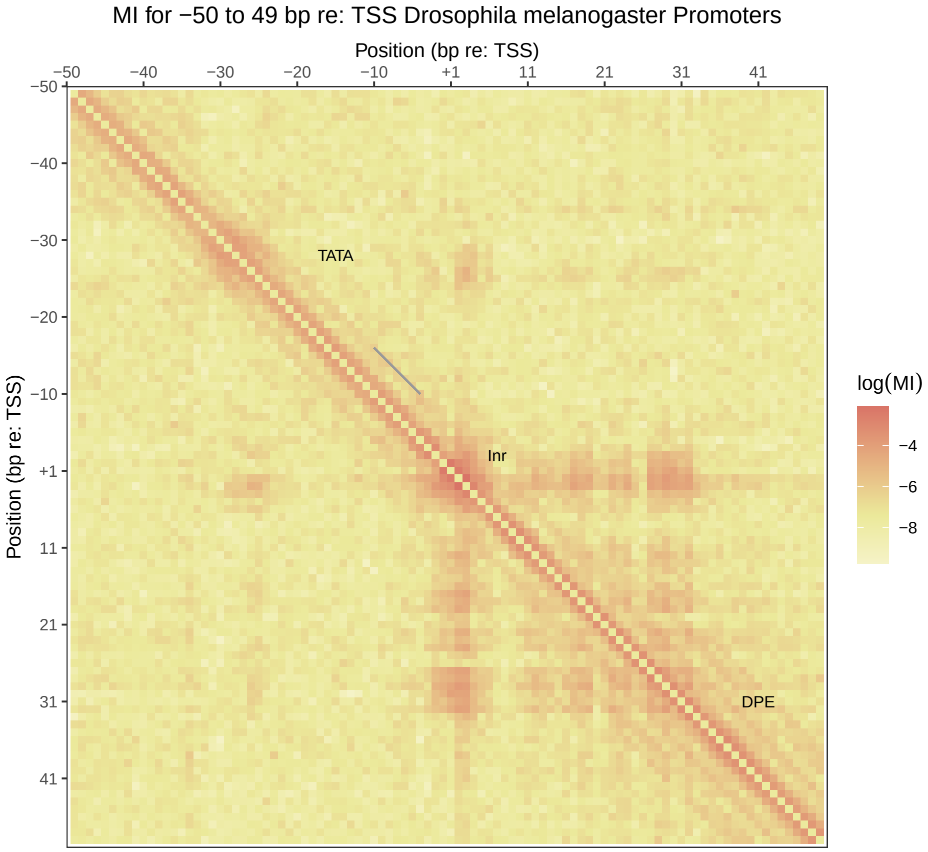

MI for −50 to 49 bp re: TSS Drosophila melanogaster Promoters
Position (bp re: TSS)
−50
−40
−30
−20
−10
+1
11
21
31
41
Position (bp re: TSS)
TATA
Inr
DPE
log(MI)
−4
−6
−8


Figure S20

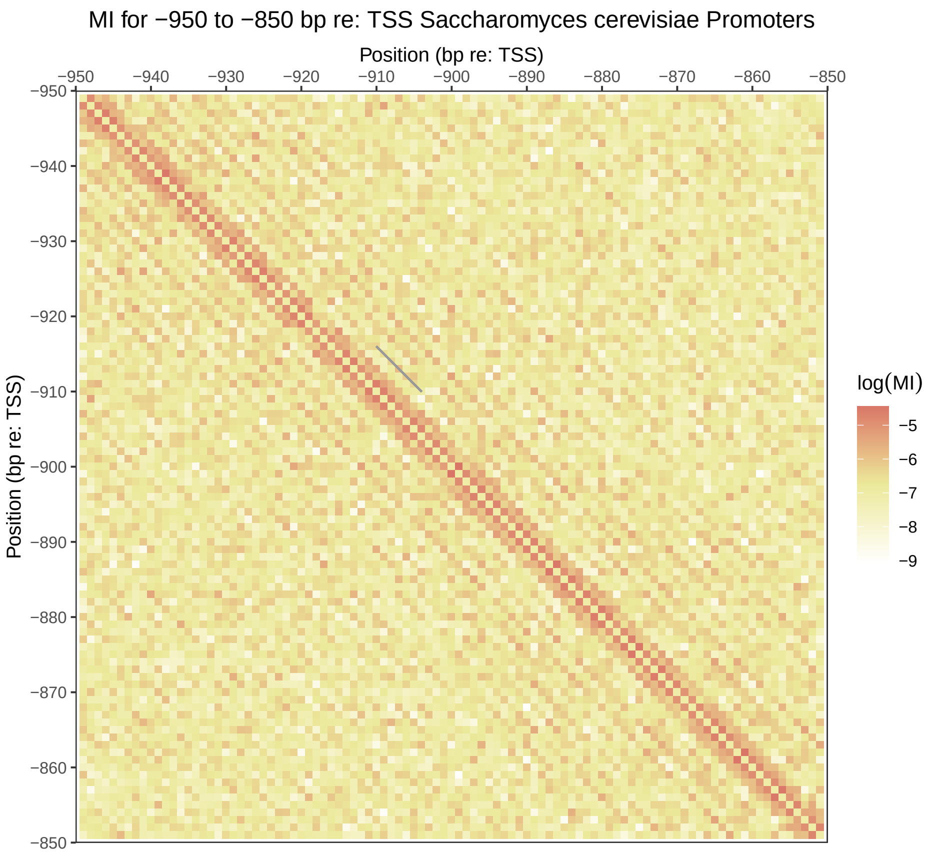
MI for −950 to −850 bp re: TSS Saccharomyces cerevisiae Promoters
Position (bp re: TSS)
−950
−940
−930
−920
−910
−900
−890
−880
−870
−860
−850
Position (bp re: TSS)
log(MI)
−5
−6
−7
−8
−9

Figure S21

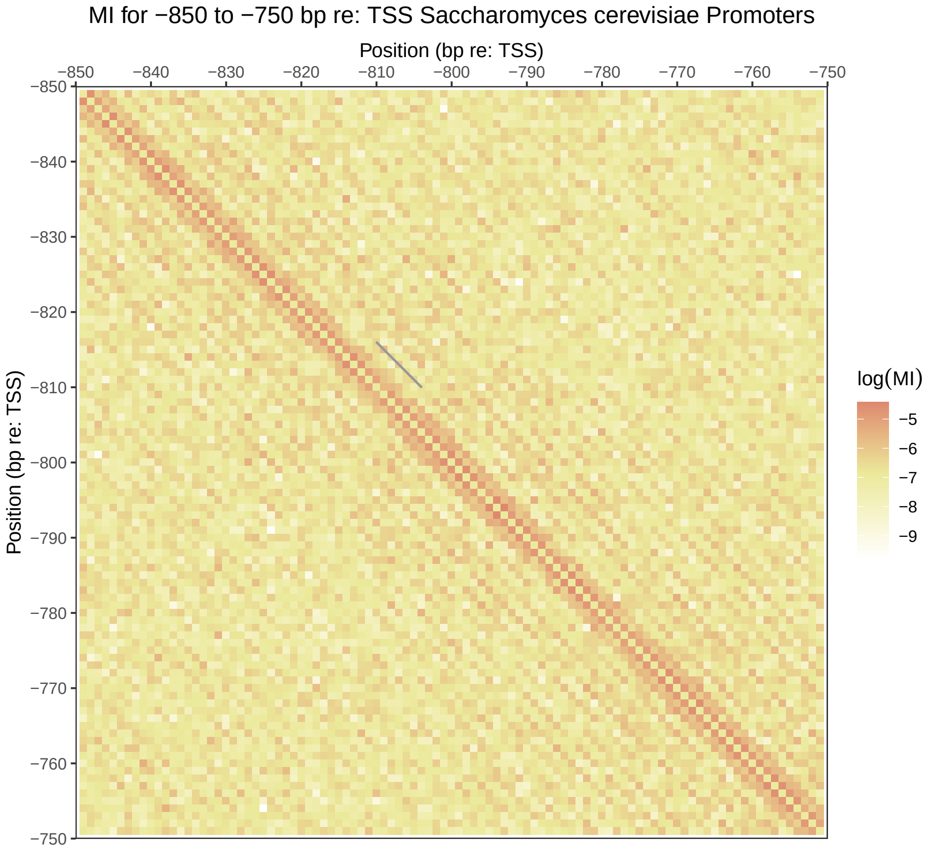
MI for −850 to −750 bp re: TSS Saccharomyces cerevisiae Promoters
Position (bp re: TSS)
−850
−840
−830
−820
−810
−800
−790
−780
−770
−760
−750
Position (bp re: TSS)
log(MI)
−5
−6
−7
−8
−9

Figure S22

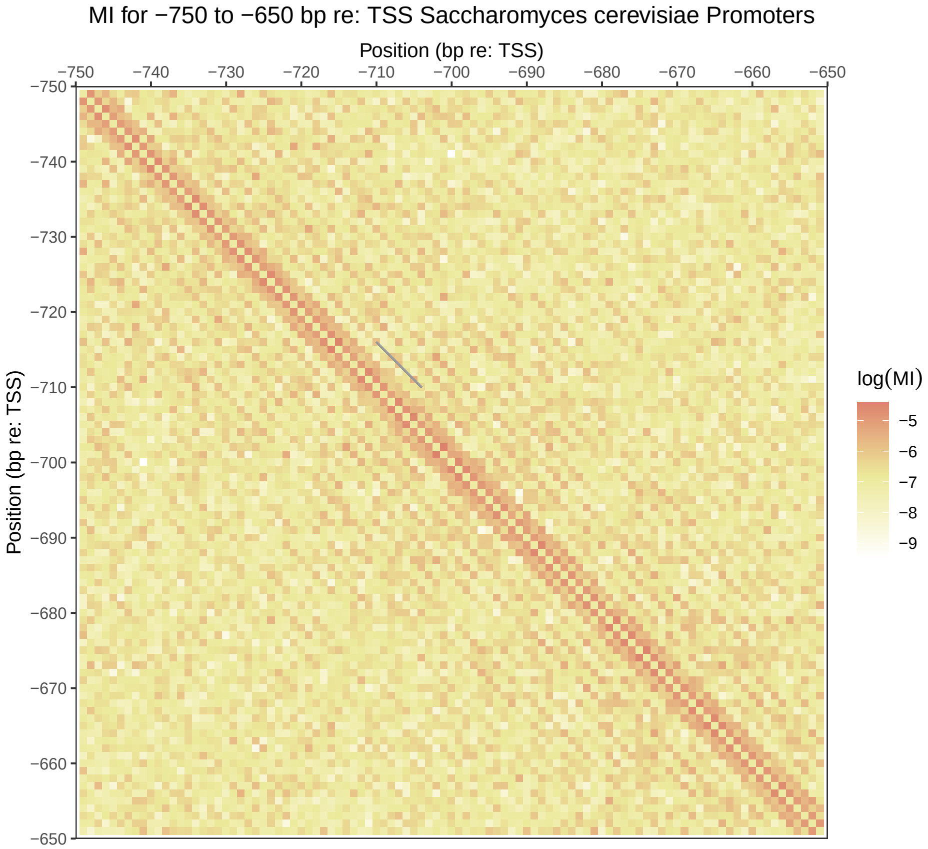

MI for −750 to −650 bp re: TSS Saccharomyces cerevisiae Promoters
Position (bp re: TSS)
−750
−740
−730
−720
−710
−700
−690
−680
−670
−660
−650
Position (bp re: TSS)
log(MI)
−5
−6
−7
−8
−9


Figure S23

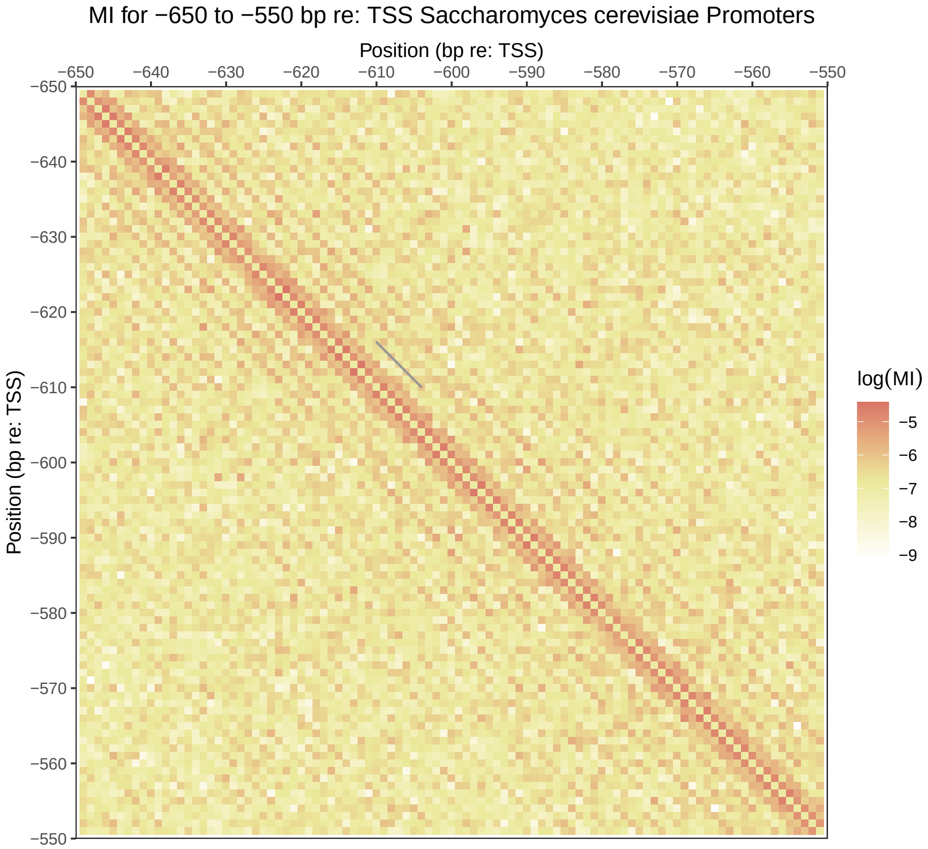
MI for −650 to −550 bp re: TSS Saccharomyces cerevisiae Promoters
Position (bp re: TSS)
−650
−640
−630
−620
−610
−600
−590
−580
−570
−560
−550
Position (bp re: TSS)
log(MI)
−5
−6
−7
−8
−9

Figure S24

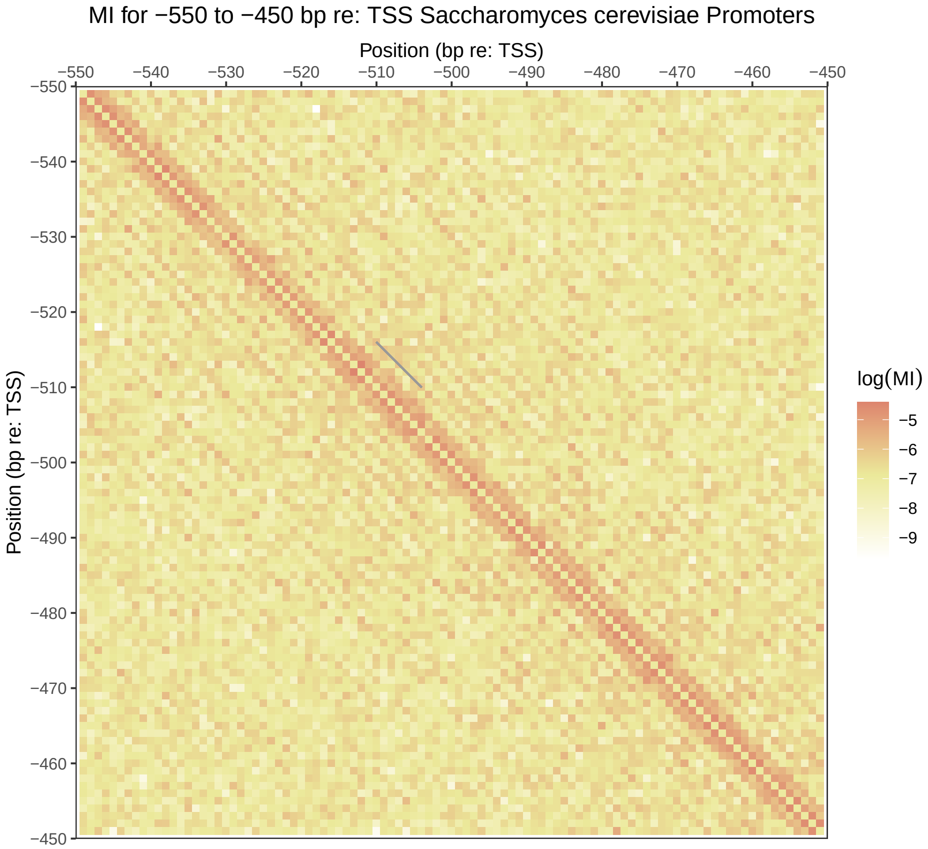
MI for −550 to −450 bp re: TSS Saccharomyces cerevisiae Promoters
Position (bp re: TSS)
−550
−540
−530
−520
−510
−500
−490
−480
−470
−460
−450
Position (bp re: TSS)
log(MI)
−5
−6
−7
−8
−9

Figure S25

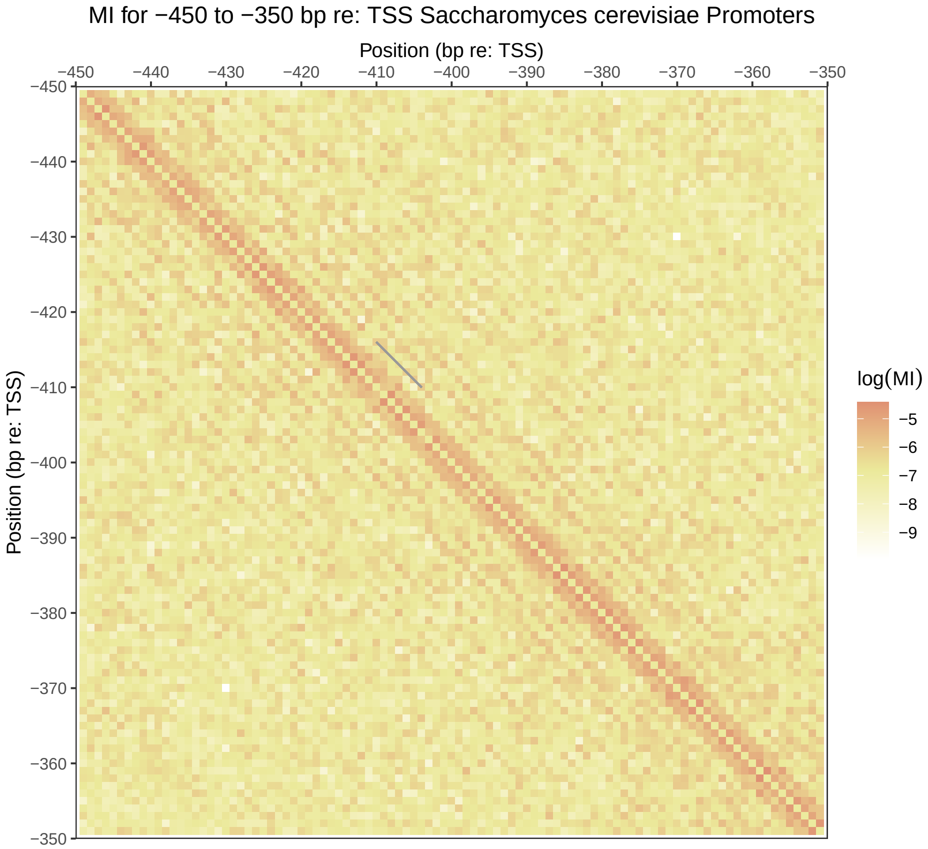
MI for −450 to −350 bp re: TSS Saccharomyces cerevisiae Promoters
Position (bp re: TSS)
−450
−440
−430
−420
−410
−400
−390
−380
−370
−360
−350
Position (bp re: TSS)
log(MI)
−5
−6
−7
−8
−9

Figure S26

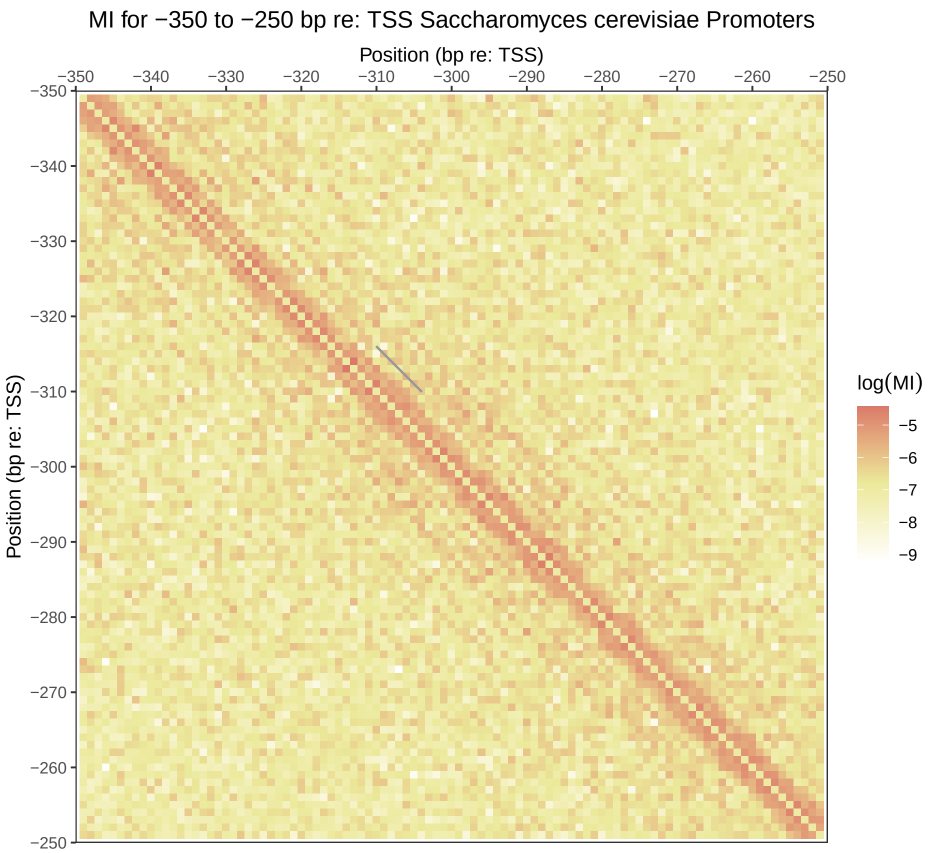
MI for −350 to −250 bp re: TSS Saccharomyces cerevisiae Promoters
Position (bp re: TSS)
−350
−340
−330
−320
−310
−300
−290
−280
−270
−260
−250
Position (bp re: TSS)
log(MI)
−5
−6
−7
−8
−9

Figure S27

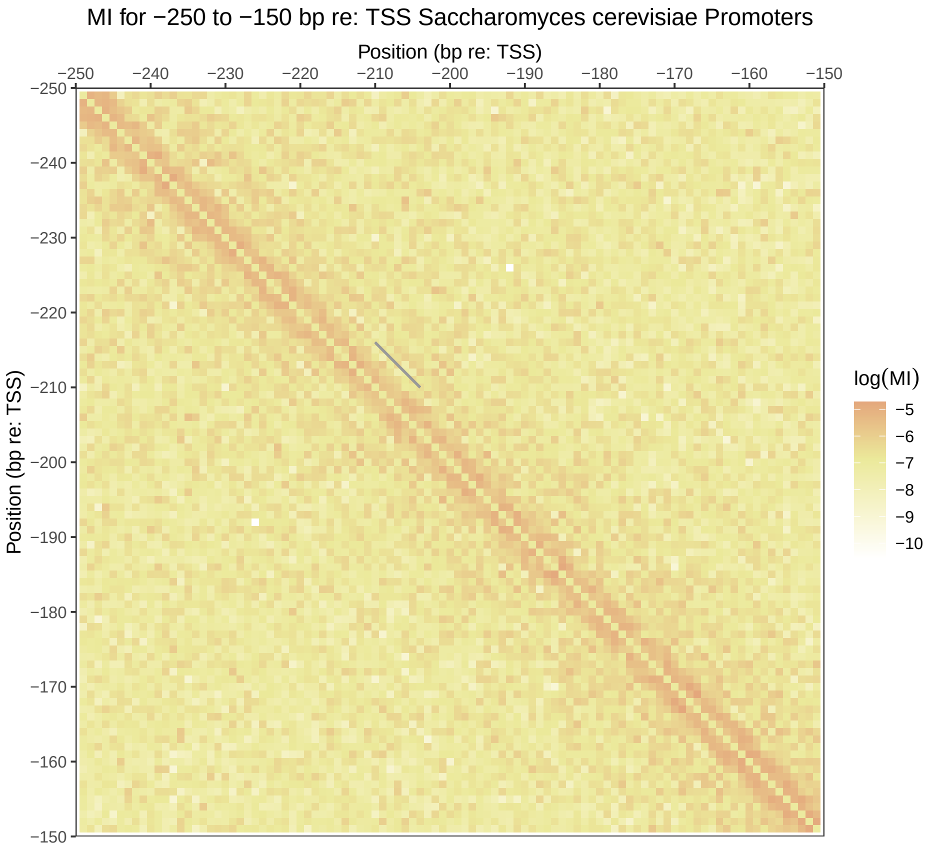
MI for −250 to −150 bp re: TSS Saccharomyces cerevisiae Promoters
Position (bp re: TSS)
−250
−240
−230
−220
−210
−200
−190
−180
−170
−160
−150
Position (bp re: TSS)
log(MI)
−5
−6
−7
−8
−9
−10

Figure S28

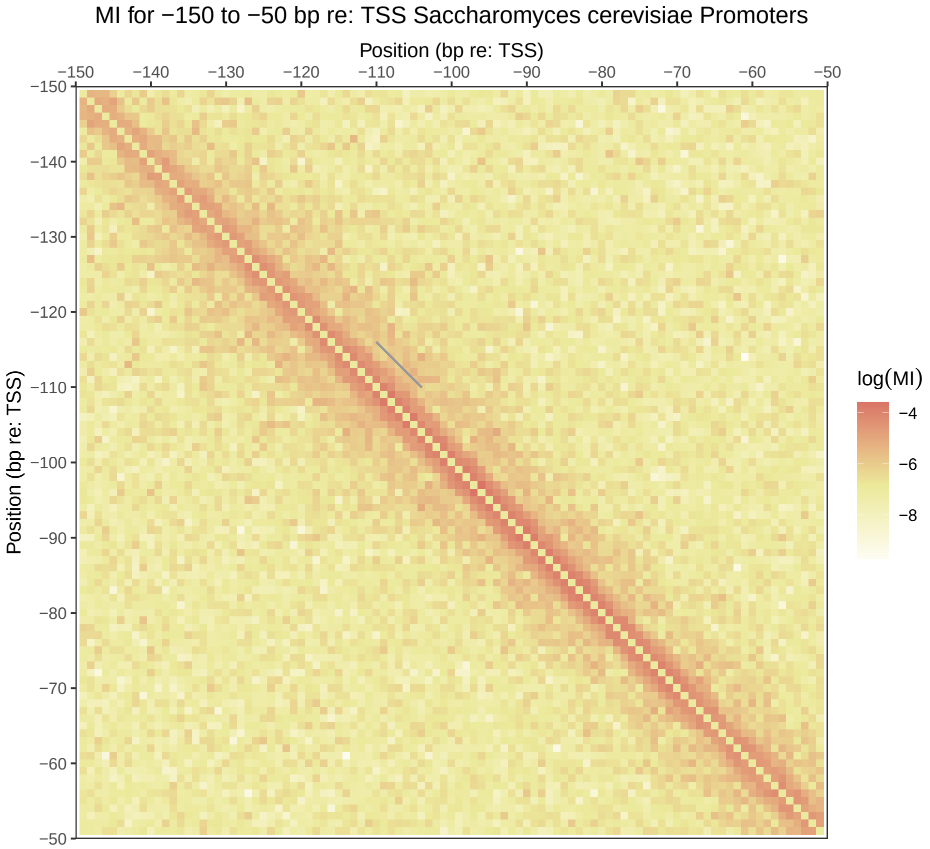
MI for −150 to −50 bp re: TSS Saccharomyces cerevisiae Promoters
Position (bp re: TSS)
−150
−140
−130
−120
−110
−100
−90
−80
−70
−60
−50
Position (bp re: TSS)
log(MI)
−4
−6
−8

Figure S29

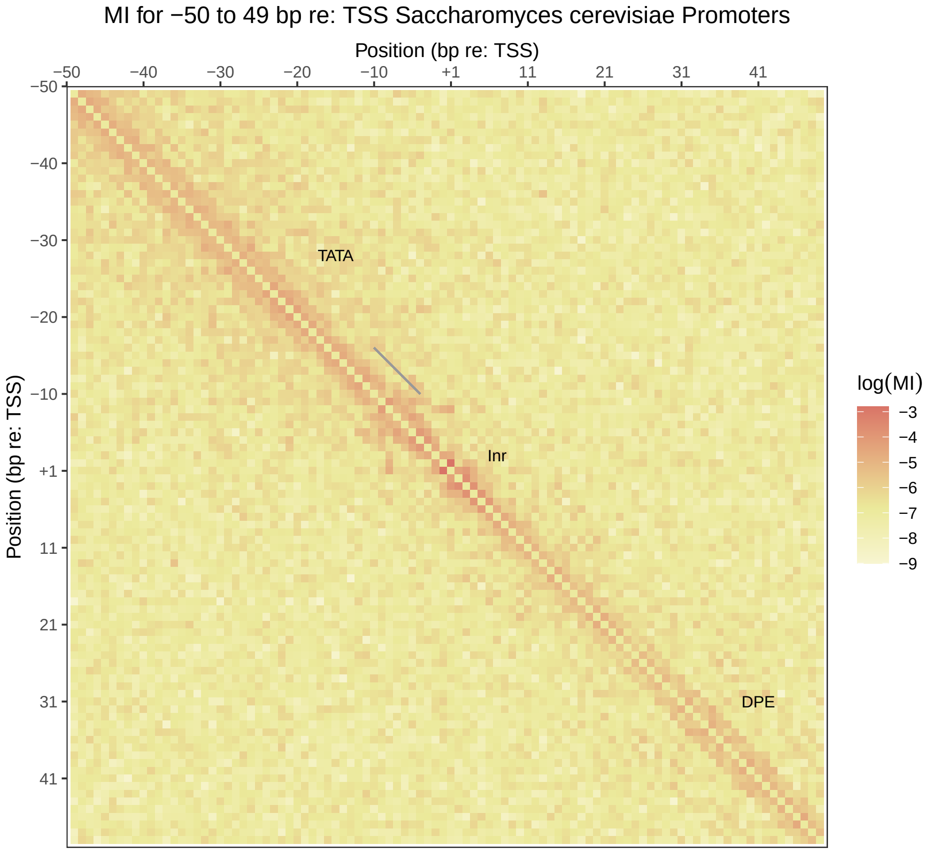
MI for −50 to 49 bp re: TSS Saccharomyces cerevisiae Promoters
Position (bp re: TSS)
−50
−40
−30
−20
−10
+1
11
21
31
41
Position (bp re: TSS)
TATA
Inr
DPE
log(MI)
−3
−4
−5
−6
−7
−8
−9

Figure S30

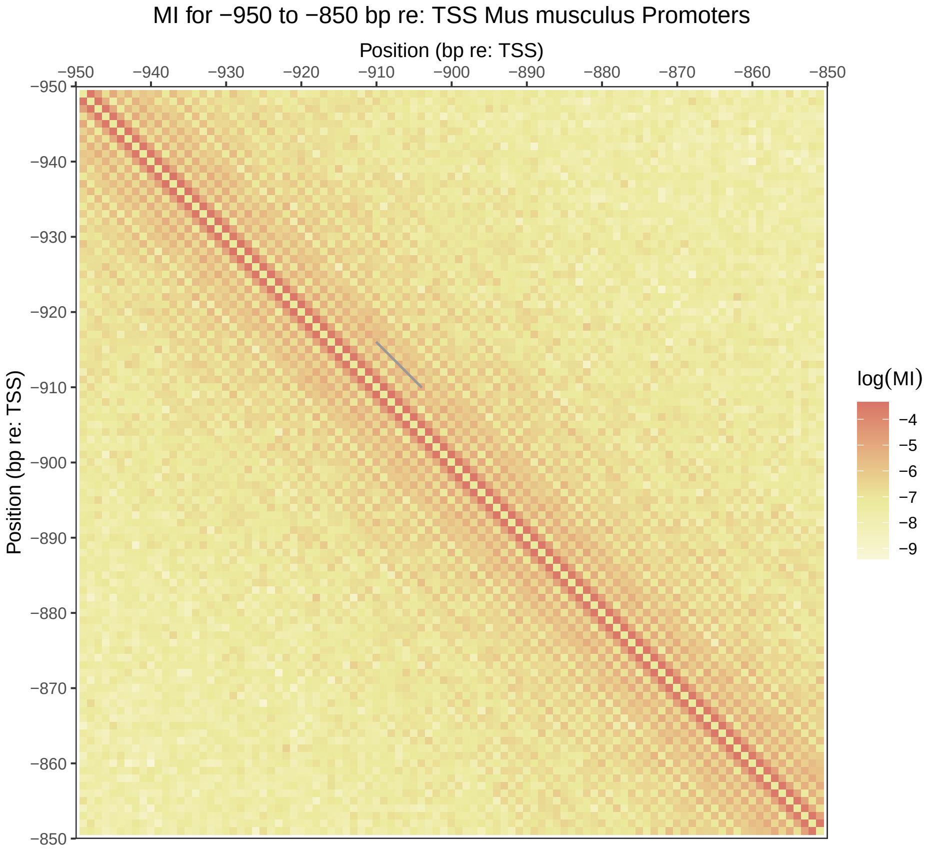
MI for −950 to −850 bp re: TSS Mus musculus Promoters
Position (bp re: TSS)
−950
−940
−930
−920
−910
−900
−890
−880
−870
−860
−850
Position (bp re: TSS)
log(MI)
−4
−5
−6
−7
−8
−9

Figure S31

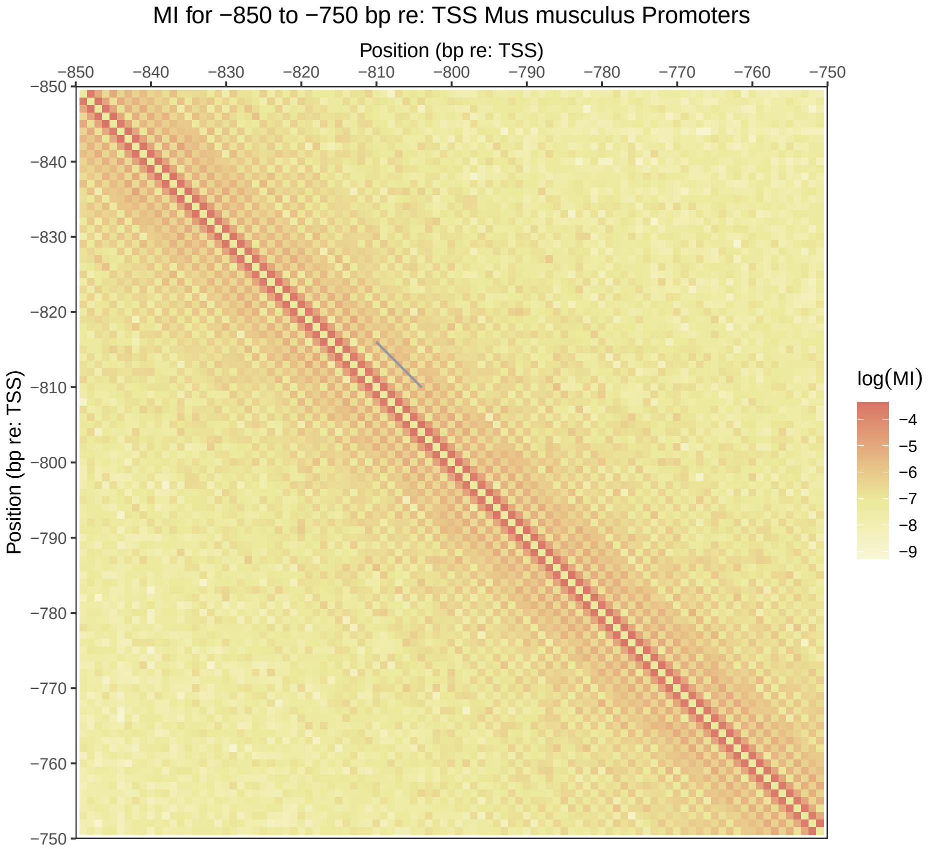
MI for −850 to −750 bp re: TSS Mus musculus Promoters
Position (bp re: TSS)
−850
−840
−830
−820
−810
−800
−790
−780
−770
−760
−750
Position (bp re: TSS)
log(MI)
−4
−5
−6
−7
−8
−9

Figure S32

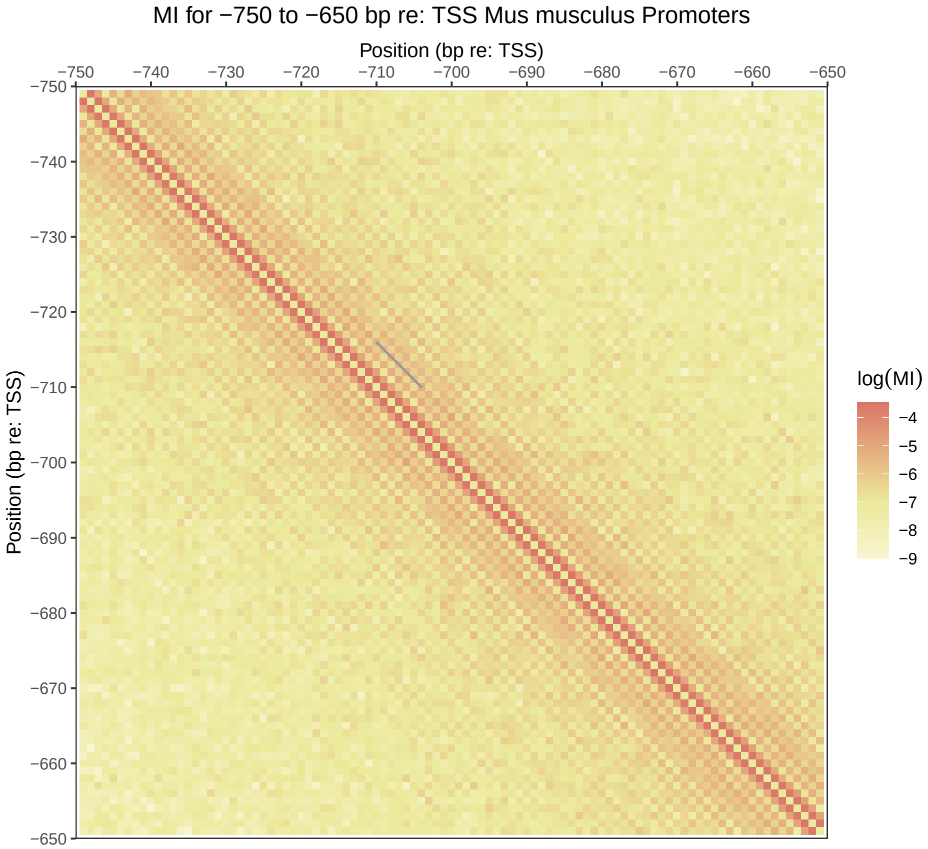
MI for −750 to −650 bp re: TSS Mus musculus Promoters
Position (bp re: TSS)
−750
−740
−730
−720
−710
−700
−690
−680
−670
−660
−650
Position (bp re: TSS)
log(MI)
−4
−5
−6
−7
−8
−9

Figure S33

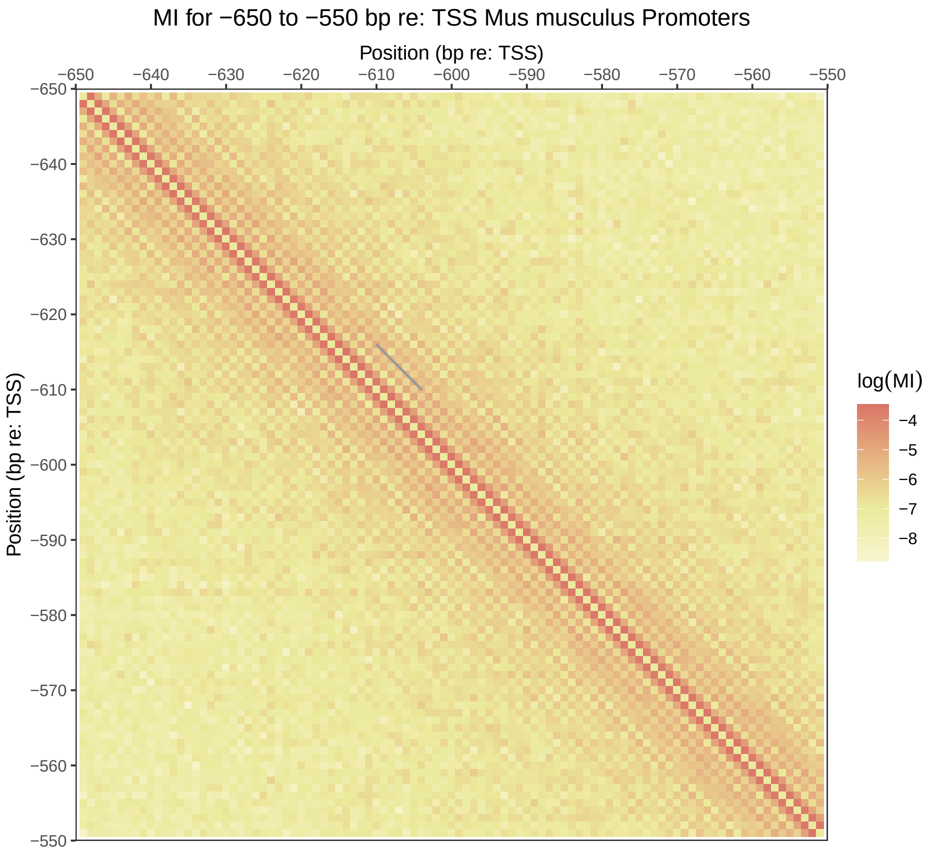
MI for −650 to −550 bp re: TSS Mus musculus Promoters
Position (bp re: TSS)
−650
−640
−630
−620
−610
−600
−590
−580
−570
−560
−550
Position (bp re: TSS)
log(MI)
−4
−5
−6
−7
−8

Figure S34

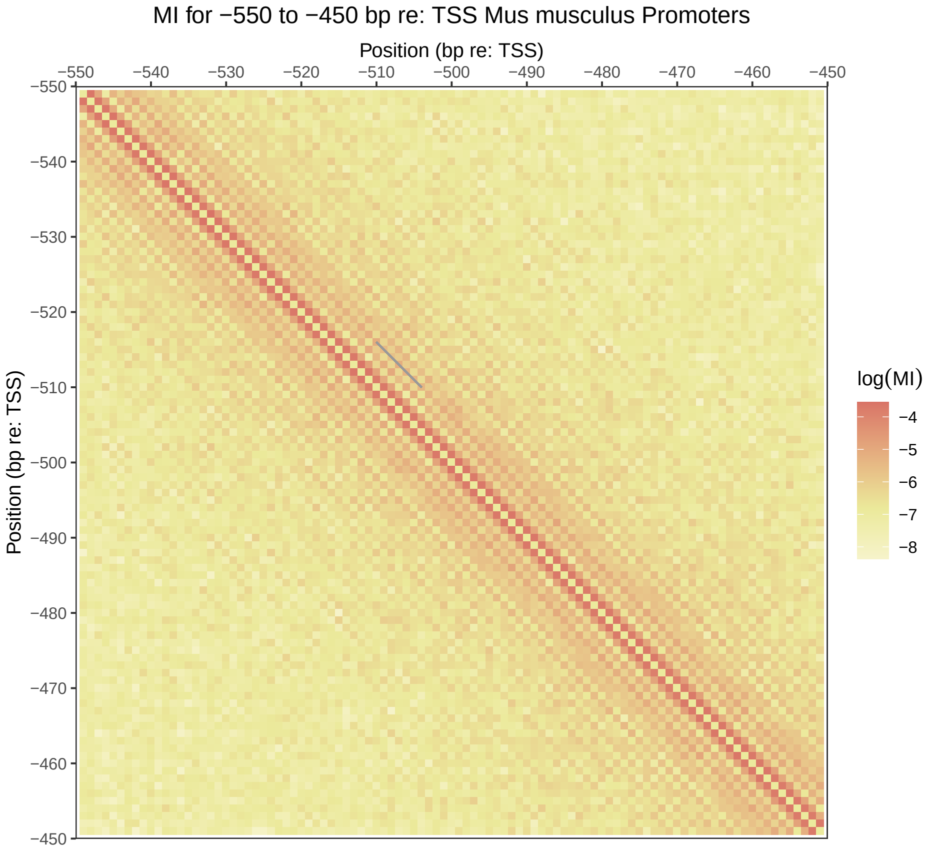
MI for −550 to −450 bp re: TSS Mus musculus Promoters
Position (bp re: TSS)
−550
−540
−530
−520
−510
−500
−490
−480
−470
−460
−450
Position (bp re: TSS)
log(MI)
−4
−5
−6
−7
−8

Figure S35

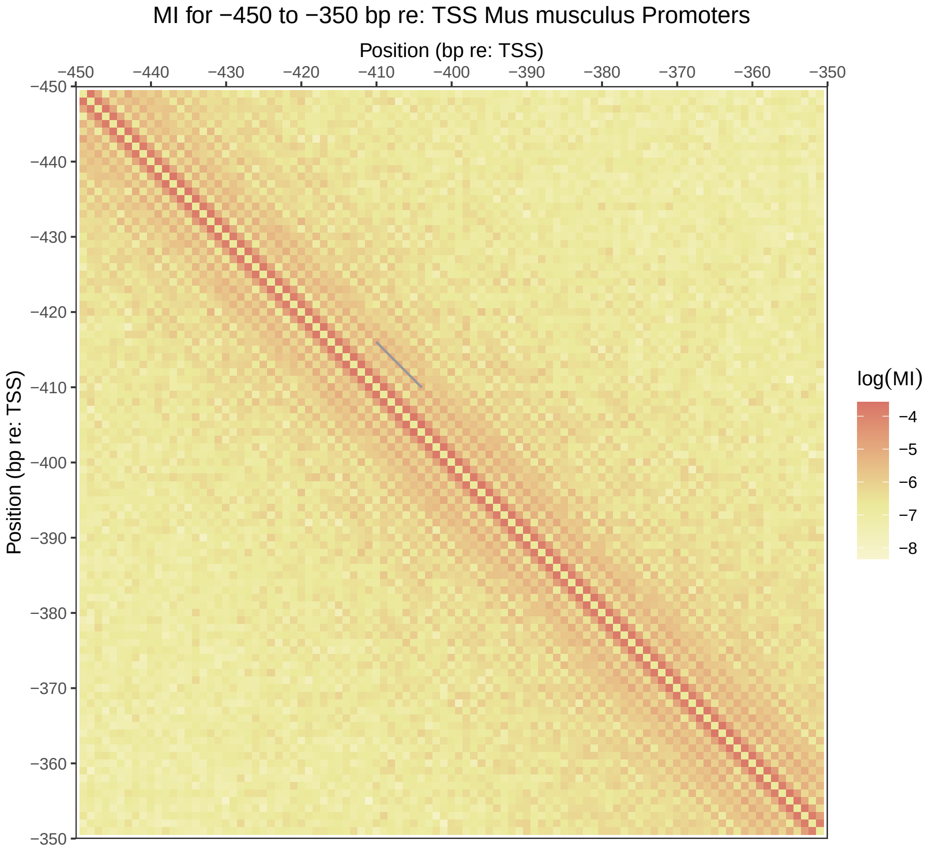
MI for −450 to −350 bp re: TSS Mus musculus Promoters
Position (bp re: TSS)
−450
−440
−430
−420
−410
−400
−390
−380
−370
−360
−350
Position (bp re: TSS)
log(MI)
−4
−5
−6
−7
−8

Figure S36

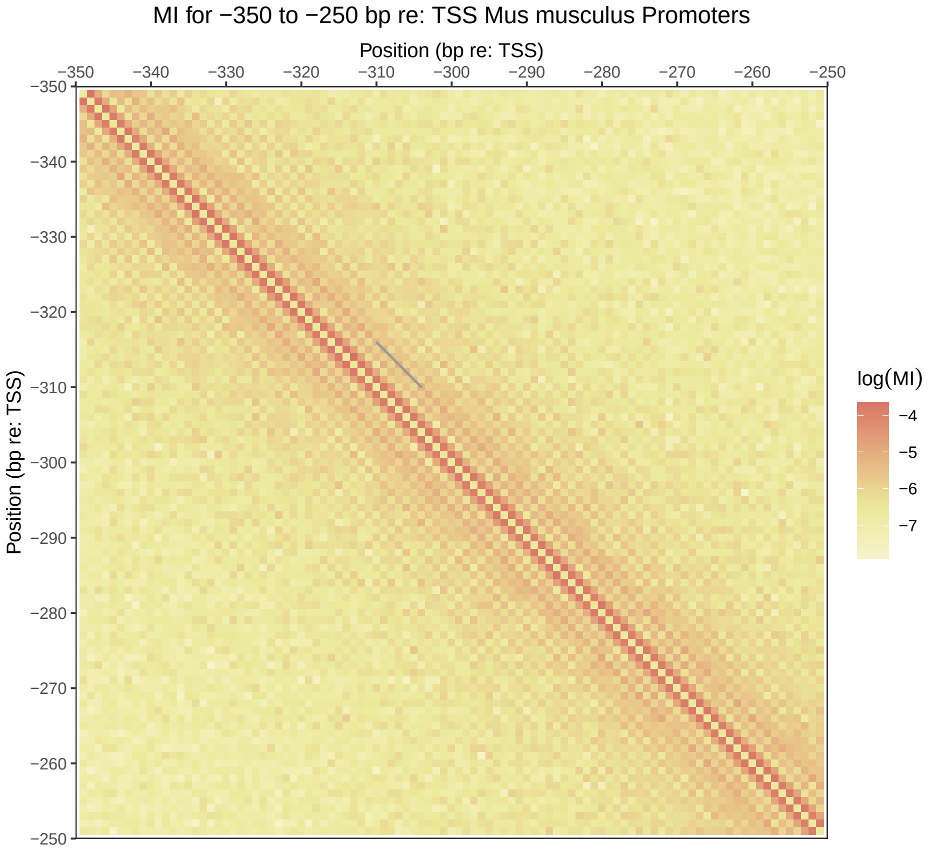
MI for −350 to −250 bp re: TSS Mus musculus Promoters
Position (bp re: TSS)
−350
−340
−330
−320
−310
−300
−290
−280
−270
−260
−250
Position (bp re: TSS)
log(MI)
−4
−5
−6
−7

Figure S37

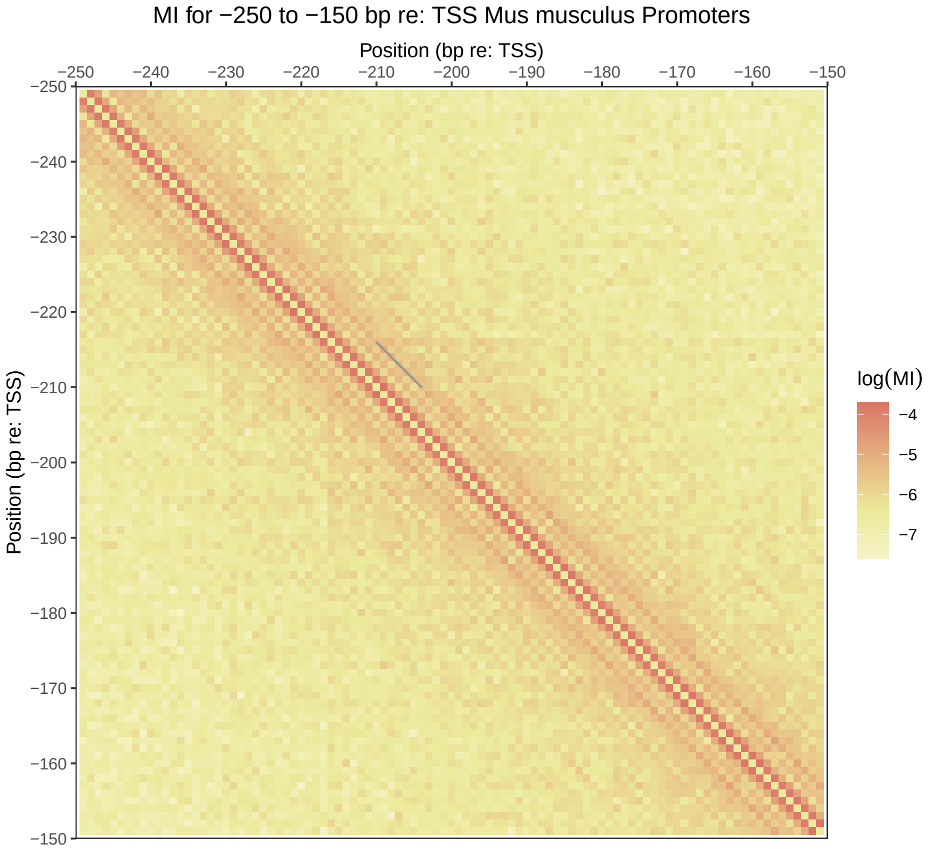
MI for −250 to −150 bp re: TSS Mus musculus Promoters
Position (bp re: TSS)
−250
−240
−230
−220
−210
−200
−190
−180
−170
−160
−150
Position (bp re: TSS)
log(MI)
−4
−5
−6
−7

Figure S38

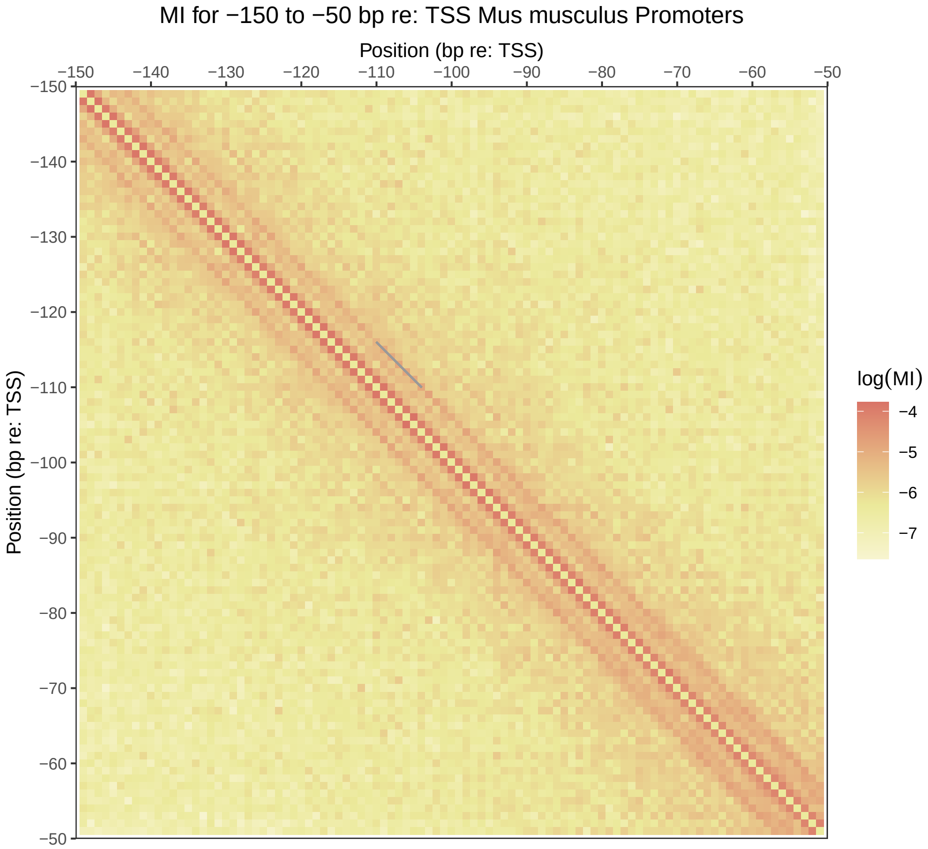
MI for −150 to −50 bp re: TSS Mus musculus Promoters
Position (bp re: TSS)
−150
−140
−130
−120
−110
−100
−90
−80
−70
−60
−50
Position (bp re: TSS)
log(MI)
−4
−5
−6
−7

Figure S39

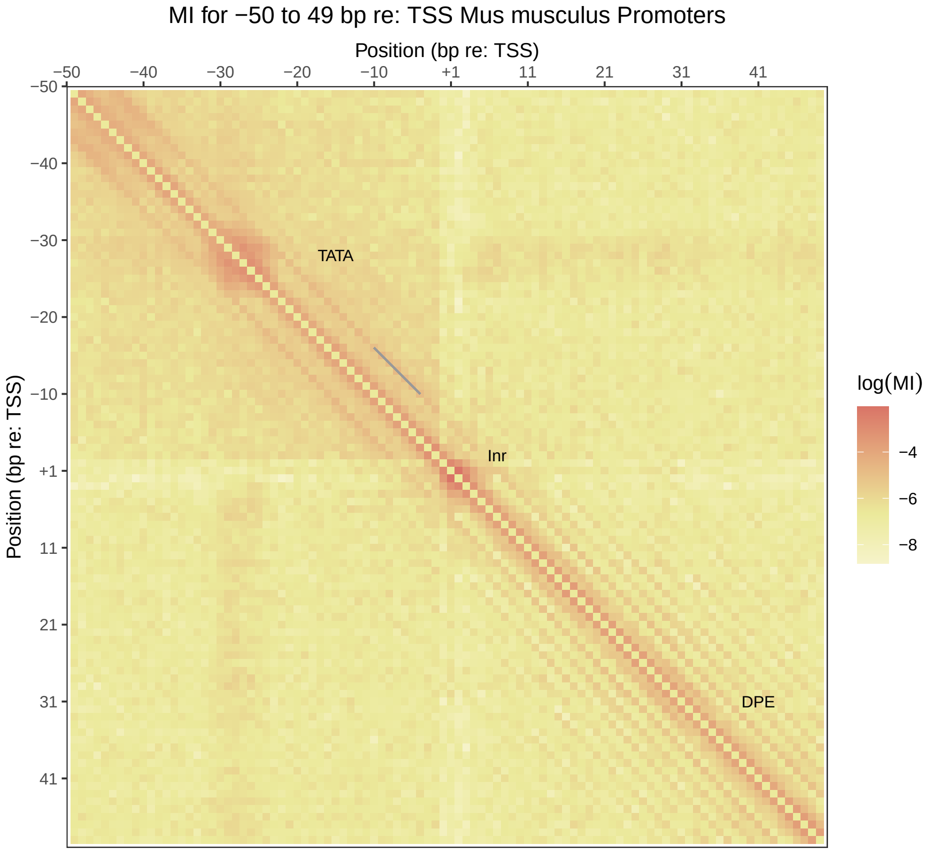
MI for −50 to 49 bp re: TSS Mus musculus Promoters
Position (bp re: TSS)
−50
−40
−30
−20
−10
+1
11
21
31
41
Position (bp re: TSS)
TATA
Inr
DPE
log(MI)
−4
−6
−8

Figure S40

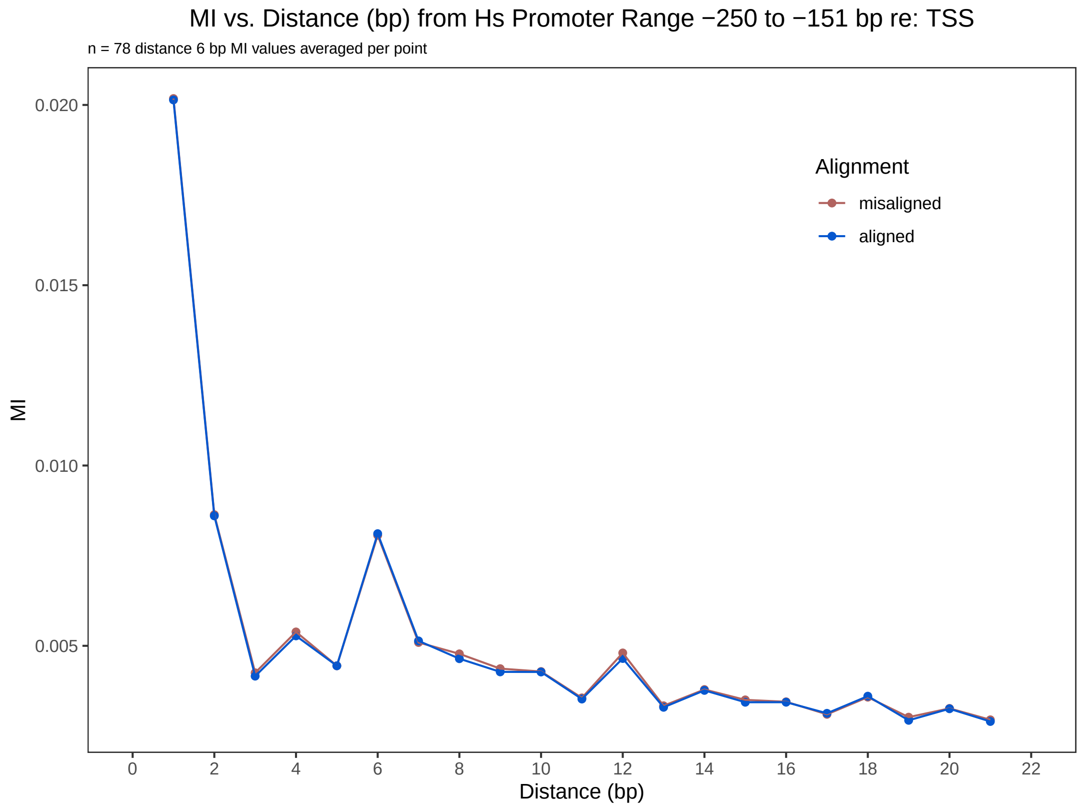


Figure S41: Average MI vs. distance for Homo sapiens promoters from -250 to -151 bp re: TSS. Each of the misaligned sequences was misaligned by a random integer in the range from -40 to +40 bp.

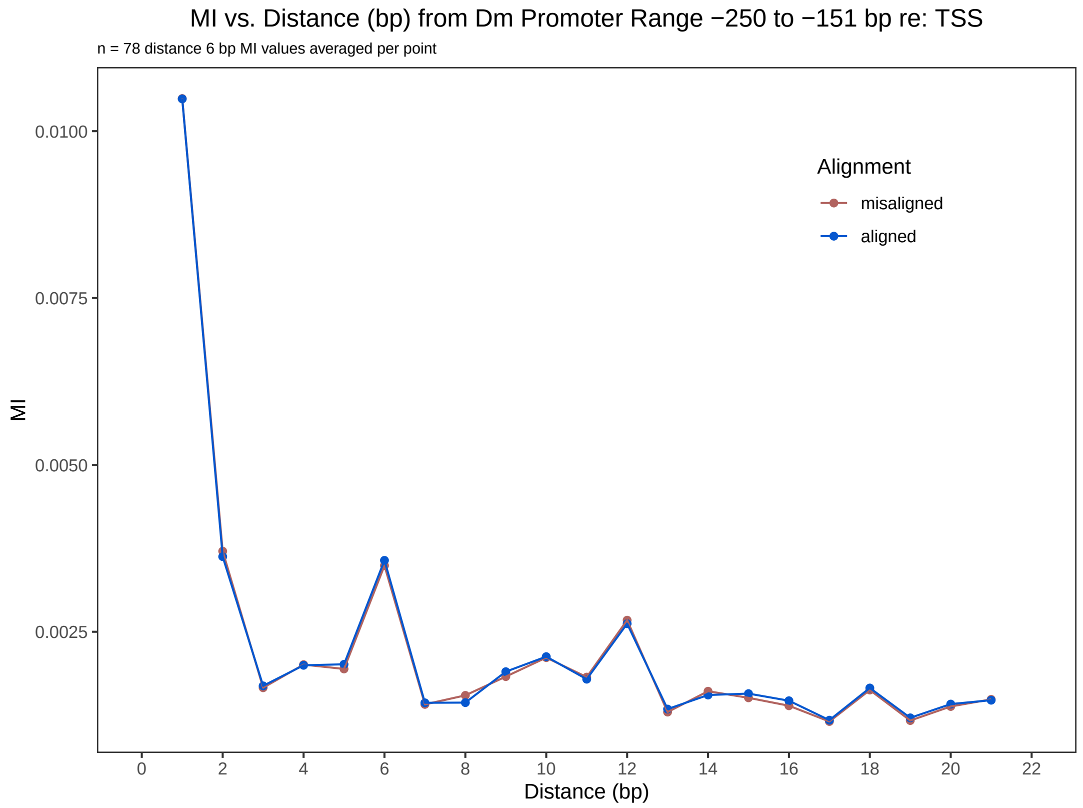


Figure S42: Average MI vs. distance for Drosophila melanogaster promoters from -250 to -151 bp re: TSS. Each of the misaligned sequences was misaligned by a random integer in the range from -40 to +40 bp.

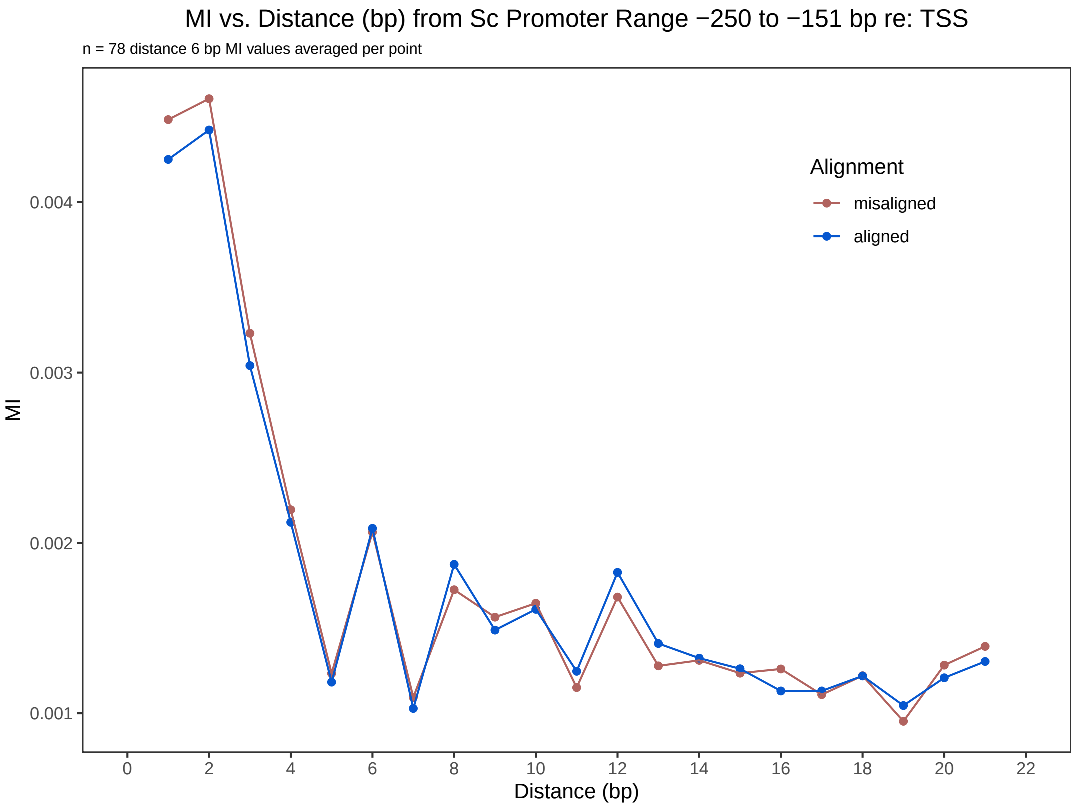


Figure S43: Average MI vs. distance for Saccharomyces cerevisiae promoters from -250 to -151 bp re: TSS. Each of the misaligned sequences was misaligned by a random integer in the range from -40 to +40 bp.

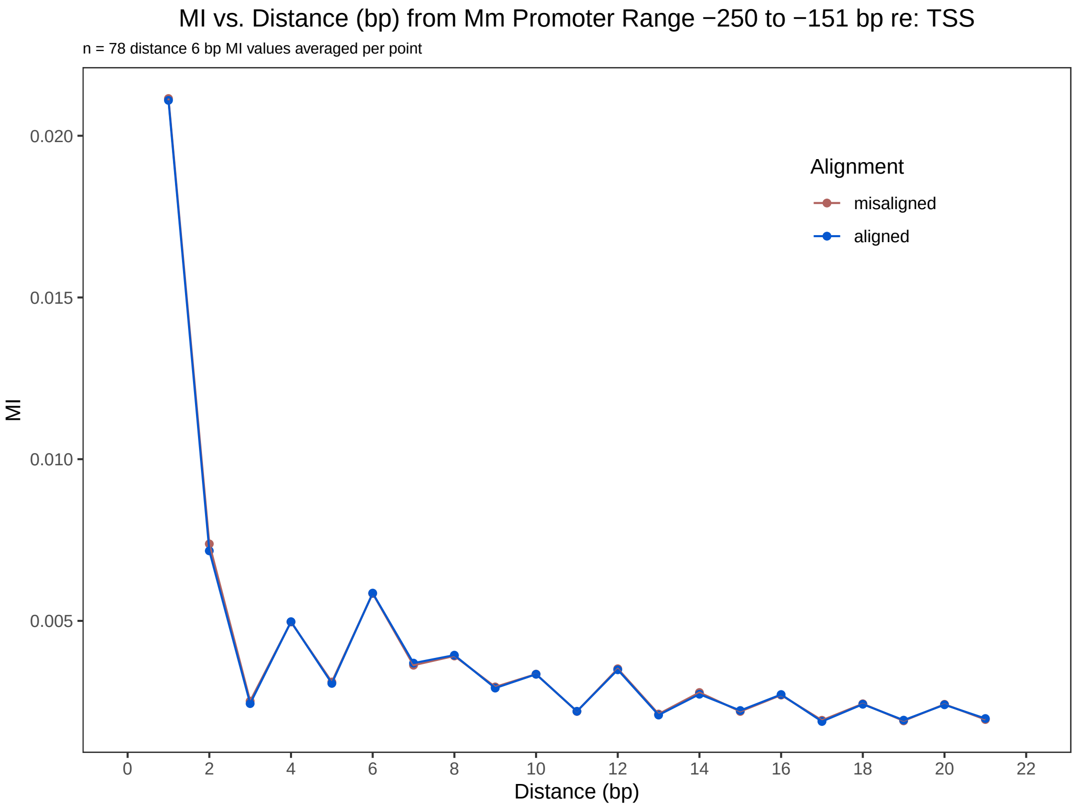


Figure S44: Average MI vs. distance for Mus musculus promoters from -250 to -151 bp re: TSS. Each of the misaligned sequences was misaligned by a random integer in the range from -40 to +40 bp.

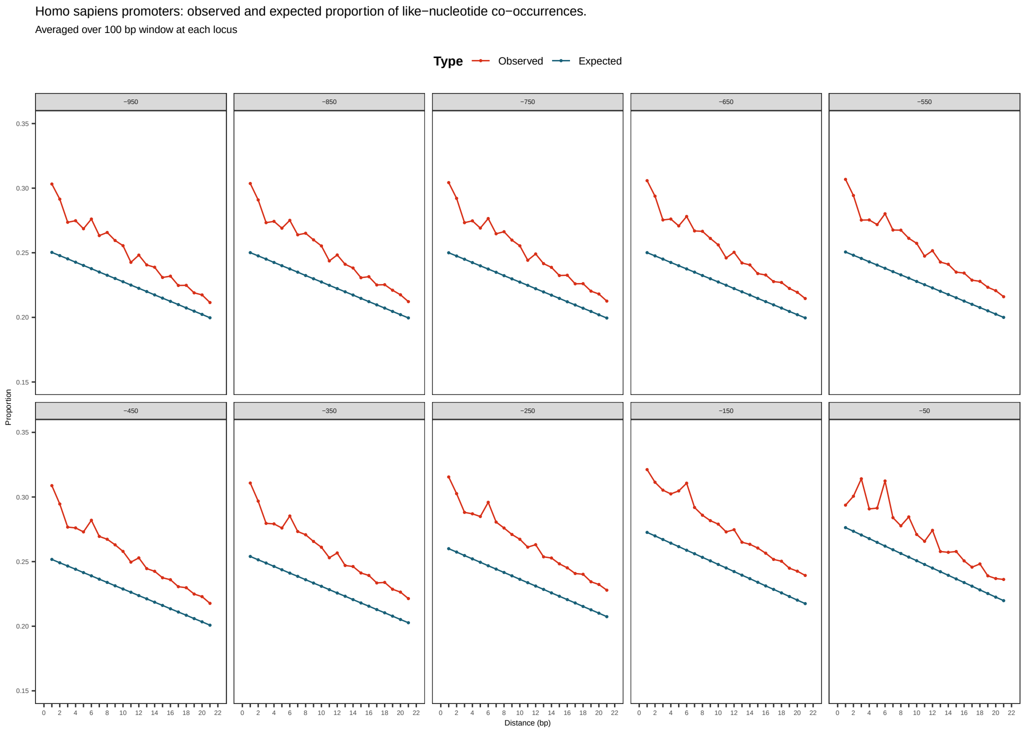


Figure S45: Observed and expected proportions of like – nucleotides at distances from 1 to 21 bp at 10 different loci over the range from -950 to -50 bp re: TSS. Proportions were computed averaging over 100 bp windows. Please note that the y – axis does not start at 0.

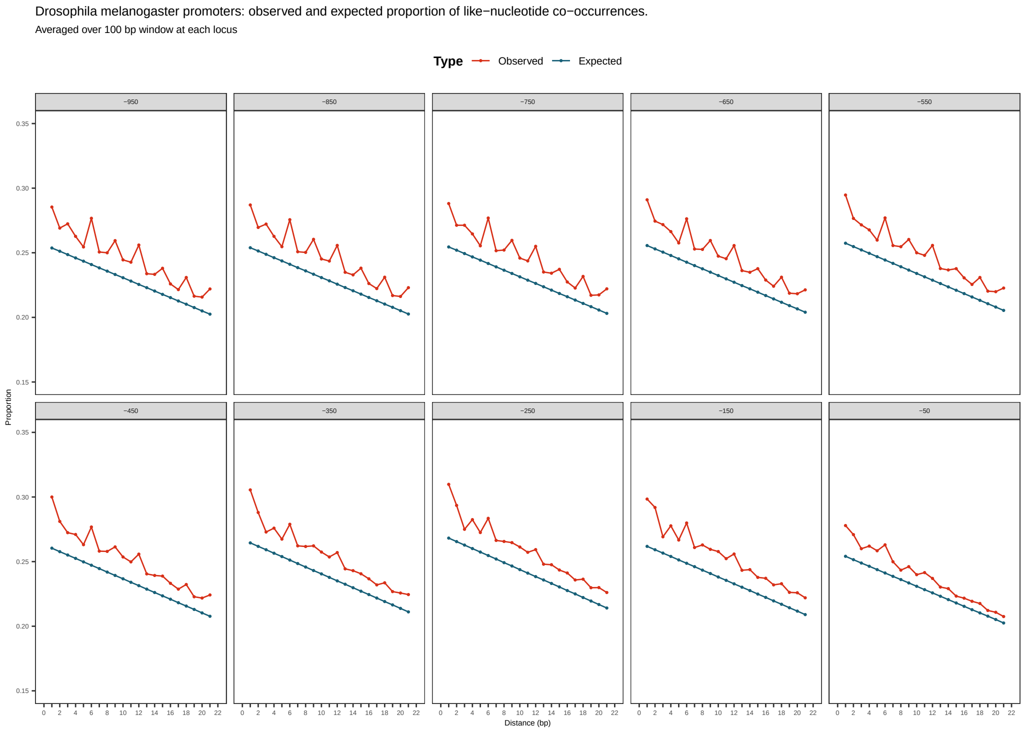


Figure S46: Observed and expected proportions of like – nucleotides at distances from 1 to 21 bp at 10 different loci over the range from -950 to -50 bp re: TSS. Proportions were computed averaging over 100 bp windows. Please note that the y – axis does not start at 0.

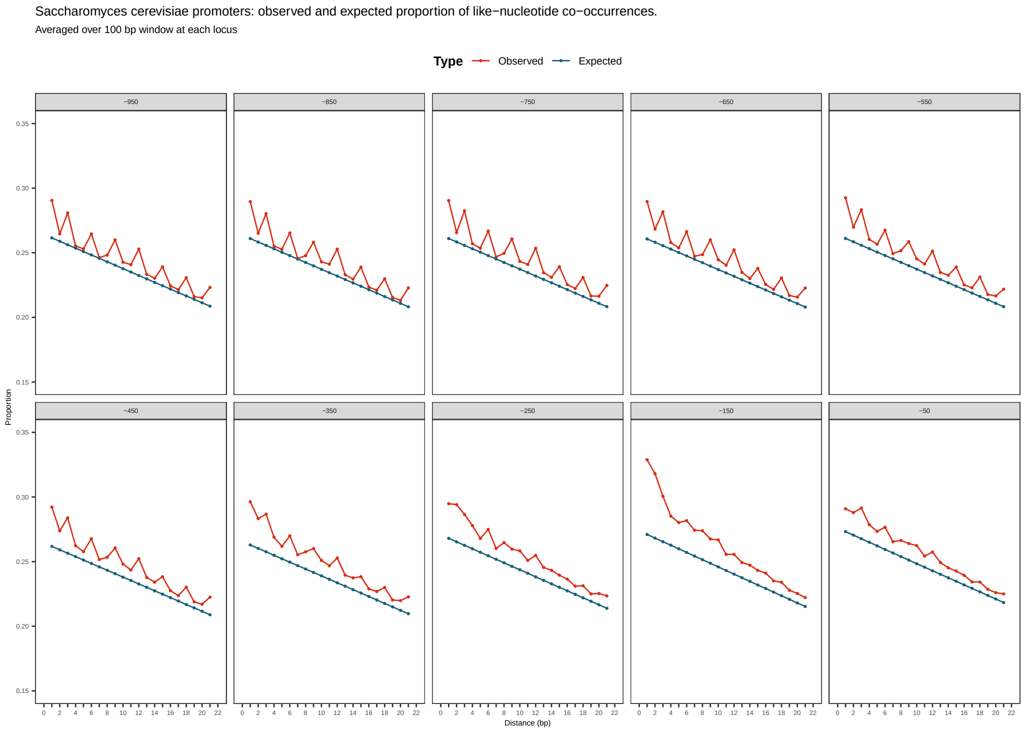


Figure S47: Observed and expected proportions of like – nucleotides at distances from 1 to 21 bp at 10 different loci over the range from -950 to -50 bp re: TSS. Proportions were computed averaging over 100 bp windows. Please note that the y – axis does not start at 0.

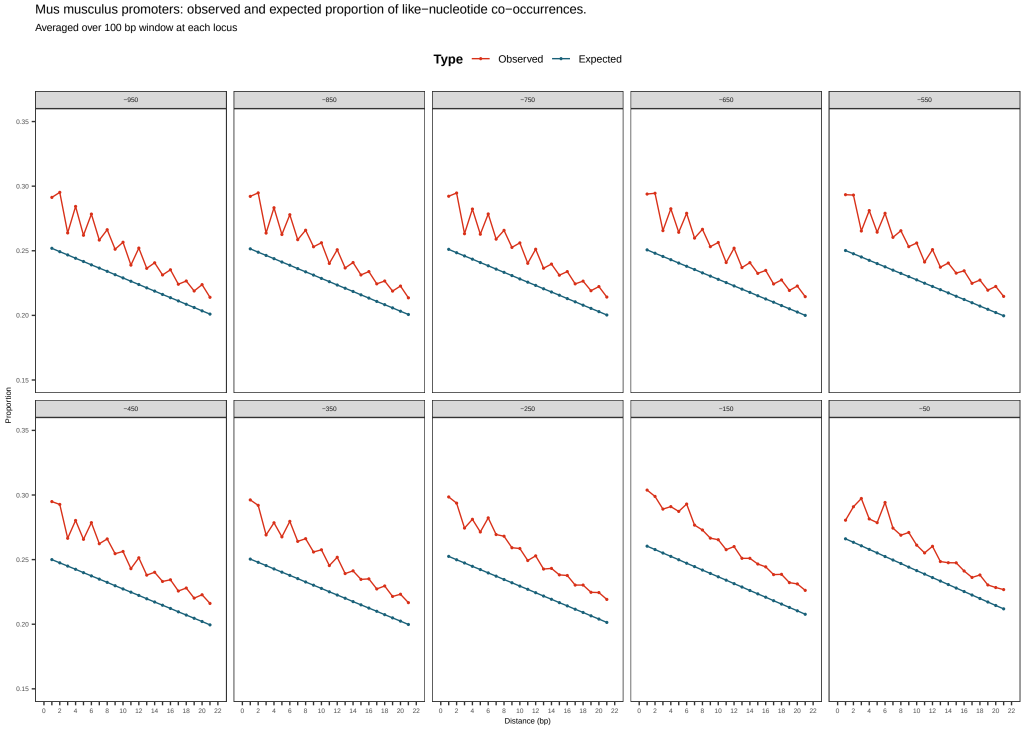


Figure S48: Observed and expected proportions of like – nucleotides at distances from 1 to 21 bp at 10 different loci over the range from -950 to -50 bp re: TSS. Proportions were computed averaging over 100 bp windows. Please note that the y – axis does not start at 0.
